# Radiant DIA: A Fast, Sensitive, and Accurate Search Engine for Quantitative Proteomics

**DOI:** 10.64898/2026.04.29.721743

**Authors:** Seth Just, Lee S. Cantrell, Andrew Nichols, Jian Wang, Janos Kis, Iman Mohtashemi, Theodore Platt, Omid Farokhzad, Serafim Batzoglou

## Abstract

In mass spectrometry-based proteomics, robust and efficient search engines are essential for accurate peptide and protein identification and quantification. Advances in sample preparation and instrumentation have increased the demand for highly scalable processing tools, with datasets comprising hundreds or thousands of samples in single-cell and population studies. Here we present Radiant DIA, a novel Data-Independent Acquisition search engine which achieves 4x faster processing and 10x lower cloud compute costs for large experiments while ensuring rigorous control of false discovery rate (FDR) and maintaining similar sensitivity, precision, and quantitative accuracy to widely-used tools. The Radiant DIA search engine is paired with a modular pipeline deployable on cloud and desktop environments comprising individual modules for distributed re-scoring, FDR estimation, protein inference and quantification. Unlike traditional monolithic applications, this architecture enables high-performance, cloud-scale analysis without sacrificing local usability. Together, the Radiant DIA and Fulcrum Pipeline tools enhance computational efficiency to facilitate biological discovery in large-scale proteomics, as demonstrated by analyses of real-world experiments up to thousands of MS acquisitions.

## Introduction

Proteomics, the large-scale study of protein composition and dynamics, is a central tool to understanding cellular processes and disease mechanisms.^1–5^ Data-Independent Acquisition (DIA) is a discovery mass spectrometry (MS) technique that allows for sensitive and reproducible protein quantification across complex biological samples.^6–8^ However, the performance and scalability of DIA relies heavily on the performance of the search engines used to analyze the massive datasets generated.^9–13^ Whereas affinity proteomic studies can already scale to hundreds of thousands of samples, DIA proteomics studies have traditionally comprised tens to a few hundred acquisitions, though recent advances in instrumentation and sample preparations have enabled emerging cohort-scale studies comprising thousands of samples.^14,15^ Analysis of single-cell atlases, population proteomics, and clinical cohorts is increasingly limited by computational throughput. Bridging this gap requires search engines and pipelines designed explicitly for population-scale MS data.^14,16^

Traditional search engines for DIA proteomics bundle the entire processing pipeline—from scoring to FDR estimation—into monolithic desktop applications.^9–11^ Although these tools have been foundational in advancing proteomics research, their tightly coupled framework can make integration with modern distributed-data processing frameworks more challenging. Recent efforts have introduced cloud distribution for existing engines for portions of the processing workflow, but these approaches still require global processing steps such as FDR estimation and protein inference to run on a single node, restricting scalability.^17–19^ Because global processing still occurs on a single node, and CPU, memory and storage requirements grow super-linearly with dataset size, processing experiments larger than a few thousand MS acquisitions becomes impractical.^19,20^ Resource limitations hinder the analysis of large datasets and the application of innovative data science techniques.^19^

To address limitations we introduce Radiant DIA™, a novel DIA proteomics search engine, and Fulcrum Pipeline™, a scalable, modular processing pipeline, that together addresses these limitations. To our knowledge, this is the first DIA pipeline designed explicitly for distributed search with cluster-scale post-processing. Radiant DIA is an optimized C++ program focused solely on identification and scoring of peptide-spectrum matches (PSMs) in individual files, enabling efficient rapid parallel processing in a cloud-computing environment. Radiant DIA search results are processed by the Fulcrum Pipeline workflow in a distributed cluster computing environment (Apache Spark). Fulcrum Pipeline modules include scalable versions of key workflow components – re-scoring,^21–24^ FDR estimation,^25–29^ protein inference^30,31^ and quantification,^20,32^ – and supports single-pass (“library-free”) and two-pass (“match between runs” or “MBR”) search workflows.^11^ The modular, plug-in Fulcrum Pipeline architecture enables implementation of new workflows using existing modules or to add new modules. Each module can be distributed across a compute cluster, enabling the processing of extremely large experiments without high memory or storage requirement on any single node.

In this work we benchmark Radiant DIA and Fulcrum Pipeline analysis against a widely employed free-to-use DIA workflow across datasets and instruments, assessing computational throughput, identification sensitivity, FDR control, and quantitative performance. We further demonstrate scalability from desktop-scale analyses to multi-instrument cohorts exceeding 2,500 runs, compare with our previous work in cloud-based parallelization of DIA processing workflows, and illustrate biological utility in PTM-rich tissue and a plasma case-control study.

## Methods

### Radiant DIA Search Engine

Radiant DIA is a peptide-centric DIA search engine^10,33,34^ implemented in C++ which reads `mzML` or Parquet files containing MS data. Peptides to be identified can be provided in a TSV or Parquet file, supporting library input from a wide variety of sources and tools (Supplementary Fig. 16). Search engine configuration is read from a TOML file to enable easy recording and reuse of settings across multiple analyses. Comprehensive output is provided in Parquet format, allowing for efficient processing of results. Importantly, this output file can optionally include decoy PSMs and/or the full set of scoring features, allowing for flexible downstream processing, as well as inspection and verification of results.

Reading files is performed in parallel and memory mapping is used whenever possible as reading large libraries / data files accounts for a substantial portion of total runtime in other algorithms.

The program is optimized to minimize the memory footprint of data to ensure the smallest possible resource requirements while still enabling extremely efficient, parallelized implementations of data-intensive operations that allow full utilization of available compute resources.

The Radiant DIA search algorithm utilizes a diverse set of sensitive scoring features. A subset of these features is utilized to compute a discriminant score by linear discriminant analysis (LDA), giving a single score for each candidate PSM that separates targets and decoys. Subsequently, all features are used to train a neural network (NN) classifier that gives final PSM scores. For more information, see Supplementary Methods and Supplementary Fig. 17a.

### Fulcrum Pipeline Orchestration Engine

Fulcrum Pipeline is a flexible and modular proteomics processing pipeline written in Python that leverages Apache Spark to provide highly scalable execution in environments from a single processor up to clusters of thousands of nodes. Fulcrum Pipeline provides full processing workflows for proteomics data processing starting from raw MS data or pre-computed search results and including some or all: PSM rescoring, estimation of PSM/precursor/peptide FDR, protein inference, protein group scoring, protein group FDR estimation, precursor quantification, precursor quant normalization, protein quantification, and output of precursor and protein results (Supplementary Fig 17b).

All aspects of Fulcrum Pipeline utilize a plug-in architecture, allowing for the straightforward addition of new tools, algorithms, or implementations that can replace one or more step in an existing workflow, or new workflows that can immediately leverage existing modules. Simple, well-defined interfaces are used to describe and communicate data and calculations, allowing a diverse set of algorithm implementations and workflows to interoperate, allowing for flexible reuse of implementations. This flexibility allows existing Fulcrum Pipeline workflows to handle a wide variety of use cases without modification and enables rapid development of new approaches. Installed workflows and modules are automatically discovered and can be easily activated. All configuration is provided via a hierarchical TOML file which permits selecting and configuring the workflow and module configurations within a simple, human-writable format.

In this work we utilize a limited set of implementations within the Fulcrum Pipeline workflows, but multiple implementations are available for most steps, offering various algorithmic, statistical, and performance qualities. These include: PSM rescoring using mokapot^24^ or a distributed Spark implementation; global FDR estimation using TDC or MixMax^26^ algorithms as implemented in `mokapot`^24^ or `crema`,^28,29^ or a novel Spark implementation of MixMax, which we call “blocking MixMax” (described in detail in Supplementary Methods); and protein group quantification by sum, max, or DirectLFQ.^20,32^ A complete description and evaluation of each of these modules is outside the scope of this work. For results presented here, we employ the steps described below. Further details are provided in Supplementary Methods.

#### Search and Scoring

Plugins within the Fulcrum Pipeline framework are used to execute Radiant DIA search steps either locally, running sequentially on all input MS files, or in parallel within a cloud execution environment hosted in a Kubernetes cluster (see Supplementary Methods). Search results from each file are filtered at 50% FDR, including decoys, to allow for downstream estimation of global FDR. Radiant DIA NN classifier scores are used without downstream rescoring.

#### PSM and Precursor FDR Estimation by Blocking MixMax

In proteomics experiments it is important to control the rate of identification errors at multiple levels, namely the PSM, precursor, and protein group (PG) levels. By precursor, we mean a unique peptide sequence, modification state, and charge state. A PSM is a precursor identified in a single file. Fulcrum Pipeline workflows control PSM and precursor error rate by estimating the false discovery rate (FDR) at both PSM and precursor levels. PSM-level error rate control ensures that the proportion of false identifications is below a given threshold. However, because samples across an experiment are largely similar, the set of true-positive precursors will be highly correlated between files, while random false-positive matches will not be, the rate of falsely-identified precursors grows with the number of samples, necessitating control of precursor FDR.

To ensure accurate FDR control with optimal sensitivity while avoiding expensive and long-running *q*-value calculations, we have developed “Blocking MixMax”, an efficient bounded approximation of the MixMax^26^ algorithm, designed to run in an Apache Spark cluster computing environment. We use this implementation in all analyses presented in this manuscript. For more information, see Supplementary Methods.

#### Protein Inference, Scoring, and FDR Estimation

Protein inference is performed using Proffer, an efficient, multi-threaded Python implementation of parsimonious protein inference that employs sparse matrix multiplication to efficiently implement graph manipulation operations. Precursors identified at a configurable threshold (1% in this manuscript’s analyses) are associated with protein sequences from a FASTA file by Preppers, an optimized substring matching implementation written in Rust, using the adaptive radix tree data structure.^35^ Peptides are associated with any protein containing their sequence, regardless of enzymatic cleavage locations. For more information, see Supplementary Methods.

#### Precursor and Protein Group Quantification

In this manuscript’s analyses we employ precursor quantities directly from Radiant DIA results. However, Fulcrum Pipeline workflows are designed to allow plug-in modules for precursor quantification, separately from the search step. We envision future development of plugins that provide transition refinement, interference removal, or targeted quantitative re-extraction from MS files.

Precursor quantities may optionally be normalized. Fulcrum Pipeline normalization modules include median or “median-dense” (using the median intensity of analytes detected in a configurable minimum fraction of files) normalization. These implementations optionally support using a fast bounded approximation provided by Apache Spark to improve performance on extremely large datasets (not utilized in the analyses presented here). Support for plug-in normalization modules is provided to allow alternative normalization techniques. For example, analyses of Seer-generated datasets in this manuscript employ a modified median-dense implementation which operates separately on acquisitions prepared with each distinct nanoparticle well from the same biosample in the Proteograph assay. Equivalent normalization was applied to DIA-NN analyses of these datasets when compared.

Normalized intensities are rolled up to the protein group level. Prior to roll-up intensities may optionally be filtered for proteotypicity, or at a stricter FDR threshold than that used for protein grouping, though this can result in an inability to quantify some protein groups. All analyses in this manuscript employ DirectLFQ^20^ for rollup without additional filtering. DirectLFQ rollup is performed in parallel across protein groups, using Spark for distributed execution. The combination of parallelization in a cluster context and the linear scalability of DirectLFQ enable fast execution of this step, even when processing thousands of files.

#### Match Between Runs

Fulcrum Pipeline workflows include an implementation of two-pass or “match-between-runs” searching. In this workflow, search results are generated for all files by searching with the same predicted library used for single-pass “library-free” search. Global precursor-level FDR estimates are then computed as described above. A library (the “MBR library”) is built using empirical spectra and iRT values taken from the highest-scoring identification for each precursor identified at a configurable FDR threshold (1% in this work), including decoys. All files are searched again using the MBR library, and results from this second-pass search used for global PSM FDR estimation. Protein inference, scoring, and FDR estimation are performed using first-pass precursor scores. All remaining steps proceed as described above.

## Results

To evaluate the performance, sensitivity, and scalability of the combined Radiant DIA and Fulcrum Pipeline workflow, we assembled a diverse set of datasets spanning multiple instruments, gradient lengths (15-90 min), sample types (cell lysate, plasma, eye lens), and acquisition complexities. These datasets include PTM-rich tissues, matrix matched calibration curves, a heterogeneous case-control study, and a 2,561-run multi-instrument cohort. Together, they represent a rigorous test bed for assessing identification sensitivity, error-rate calibration, quantitative stability, and cloud-scale execution. For each dataset, we compared the Radiant DIA and Fulcrum Pipeline to DIA-NN 1.8.1, with both workflows run in both library-free (single-pass) and MBR (two-pass) modes. For each comparison, both modes employed the same predicted library as the starting point for search, as described in Supplementary Methods.

DIA-NN version 1.8.1 was selected as a widely used comparison engine with high-quality figures of merit and broad licensing accessibility. We have chosen to use DIA-NN as a baseline as it also employs a peptide-centric search paradigm, offers high performance and sensitivity, and has previously been incorporated in high-throughput cloud processing pipelines, including the “Scalable DIA” workflow utilized for cloud execution analyses in this work.^18,19^ Several alternative tools exist with various search paradigms, performance and sensitivity characteristics.^11,36–38^ These include alphaDIA, which is also freely-usable, peptide-centric, and supports some degree of parallelization for search steps, but is not fully compatible with the fixed-predictions library strategy employed in this work’s comparisons.^39^

### Throughput and Cost

The Radiant DIA search engine is written to make efficient use of computational resources. Execution times for searching identical `mzML` file(s) with the same library are 5-6x faster than DIA-NN 1.8.1 using equivalent processing resources (Supplementary Fig. 1). Efficient use of compute resource enables Radiant DIA to process studies of increasing biological scale.

For large experiments of dozens to thousands of MS acquisitions, local execution can become impractical due to storage limitations and the time required for processing.^19^ Modern mass spectrometers generate data at rates exceeding 10 GB/hour, and datasets comprising 1,000 files exceed 5 TB. Recent advances in sample preparation, chromatography, and instrumentation enable increased study sample count with improved depth of content alongside larger raw data volume to uncover this burden.^40^

Parallelized cloud pipelines address these challenges. By distributing acquisitions across scalable compute resources, Radiant DIA and Fulcrum Pipeline execution reduce the overall time required for data processing. In practice, these architectures reduce analyses that once required weeks of computation to only a few hours. Building on our prior “Scalable MBR” DIA-NN workflow, cloud execution of Radiant DIA and Fulcrum Pipeline delivers further gains in search speed and compute efficiencies (Figure 1a and b), achieving lower per-file cost and shorter end-to-end runtime across cohorts of various size. Although evaluation of cloud pipeline performance is inherently variable due to configuration and resource availability, Radiant DIA and Fulcrum Pipeline processing shows a reproducible advantage in both speed and cost to exceed the scalability of prior solutions by more than 10-fold.^19^

**Figure 1.**
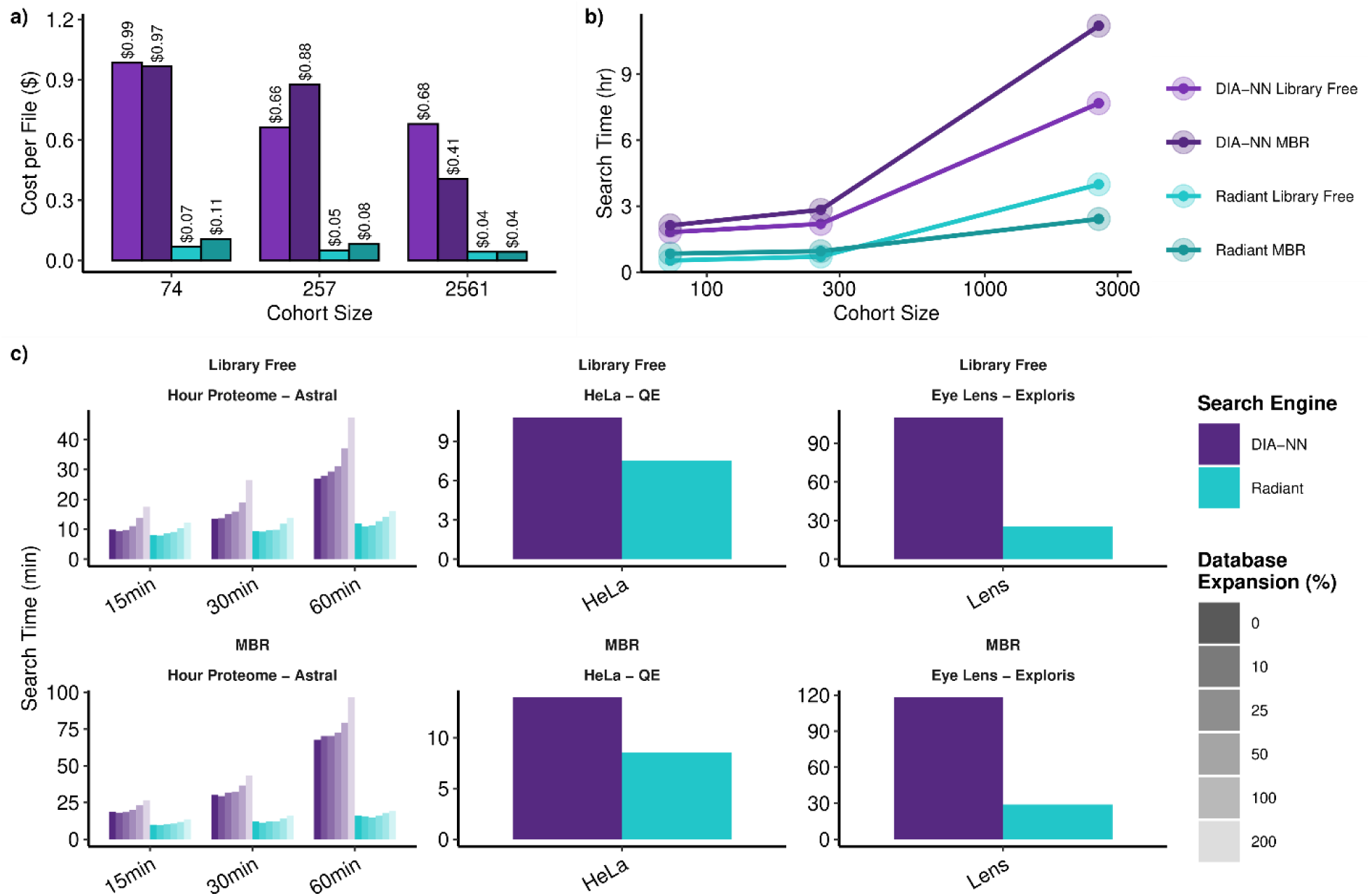
Comparison of parallelized cloud processing pipelines, demonstrating the speed and cost efficiency of the Radiant DIA and Fulcrum Pipeline workflow (teal) versus the “Scalable MBR” DIA-NN cloud pipeline (purple). **A)** Amortized per-file cost for end-to-end cloud processing of cohorts of different size. Cost is inclusive of instance start, file download, and global FDR control, among other steps. **B)** Total time of execution for end-to-end cloud processing of cohorts of different size. Actual performance may vary due to resource availability and pricing at the time of processing. **C)** Benchmarks for Radiant DIA / Fulcrum Pipeline and DIA-NN running in a local execution environment, using a c7a.8xlarge instance with 32 CPUs and c4 GiB of memory.

For smaller experiments, local execution remains attractive. When processing is confined to a single desktop node, runtime is dominated by sequential file analysis. Using a standardized 32-CPU environment we benchmarked three community datasets acquired on different MS instruments (Hour Proteome: Thermo Fisher™ Orbitrap™ Astral™; HeLa: Thermo Fisher

Q-Exactive™ HF; Eye Lens: Thermo Fisher Orbitrap Exploris™ 480). Figure 1c shows that local execution of Radiant DIA and Fulcrum Pipeline tools provide a substantial performance gain over DIA-NN for both library-free and MBR workflows. The minimal runtime penalty for MBR over library free in Fulcrum Pipeline analyses contrasts with the substantial overhead observed for DIA-NN, which results primarily from slower MBR library-generation steps.

Previous efforts to accelerate this step – for example, the `quantms` pipeline – have reduced DIA-NN library-generation time, but do not incorporate empirical evidence that aids MBR performance.^10,19,41^ Fulcrum Pipeline mitigates this by efficiently constructing the MBR library directly from first-pass results, while the speed of Radiant DIA search minimizes cost of second-pass reprocessing. The empirical subset of input spectral library in two-pass search further reduces the computational requirements for downstream processing steps, even as data volume and search space expand, as demonstrated by database- and gradient-extension tests on the Hour Proteome Dataset (Figure 1c).

Finally, Radiant DIA executables can be compiled for ARM processors, enabling acquisition of lower-cost cloud compute. Although runtime of the search step is similar between ARM and x86 architectures, ARM deployment further reduces cost by 19% on AWS (`m7g.2xlarge` vs `m7i.2xlarge`; Supplementary Fig. 1). This enables an additional avenue for cost-efficient cloud analysis or desktop analysis utilizing ARM processors, such as on modern Apple computers, which are not supported by existing software. Together, these results demonstrate that Radiant DIA and Fulcrum Pipeline tools enable practical analysis of datasets that exceed the limits of prior DIA pipelines, both in time-to-results and cost efficiency.

### Identification Sensitivity and Error Rate Control

Search performance is often summarized through per-run peptide spectrum matches (PSM), and resulting precursor, peptide and protein group identifications aggregated across the study. While identification counts are uninformative to quantitative interpretation, they are practical indicators for tool capability.^1,2,8,36,42^ Across all datasets, Radiant DIA results closely match DIA-NN in overall precursor identifications (Figure 2) with modest advantage in the Eye Lens MBR dataset enabled by improved handling of post-translational modification (PTM) rich spectra (further discussed below).

**Figure 2.**
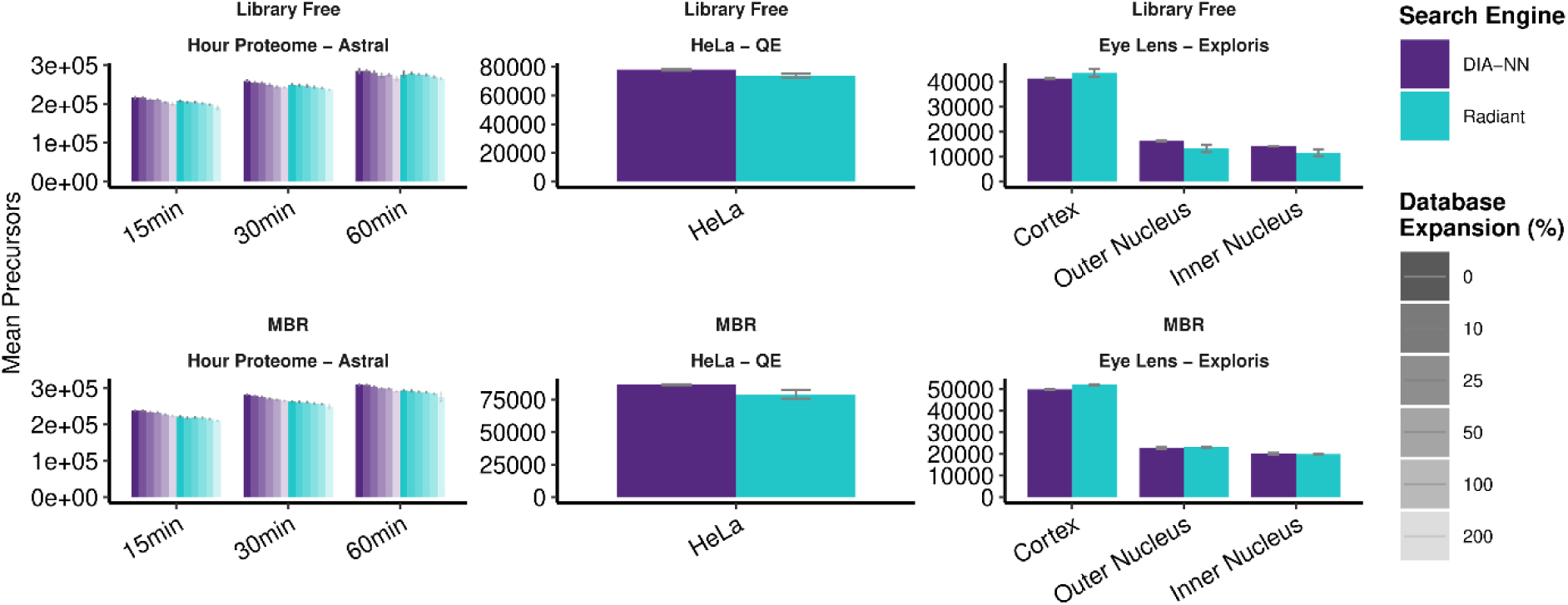
Overview of precursor identification counts from Hour Proteome, HeLa, and Eye Lens datasets.

However, the sensitivity of a pipeline must be assessed in tandem with error rate control for type I error prevention by way of *q*-value verification.^13,25^ We assessed calibration of PSM- and precursor-level *q*-value estimates using entrapment analysis from the 100% and 200% database expansion searches of the Hour Proteome dataset, calculating lower-bound and upper-bound (“combined”) estimates of the true false discovery proportion (FDP) as defined by Wen et al.^13^ Estimates for each analysis were classified as valid, inconclusive, or invalid according to whether these bounds were below, surrounded, or above the nominal *q*-value estimates. At the precursor level Radiant DIA results demonstrated valid FDR control in 8 of 12 analyses, with the remaining analyses inconclusive. In contrast, DIA-NN’s estimates were invalid in 3 of 12 analyses, inconclusive in 6, and valid in 3 (Supplementary Fig. 2).

We also evaluated PSM-level FDR estimates via entrapment. All library-free Radiant DIA analysis demonstrated valid PSM-level FDR estimates, while only 3 of 6 DIA-NN analyses were valid, with the remainder inconclusive (Supplementary Fig. 3). Assessment of PSM-level FDR control is particularly relevant to MBR workflows, because false transfer is a run-specific error: a precursor may be accepted in a recipient run because of cross-run matching. Because both Radiant DIA and DIA-NN perform MBR as a full second-pass search and assign *q*-values to all resulting PSMs, PSM-level entrapment directly evaluates calibration of error rate control in MBR results. We therefore interpret PSM-level entrapment FDP as an entrapment-based assessment of the false-transfer burden in the final MBR result, where false transfer is defined as a false accepted run × precursor match. In MBR analyses, both Radiant DIA and DIA-NN results showed valid PSM-level FDR estimates in 5 of 6 analyses, with the remaining analyses inconclusive. We note that this entrapment analysis is not a gold-standard true-absence-transfer assessment. Such an assessment would require an experimental design in which analytes are present in donor runs and known absent in recipient runs, such as a two-proteome or spike-in true-absence design. Therefore, our analysis supports calibrated MBR *q*-values and controlled entrapment-based false-transfer burden at the PSM level, but it should not be interpreted as a direct measurement of biological FTR into known-absence samples.

These findings indicate that Radiant DIA scoring maintains accurate error-rate control for PSMs and precursors, including PSM-level control of false-transfer burden in MBR workflows, while preserving identification sensitivity. Recent versions (2.6 at time of writing) of DIA-NN reportedly implement more conservative FDR handling^43^, but licensing restrictions limit its broad applicability and preclude evaluation in this work.

At the protein group level, DIA-NN typically reports a larger number of protein groups at 1% FDR (Supplementary Figs. 4, 7, and 12). However, protein group counts are difficult to interpret and do not necessarily indicate a meaningful difference in identification rate or biological content, as differences in protein inference methodology can lead to significant differences in the number of identified groups. The numbers presented in these analyses reflect each workflow’s inference implementations and protein group-level FDR control (see Supplementary Methods). Due to limited information regarding DIA-NN’s protein inference and scoring methodology and lack of decoy results it is not possible to apply matching inference to both sets of results.

Notably, DIA-NN tends to identify a larger number of single-peptide protein groups (“one-hit wonders), accounting for most of the difference in protein group count (Supplementary Figs. 7 and 12).

### Quantitative Precision and Accuracy

Run-to-run quantitative precision was evaluated using precursor-level coefficients of variation (CV) in the Hour Proteome dataset (Figure 3a). Across all gradient lengths and in both library-free and MBR modes, Radiant DIA quantification results in lower median precursor CVs than DIA-NN. When the analysis was restricted to identifications shared between engines, the difference in median CV narrowed but remained in favor of Radiant DIA, indicating that its quantitative measurements are not only deep but also reproducible across repeated injections. Together, these results support Radiant DIA as a precise platform for precursor-level quantification.

**Figure 3.**
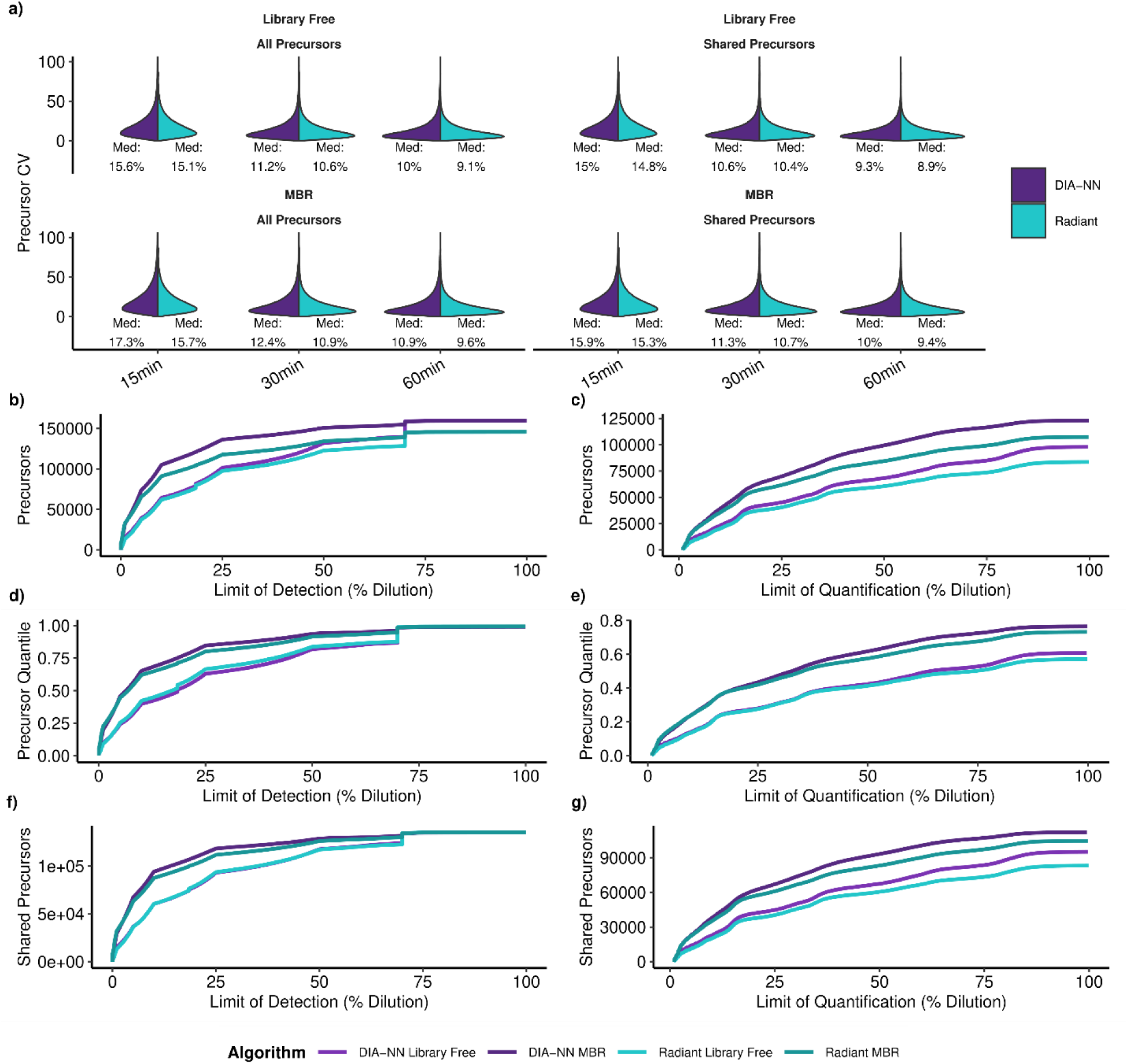
**A)** Quantitative precision across replicates for precursors in the Hour Proteome dataset. **B,C)** Count of precursors with LoD or LoQ at or below given level (fraction of endogenous concentration). **D,E)** As in B,C, but showing the fraction of precursors within each analysis. **F,G)** As in B,C, but showing only precursors identified in all analyses.

We next evaluated quantitative accuracy using mixed-proteome samples analyzed with a Matrix-Matched Calibration Curve (MMCC2) approach.^8^ This framework enables direct estimation of precursor-level quantitative performance metrics, including limits of detection and quantification (LoD and LoQ). Figure 3b-g and Supplementary Fig. 5 summarize these MMCC-derived measurements for Radiant DIA and DIA-NN in the Astral and Q-Exactive HF datasets, respectively.

Across both library-free and MBR analyses, Radiant DIA demonstrated robust quantitative performance, with LoD and LoQ profiles that generally tracked those of DIA-NN across dilution ranges and feature subsets. Although DIA-NN was numerically favored in several aggregate

MMCC summaries, Radiant DIA results show strong quantitative behavior across conditions, indicating that its performance remains practically competitive for both detection and quantification. We observe similar performance in a “LFQBench”-style mixed-species analysis, though this covers only a small number of fixed fold change values (Supplementary Fig. 6).^7,44,45^ Taken together with the improved run-to-run precision observed above, these results support Radiant DIA as a quantitatively capable and reliable workflow for large-scale DIA proteomics experiments.^8,36^

### Case Study: Eye Lens

The eye lens is a highly specialized, heterogeneous tissue organized in concentric layers of fiber cells formed throughout life. As lens fiber cells mature, they lose their organelles and persist indefinitely, accumulating age-related PTMs over decades of aging including pervasive deamidation.^46^ The small mass shift associated with deamidation (+0.984016 Da) poses substantial challenge for DIA identification algorithms as it competes against the nearby isotopic peak, making this dataset an ideal stress test for evaluating non-labile PTM handling.

We reanalyzed a previously published dataset (acquired on Thermo Fisher Orbitrap Exploris™ 480) consisting of three lens regions – Cortex, Outer Nucleus and Inner Nucleus – and compared identification performance across modified and unmodified peptides (Supplementary Fig. 7). In total, Radiant DIA search identified more precursors than DIA-NN in both library-free and MBR workflows, and reports a larger absolute number of candidate deamidated precursors, particularly in MBR searches, recapitulating the expected biological gradient from cortex to nucleus.^47^ The increased number of candidate deamidations is disproportionate to the increase in precursor sensitivity, indicating that it is not due to overall differences in sensitivity, bur rather due to differences in Radiant DIA‘s identification and scoring of PTMs. Although DIA-NN reports more protein groups, this difference likely reflects divergent strategies for protein inference, including treatment of single-peptide protein groups, as discussed above.

To compare match quality, we extracted chromatographic traces in Skyline and computed dot products between observed and library entries with respect to search engine reported chromatographic windows (Figure 4). Across all lens regions, Radiant DIA results have higher median dot product for precursors identified by both engines (-0.7 – 2.1% improvement).

**Figure 4.**
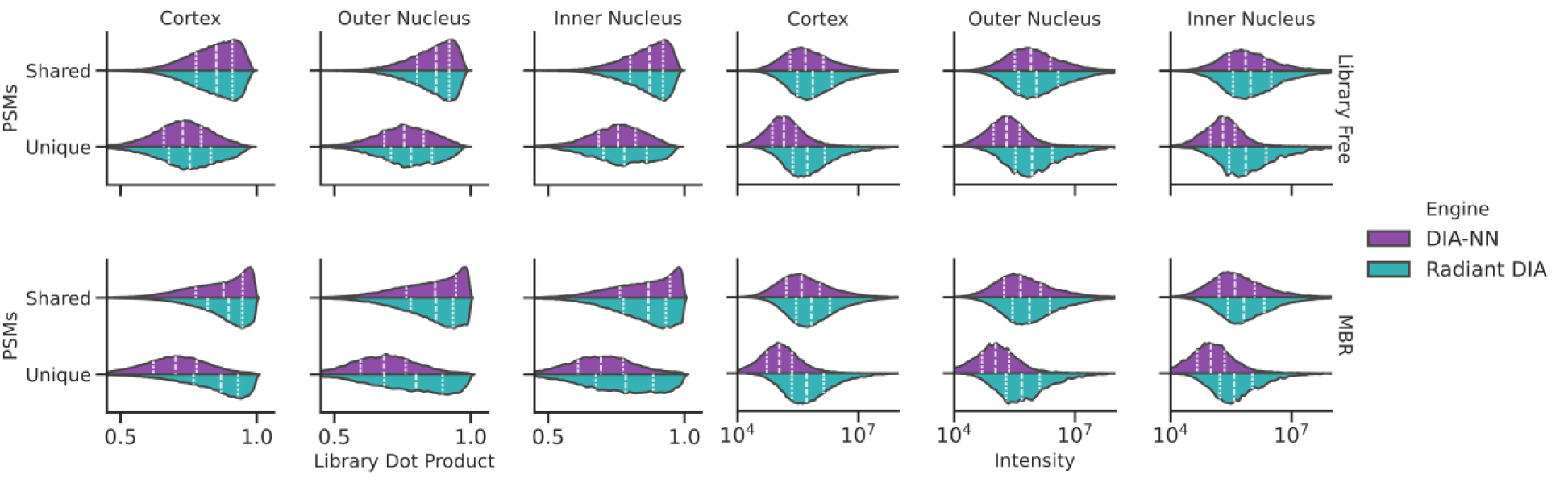
Distribution of spectral match quality assessed by Skyline (left) and measured intensity (right) for PSMs in the Eye Lens dataset. For each workflow, PSMs are grouped by whether a given precursor was identified in each acquisition by both pipelines. Lines within each distribution indicate the position of quartiles.

Differences were more pronounced for engine-unique precursors: in library-free mode, unique Radiant DIA matches exhibited 4-7% higher absolute median dot products than DIA-NN’s, and in MBR mode this gap widened substantially, with a 22-23% difference in medians. Trends remained consistent when restricting analysis to candidate deamidated precursors (Supplementary Fig. 8), highlighting robust PTM handling in this challenging dataset.

The quantitative evidence further supports these observations. Although Radiant DIA precursor intensities are slightly higher overall – with unique identifications generally less intense for both tools – DIA-NN’s unique identifications tend to fall at very low intensities, suggesting increased susceptibility to matching noise or low-intensity peaks.

We also assessed evaluated precursor matches which were putatively identified as deamidated by each workflow by extracting and classifying MS2 spectral support for the assignment and localization of deamidation, and whether there is evidence for an alternative interpretation such as a different localization or an isotopic signal (for details, see Supplementary Methods). Across lens regions, and library-free or MBR searches, a larger proportion and absolute number of Radiant DIA candidate deamidated precursors have spectral support for deamidation as compared to DIA-NN (Supplementary Figs. 9 and 10). The same trend is observed for candidate deamidation identifications shared between engines or uniquely identified by only one, indicating that Radiant DIA is better able to consistently identify signals with spectral evidence supporting the presence of deamidation modifications.

To better understand differences in Skyline-assessed spectral dot product and spectral evidence for deamidation, we also assessed retention time residuals for identifications shared between engines (Supplementary Fig. 11). Although most precursors align well between engines, deamidated species exhibit substantially larger RT residuals between Radiant DIA and DIA-NN (up to ∼10 min on a 90-min gradient). These differences indicate that RT selection is a cause of the reduced degree of spectral evidence for deamidation among DIA-NN’s shared precursor IDs, consistent with less stable localization of match or misassignment. This effect may be exacerbated by common structural isomers found in the lens that exhibit non-systematic RT shifts.^48^ Results for both unmodified and deamidated precursors indicate that Radiant DIA has stronger spectral and retention time support for its identifications. We also observe Radiant DIA maintains match quality as assessed by Skyline during evidence transfer, whereas DIA-NN’s transferred identifications exhibited deteriorated support. Together, these results indicate that Radiant DIA search gives well-supported and consistent identifications of both modified and unmodified peptides, reinforcing its strengths in complex, PTM-rich biological systems.

### Case Study: Cancer Comparison

To evaluate performance in a typical label-free discovery study, we re-analyzed a plasma dataset containing 20 cancer and 20 healthy subjects prepared with the Proteograph XT assay and acquired on an Orbitrap Astral MS (80 total injections).^49^ Data were processed using the non-redundant SwissProt library as well as 100% and 200% expansion libraries using local execution for DIA-NN and the cloud deployment of Radiant DIA and Fulcrum Pipeline. Identification performance at matched 1% PSM-, precursor-, and protein-group FDR thresholds is summarized in Supplementary Fig. 12. Precursor counts were comparable between engines (Supplementary Fig. 13) with DIA-NN reporting more single-peptide protein groups, accounting for 6-9% of its reported identifications.

We next compared run-to-run quantitative stability. Distributions of unnormalized precursor CVs were similar between engines (Figure 5), reflecting the substantial biological heterogeneity among plasma samples. Both pipelines showed increased CV under MBR, but DIA-NN displayed a larger shift towards high-CV precursors. This may indicate more accurate measurement of the dataset’s heterogeneity and noise due to more comprehensive measurement across samples, that transferred identifications include more variable or lower-confidence matches, or a combination of these effects.

**Figure 5.**
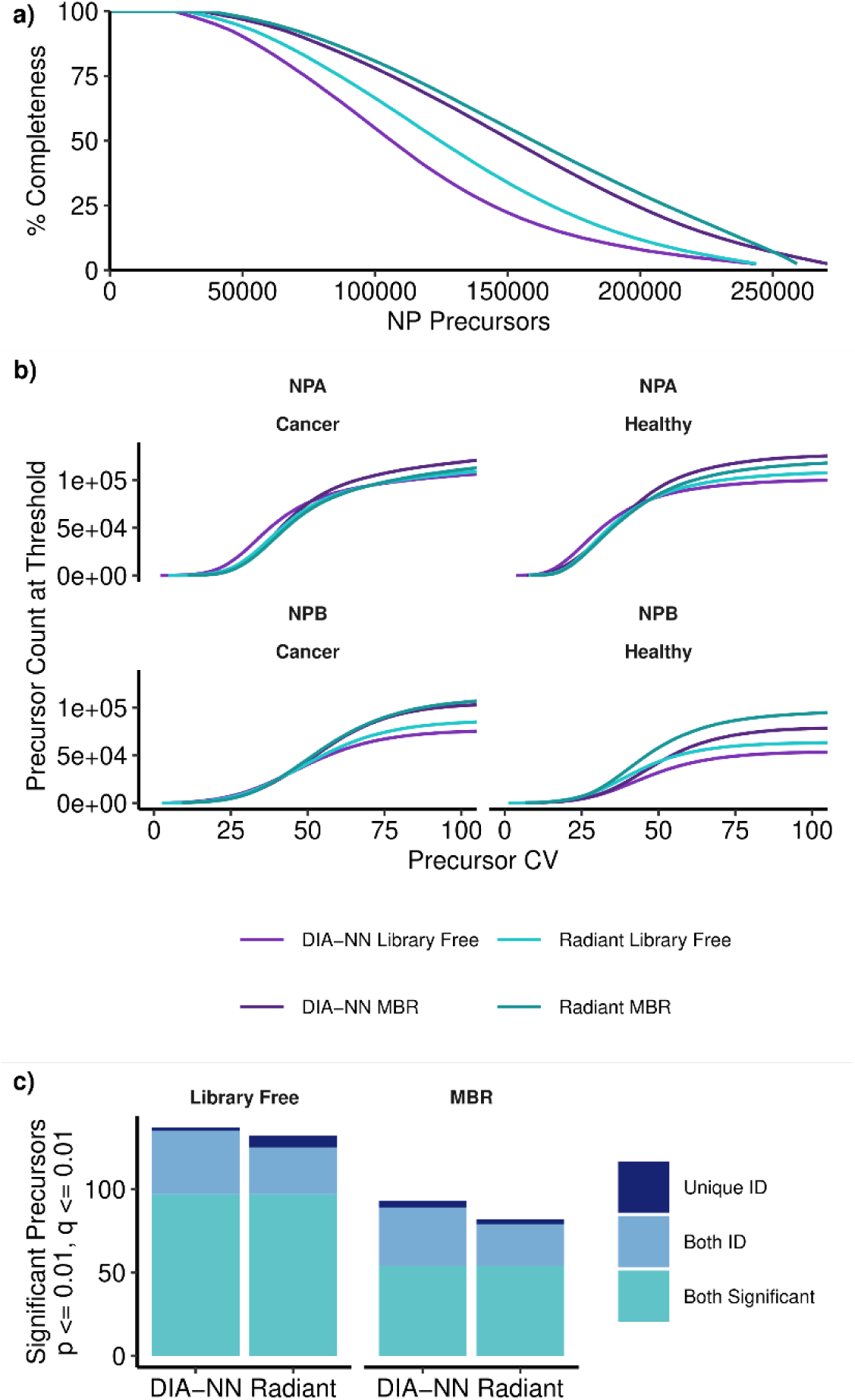
Evaluation of Cancer case study. A) Nanoparticle (NP) precursor-resolved completeness for each workflow. B) Engine-reported normalized intensity, separated by Seer NP (NPA or NPB), study group and workflow. C) Differential expression overlap comparison from NPA. Features filtered to p < 0.01 and q < 0.01. Features were categorized by ID similarity and annotated for indicators of poor quantification if retention times had high variance or elution before peak separation occurred on column. A large fraction of differentiation can be explained by potentially poor quantification.

To assess discovery performance, we conducted differential-abundance (DA) analysis across all precursor-nanoparticle pairs, controlling the DA false discovery rate at 1%. Most significant precursors were shared between engines, with each tool identifying few additional candidates not detected by the other (Figure 5). Both tools gave similar numbers of discoveries after excluding those with high retention time variability; such discoveries were more frequent in DIA-NN (Supplementary Fig. 14). Discordant RT selection across samples may indicate incorrect feature selection in some samples, which will typically increase quantitative variability, consistent with higher CV distributions. This may shift estimates of variability and effect size and result in increased erroneous DA feature discovery. While RT variability does not conclusively show that a discovery is inaccurate, it can serve as a qualitative evaluation of discovery quality and aid in prioritizing follow-up investigations.

The reduction in significant precursors under MBR for both pipelines likely reflects more complete measurement of variable signals, decreasing statistical power in a relatively small study. In contrast, library-free workflows are more susceptible to left-censoring of low-abundance precursors, which can artificially lower variance and inflate significance. Together, these results illustrate that Radiant DIA maintains robust downstream quantitative behavior in a heterogeneous biological cohort, as assessed by differential-abundance discovery concordance and reduced association with high-retention-time-variability features, while providing computation efficiency advantages for large-scale discovery studies.

### Example: Astral RCs

To demonstrate the scalability and cost-efficiency of Radiant DIA and Fulcrum Pipeline processing for large cohort studies, we processed a dataset of 2,561 run-control (RC) injections, comprising two pooled human peptide mixtures analyzed across four Orbitrap Astral instruments over an eight-month interval. These stable peptide mixtures provide an ideal test bed for measuring computational scalability independent of biological variability (Supplementary Fig. 15).

Both cloud workflows successfully processed the dataset; however, DIA-NN’s single-node execution model limits its ability to scale far beyond this study, particularly for high-depth sample sets where memory requirements become prohibitive.^19^ In contrast, Fulcrum Pipeline distributes workloads across many nodes with near-linear scaling, allowing thousands of acquisitions to be processed in parallel. Radiant DIA and Fulcrum Pipeline cloud processing of all 2,561 files finished in 3.99 and 2.42 hours of real “wall” time, with end-to-end cost of approximately $111 and $110 for library-free and MBR analyses, respectively (Figure 1).

DIA-NN, running in the cloud-distributed Scalable MBR workflow previously described, required approximately 1.92x more wall time and at least 9.5x higher cloud costs to complete the same analyses. Wall time analysis reflects limitations of cloud resource availability, as well as start-up and orchestration overhead, resulting in lower fold improvement as compared to costs.

Overall cost of execution reflects both the amount of provisioned compute resources and the time for which they were utilized, demonstrating the notable improvement in computational efficiency of the combined Radiant DIA and Fulcrum Pipeline workflow. For large datasets, costs are dominated by parallel processing of individual acquisitions, where Radiant DIA search steps can use fewer resources while still executing more quickly (Supplementary Fig. 1). Details of provisioned resources are provided in the Supplementary Methods.

## Discussion

In this work we introduce Radiant DIA and Fulcrum Pipeline, a unified pipeline for high-throughput identification and quantification in DIA-MS proteomics that integrates a fast, transparent search engine with a modular, distributed pipeline for re-scoring, FDR control, protein inference, and quantification. A central motivation behind this pipeline is the growing mismatch between advancements in MS instrumentation – yielding faster acquisition rates, deeper coverage, increasing data volume, and sample throughput – and computational capacity of existing analysis tools. Radiant DIA and Fulcrum Pipeline address this gap by enabling rigorous and scalable DIA analysis across both local and cloud environments with minimal configuration overhead.

Across datasets spanning diverse biological and analytical challenges, Radiant DIA results show PSM and precursor identification sensitivity comparable to DIA-NN 1.8.1 while exhibiting more accurate and conservative error-rate calibration, as demonstrated through entrapment analysis of PSM and precursor results. In these same datasets, unique identifications reported by DIA-NN frequently corresponded to lower-intensity signals, elevated quantitative variability, and reduced chromatographic or spectral agreement, suggesting a greater susceptibility to more permissive scoring behavior. In contrast, unique Radiant DIA identifications were consistently anchored by higher-quality evidence across PSM, precursor, and protein-group levels, reducing dependence on single-peptide protein groups and improving downstream biological interpretability.

The present evaluation was performed on Thermo Fisher q-analyzer instruments, including both Orbitrap and Astral analyzers. Performance on other instrument platforms remains an important area for future validation. Because Radiant DIA accesses DIA spectral information through `mzML`, we expect its computational efficiency relative to DIA-NN 1.8.1 to extend to conventional DIA datasets from other platforms, such as SCIEX SWATH acquisition. However, identification sensitivity, error-rate estimation, and quantitative performance may depend on instrument-specific acquisition properties and should be assessed directly. Radiant DIA does not currently support ion-mobility-aware workflows (including Bruker timsTOF diaPASEF data), or direct analysis of vendor-native formats such as Thermo Fisher `.raw`, Bruker `.d` or SCIEX `.wiff` files. Extending Radiant DIA to ion-mobility-enabled data will require development of ion-mobility-aware parsing and search functionality.

A key strength of this pipeline is its computational efficiency. Optimized Radiant DIA scoring and minimal overhead for MBR enable 5-10x faster execution than DIA-NN on equivalent hardware, while Fulcrum Pipeline enables order of magnitude reductions in cloud compute cost via scalable, parallel orchestration of thousands of acquisitions. This efficiency lowers both financial and practical barriers to performing large DIA studies, making population-scale proteomics, multi-center projects, or iterative reprocessing efforts more accessible. Importantly, Fulcrum Pipeline retains a single codebase for desktop and cloud execution; switching between environments requires only a change in workflow configuration, not infrastructure engineering.

The engineering philosophy of Radiant DIA and Fulcrum Pipeline tools emphasize analytical transparency, exposing raw scoring features, configurable decoy retention, and structured intermediate outputs that enable reproducibility, method development, and deeper algorithmic inspection. This transparency, combined with Fulcrum Pipeline’s modular architecture, facilitates rapid integration of alternative scoring models, peak-integration strategies, or advanced statistical tools – a critical need as proteomics evolves and machine-learning-based approaches continue to advance.

Taken together, these results demonstrate that Radiant DIA and Fulcrum Pipeline constitute a fast, scalable and statistically robust solution for DIA-MS data analysis. As MS moves towards population-scale profiling, single-cell proteomics, and clinical translation, computational demands will continue to intensify. By providing an efficient and extensible platform that maintains rigorous FDR control for PSMs and precursors, high quantitative accuracy and seamless scalability, Radiant DIA and Fulcrum Pipeline help make these next-generation DIA applications practical and economically feasible. We anticipate that this framework will support not only routine high-throughput studies but also enable new methodological innovations, ultimately accelerating biological discovery and improving the interpretability of large-scale proteomics experiments.

## Data Availability

Results from this work’s analyses are available via the MassIVE repository with the identifier MSV000101464.

Community datasets are available from the ProteomeXchange or MassIVE repositories with the identifiers (i) Hour Proteome:^50^ PXD049028, (ii) HeLa:^10^ MSV000082805, (iii) Eye Lens:^47^ MSV000087506, (iv) Astral MMCC:^51^ PXD042704, (v) QE MMCC:^8^ PXD014815, (vi) Cancer Study:^49^ `PXD060573`. Raw MS data from the Astral RC dataset is used in this work to demonstrate the performance characteristics of our pipeline when processing large datasets and thus it has not been made publicly available, as it is of limited scientific value.

## Code Availability

Code and binaries for Radiant DIA and Fulcrum Pipeline are publicly available and freely usable under an Apache 2.0–based source-available license.

Radiant DIA and Fulcrum Pipeline can be run via a Docker container, available from github.com/seerbio/radiant-fulcrum-container

An optional graphical interface providing interactive access to this workflow is available at github.com/seerbio/radiant-fulcrum-gui

Complete source code for Radiant DIA, Fulcrum Pipeline, and associated plugins are available on Github in the following repositories:

github.com/seerbio/radiant

github.com/seerbio/fulcrum

github.com/seerbio/radiant-fulcrum-workflow

github.com/seerbio/radiant-fulcrum-search

github.com/seerbio/wheely-radiant

github.com/seerbio/wheely-mammoth

github.com/seerbio/airpotgithub.com/seerbio/cortado

github.com/seerbio/proffergithub.com/seerbio/preppers

Cloud analyses described in this work were executed using a cloud execution plugin and deployment infrastructure that are not part of the public release.

Representative configuration files and notebooks used to generate results and figures in this manuscript are included in the results submission available on MassIVE.

## Supporting Information

Supplementary File 1:

- Supplementary Fig. 1: Benchmarks for single-file search step
- Supplementary Fig. 2: Precursor-level entrapment results
- Supplementary Fig. 3: PSM-level entrapment results
- Supplementary Fig. 4: Protein group counts with various filters
- Supplementary Fig. 5: QE MMCC results
- Supplementary Fig. 6: LFQBench results
- Supplementary Fig. 7: ID counts for Eye Lens analyses
- Supplementary Fig. 8: Dot product and intensity of deamidated Eye Lens PSMs
- Supplementary Fig. 9: Count of deamidated Eye Lens PSMs by support class
- Supplementary Fig. 10: Proportion of deamidated Eye Lens PSMs by support class
- Supplementary Fig. 11: RT residuals between workflows for Eye Lens PSMs
- Supplementary Fig. 12: Counts for Cancer Study analyses
- Supplementary Fig. 13: Overlap of precursors and peptides for Cancer Study analyses
- Supplementary Fig. 14: Significant DA discoveries for Cancer Study analyses
- Supplementary Fig. 15: Identified precursor count for each injection of the Astral RC dataset
- Supplementary Fig. 16: Comparison of alternative library creation strategies
- Supplementary Fig. 17: Block diagrams of Radiant DIA and Fulcrum Pipeline
- Supplementary Methods describing the Radiant DIA search engine and Fulcrum Pipeline workflow implementation and evaluation (PDF).

Supplementary File 2: Annotated spectra showing illustrative examples of deamidated precursor identifications for each workflow (PDF).

## Author Contributions

Ideation: S.J., A.N., I.M., T.P., S.B. Implementation: S.J., A.N., J.K., T.P. Data Analysis: S.J., A.N., L.S.C. Manuscript Preparation: S.J., A.N., L.S.C., S.B. All authors contributed to critically reviewing the results and the manuscript.

S.J and A.N. contributed equally to this work, and I.M., J.W., T.P., S.B. and O.F. jointly supervised this work.

