## Supplemental File 1 for "Radiant DIA: A Fast, Sensitive, and Accurate Search Engine for Quantitative Proteomics"

#### Table of Contents

Supplementary File 1:

- Supplementary Fig. 1: Benchmarks for single-file search step
- Supplementary Fig. 2: Precursor-level entrapment results
- Supplementary Fig. 3: PSM-level entrapment results
- Supplementary Fig. 4: Protein group counts with various filters
- Supplementary Fig. 5: QE MMCC results
- Supplementary Fig. 6: LFQBench results
- Supplementary Fig. 7: Counts for Eye Lens analyses
- Supplementary Fig. 8: Dot product and intensity of deamidated Eye Lens PSMs
- Supplementary Fig. 9: Count of deamidated Eye Lens PSMs by support class
- Supplementary Fig. 10: Proportion of deamidated Eye Lens PSMs by support class
- Supplementary Fig. 11: RT residuals between workflows for Eye Lens PSMs
- Supplementary Fig. 12: Counts for Cancer Study analyses
- Supplementary Fig. 13: Overlap of precursors and peptides for Cancer Study analyses
- Supplementary Fig. 14: Significant DA discoveries for Cancer Study analyses
- Supplementary Fig. 15: Identified precursor count for each injection of the Astral RC dataset
- Supplementary Fig. 16: Comparison of alternative library creation strategies
- Supplementary Fig. 17: Block diagrams of Radiant DIA and Fulcrum Pipeline
- Supplementary Methods describing the Radiant DIA search engine and Fulcrum Pipeline workflow implementation and evaluation.

Supplementary File 2: Annotated spectra showing illustrative examples of deamidated precursor identifications for each workflow.

#### Supplementary Figures

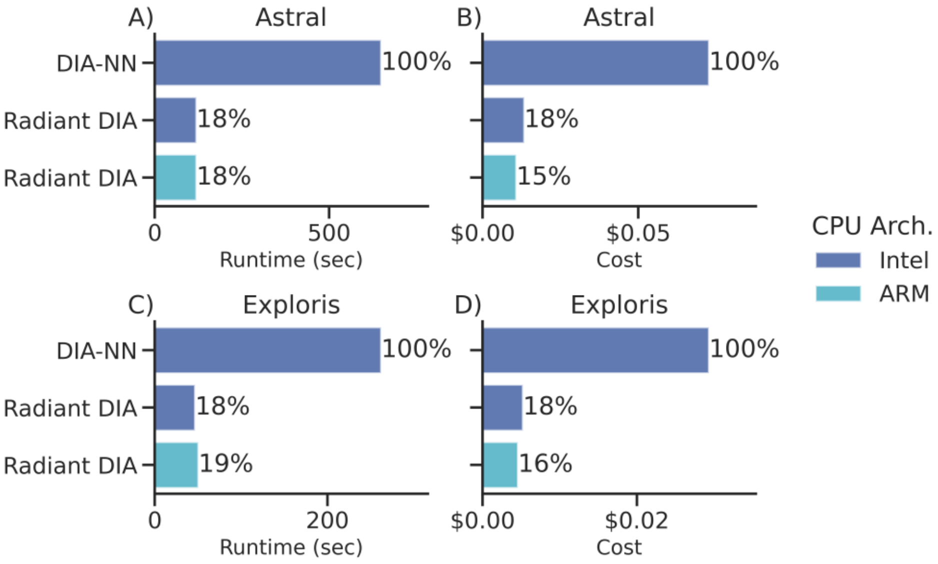


***Supplementary Figure* 1.** *Benchmarks for DIA-NN and Radiant DIA search step using m7[ig].2xlarge instances with 8 cores and 32 GiB of memory. Percentages are calculated relative to the maximum runtime or cost for each instrument.* ***A,B)*** *Runtime and cost for search of a single 30 min Astral acquisition, not including data transfer or global processing steps.* ***C,D)*** *Runtime and cost for search of a single 30 min Exploris acquisition.*

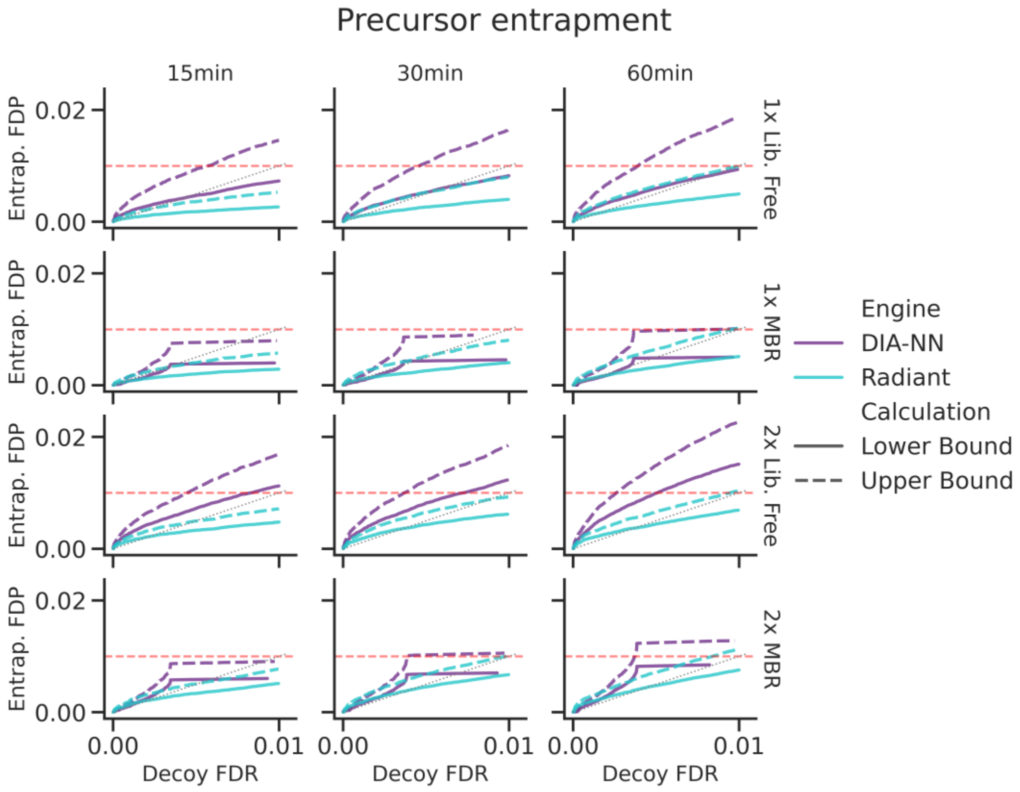


***Supplementary Figure 2.*** *Precursor-level entrapment results for Hour Proteome analyses.*

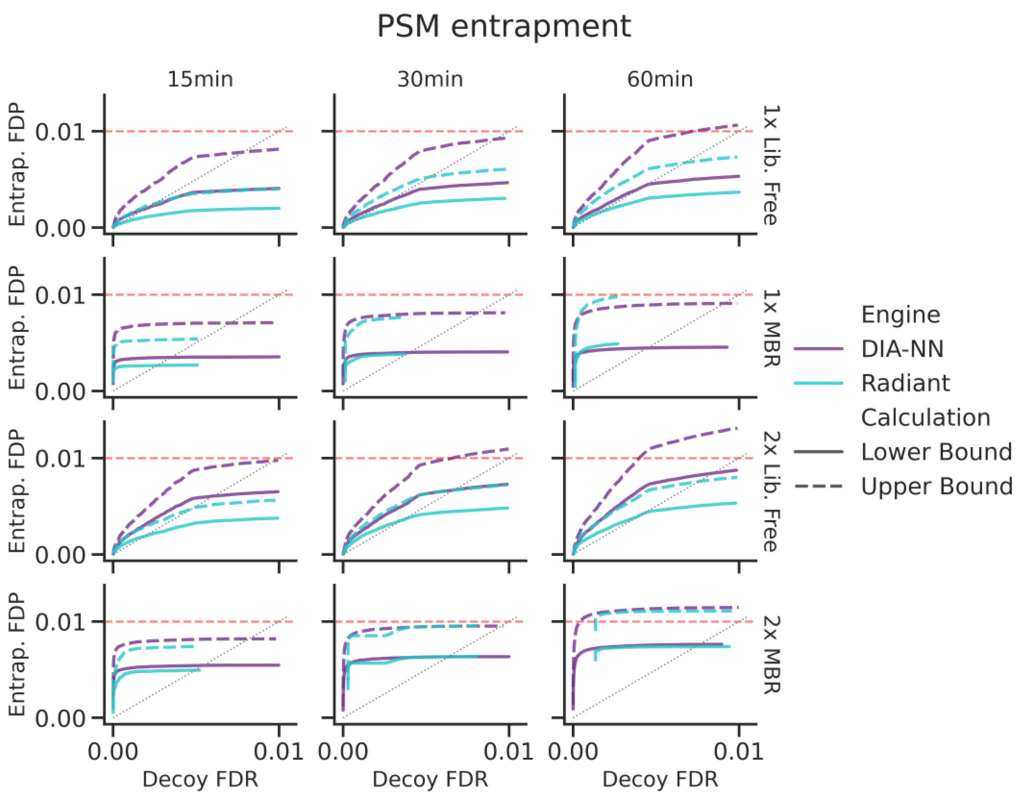


***Supplementary Figure 3.*** *PSM-level entrapment results for Hour Proteome analyses.*

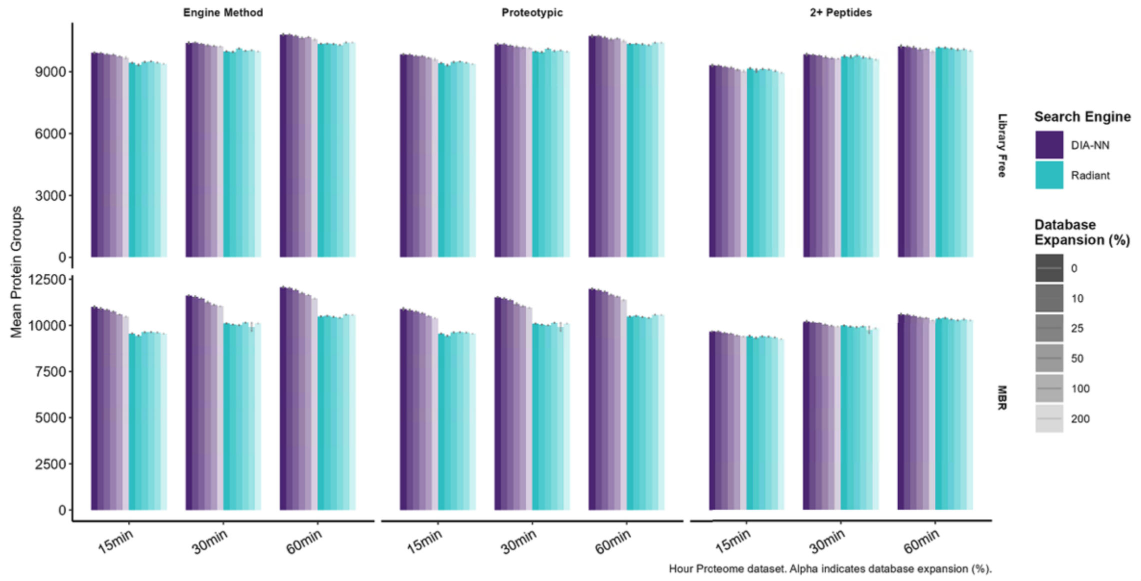


***Supplementary Figure 4.*** *Protein group counts for Hour Proteome analyses under each engine's FDR filters, when requiring proteotypic support for the group, and when requiring at least 2 peptides.*

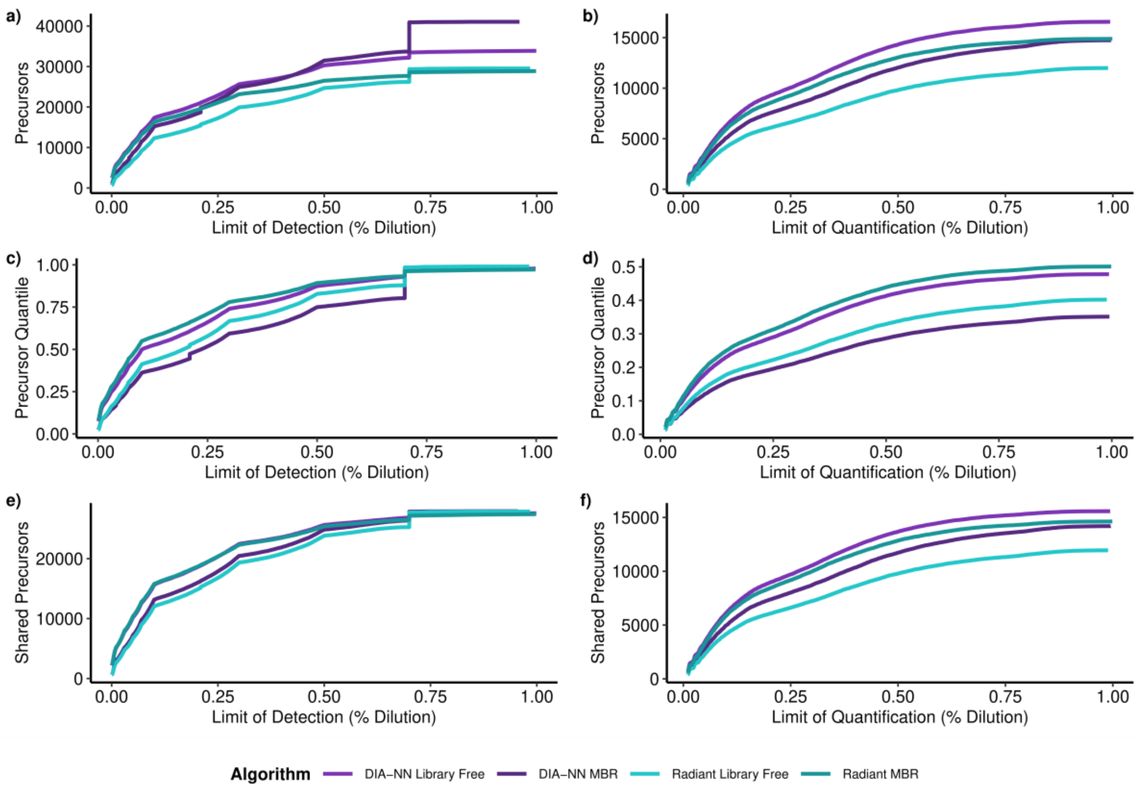


***Supplementary Figure 5.*** *Summary of LoD and LoQ results for QE MMCC dataset.* ***A,B)*** *Count of precursors with LoD or LoQ at or below given level (fraction of endogenous concentration).* ***C,D)*** *As in A,B, but showing the fraction of precursors within each analysis.* ***E,F)*** *As in A,B, but showing only precursors identified in all analyses.*

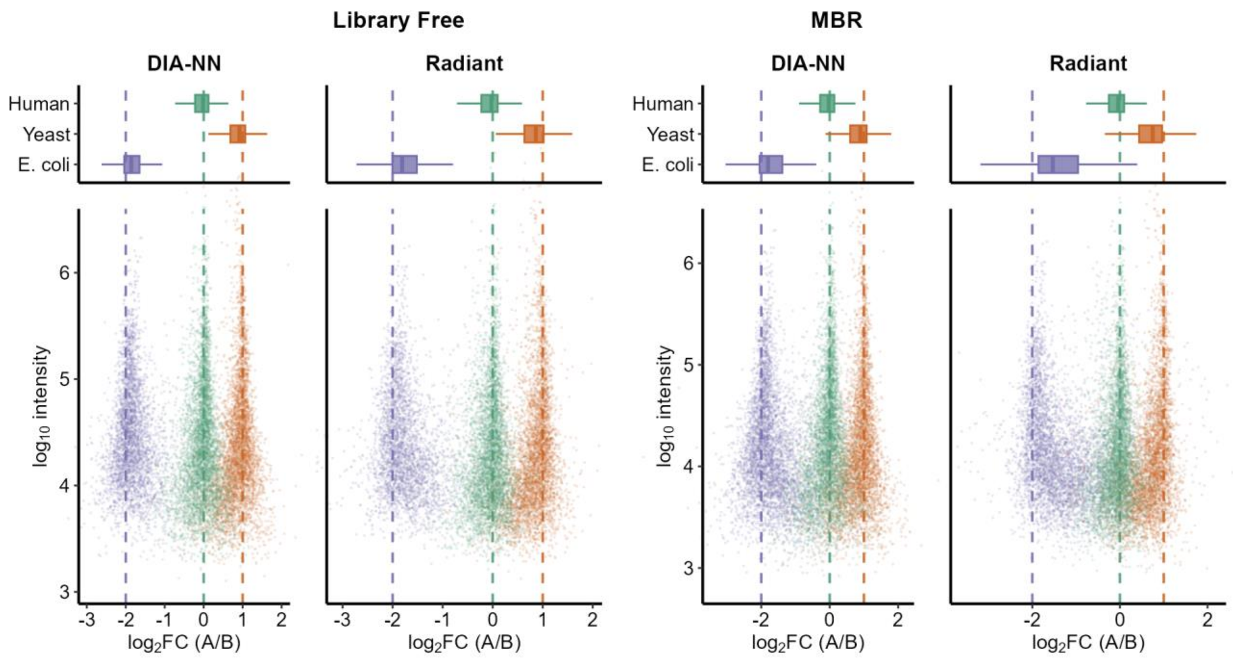


***Supplementary Figure 6.*** *Summary of fold change estimates for precursors in the mixed-species dataset from LFQBench dataset, generation beta. Both engines demonstrate ratio compression for low-intensity signals belonging to*

*the Yeast and E. Coli proteomes; this effect is slightly more pronounced for Radiant DIA, likely because no quantitative transition refinement is employed when quantifying precursors.*

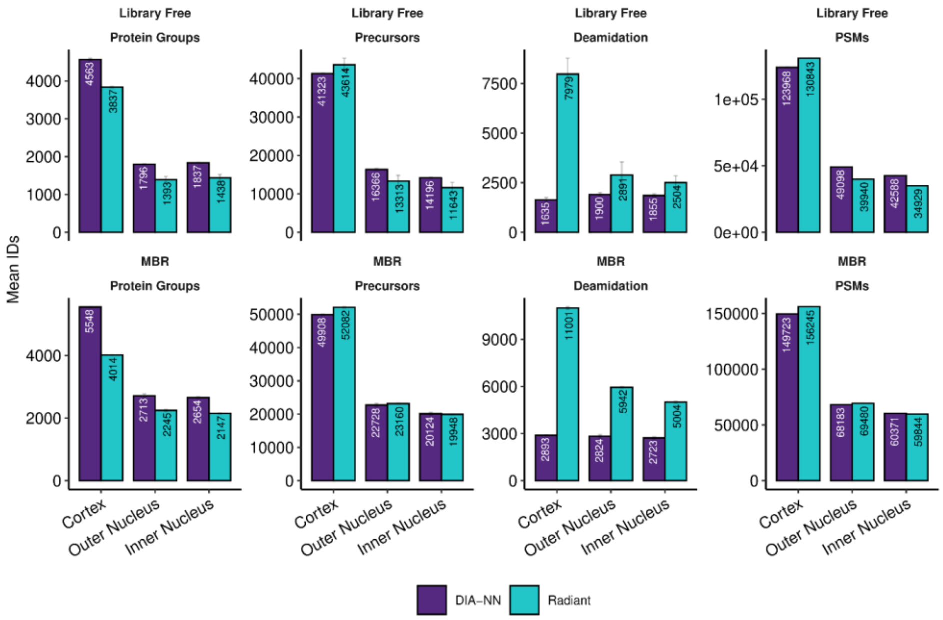


***Supplementary Figure 7.*** *Aggregate identifications for each lens region for Library Free (top) and MBR (bottom) workflows. Protein Groups, Precursors, and Deamidation show the number of unique identifications globally. PSMs shows the total number of precursor identifications summed across replicates.*

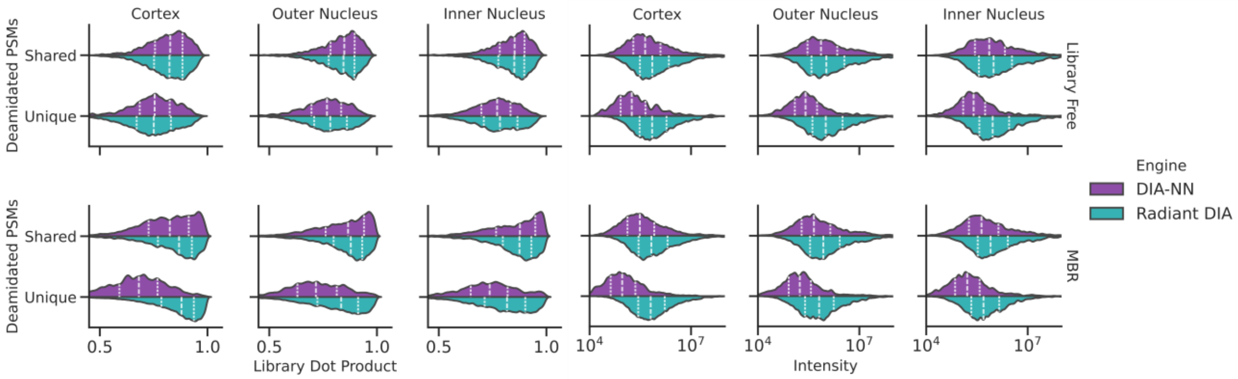


***Supplementary Figure 8.*** *Distribution of spectral match quality assessed by Skyline (left) and measured intensity (right) for deamidated PSMs in the Eye Lens dataset. For each workflow (top: library free, bottom: MBR), PSMs are grouped by whether a given precursor was identified in each acquisition by both pipelines. Lines within each distribution indicate the position of quartiles.*

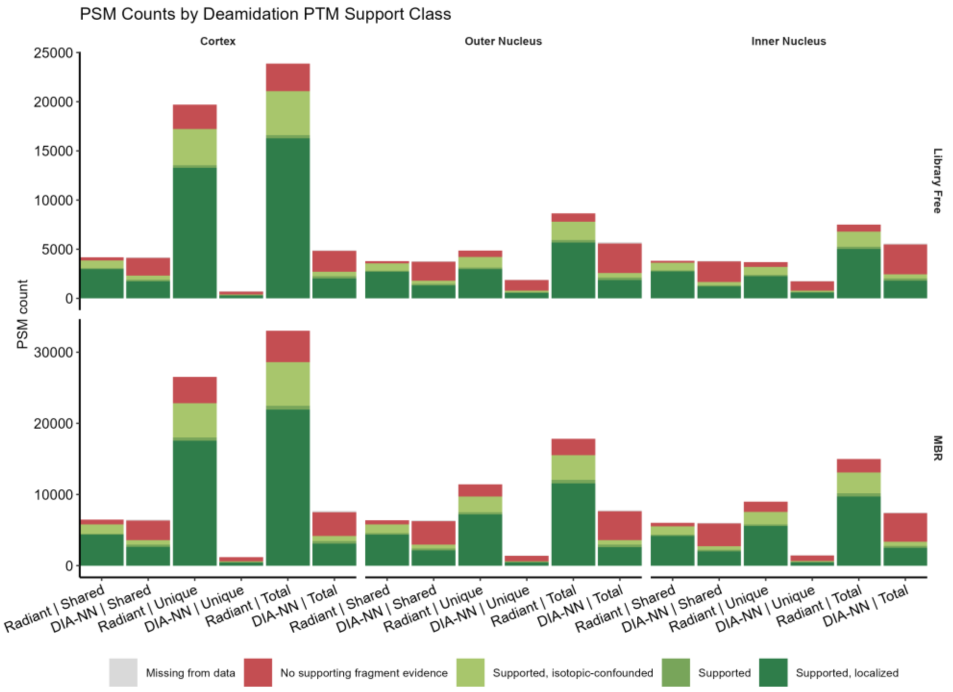


***Supplementary Figure 9.*** *Assessment of spectral support for candidate deamidated PSMs across eye lens regions and worklows, showing the number of PSMs within each category. For each workflow and region, separate bars show PSMs identified by both engines, those uniquely identified by a single engine, as well as the total number. Radiant DIA consistently shows a greater absolute number of PSMs with spectral evidence supporting deamidation.*

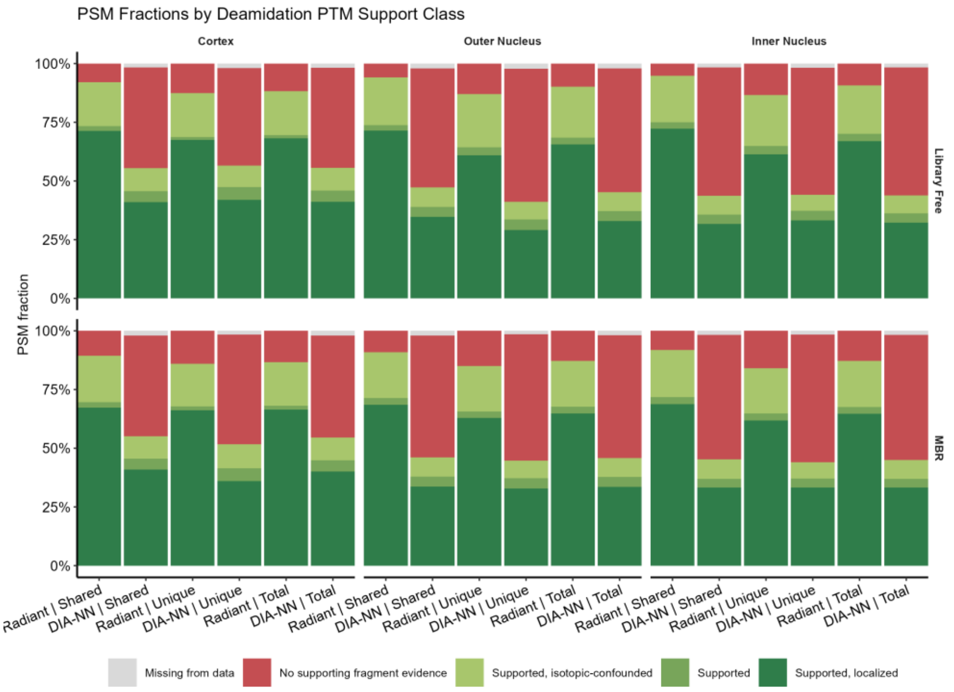


***Supplementary Figure 10.*** *Assessment of spectral support for candidate deamidated PSMs across eye lens regions and worklows, showing the number of PSMs within each category. For each workflow and region, separate bars show PSMs identified by both engines, those uniquely identified by a single engine, as well as the total number. Radiant DIA consistently shows a greater proportion of PSMs with spectral evidence supporting deamidation.*

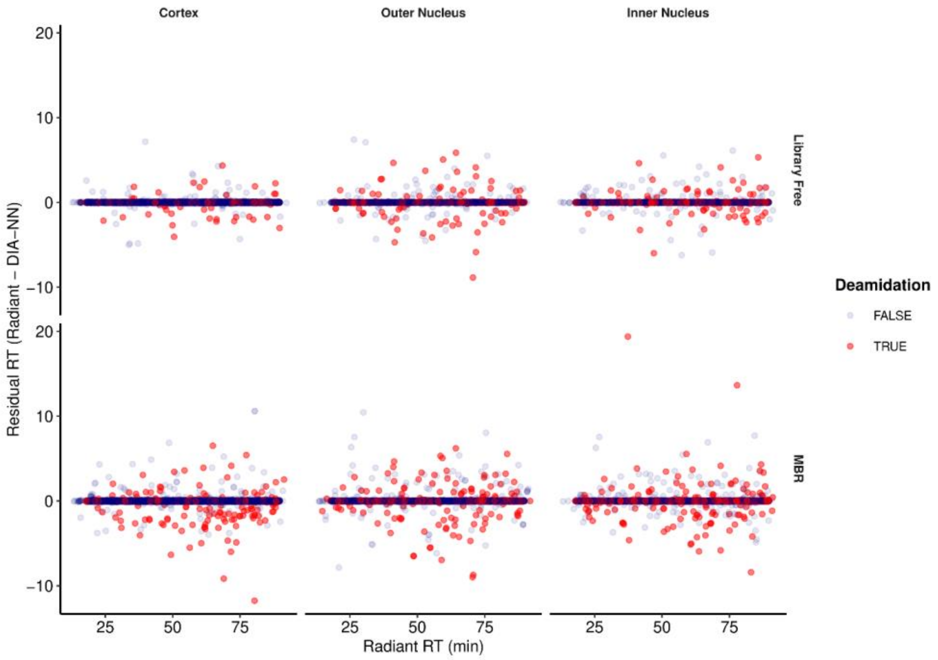


***Supplementary Figure 11.*** *Agreement of detected precursors' retention times between Radiant DIA and DIA-NN. While RTs for precursors without deamidations are largely consistent within one minute, deamidated precursors are detected with greater RT residuals, especially in MBR mode. Differences in RT assignment are likely responsible for different levels of spectral support for deamidation among shared identifications*.

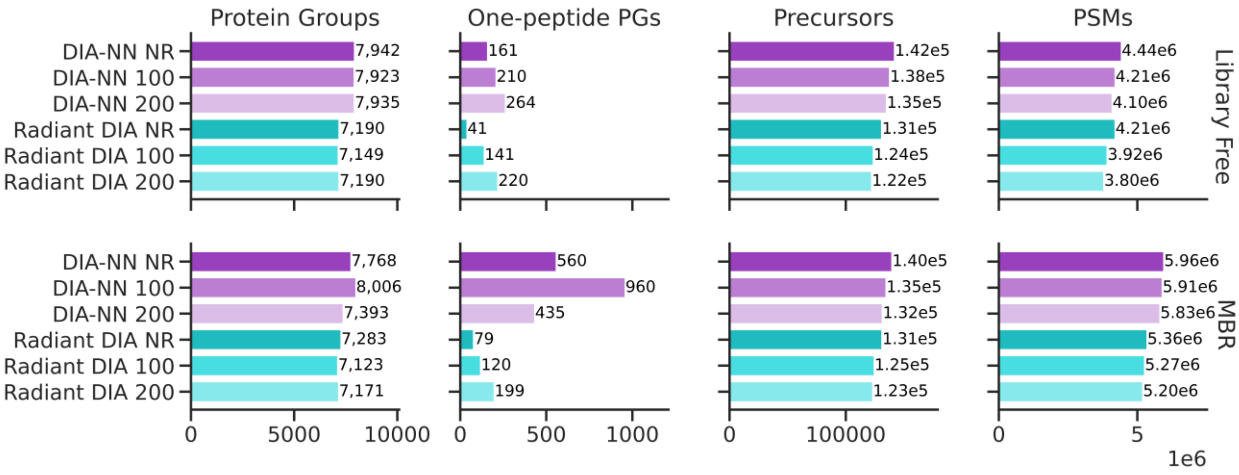


***Supplementary Figure 12.*** *ID counts at 1% PSM-, precursor- and PG-level FDR for* Cancer study*. From left to right: Number of protein groups (PGs), number of PGs identified by a single peptide, number of precursors, total number of PSMs.*

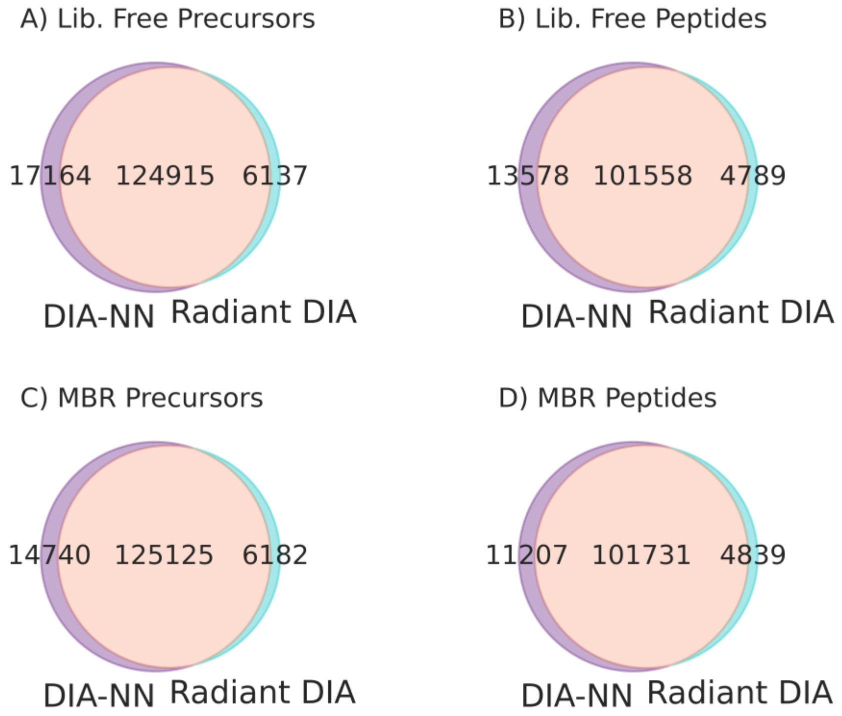


***Supplementary Figure 13.*** *Overlap of identifications at precursor and peptide levels, for library free (A,B) and MBR (C,D) searches of Cancer Study acquisitions with NR library. Results are not filtered for PG-level FDR, demonstrating the overlap of identifications prior to protein inference.*

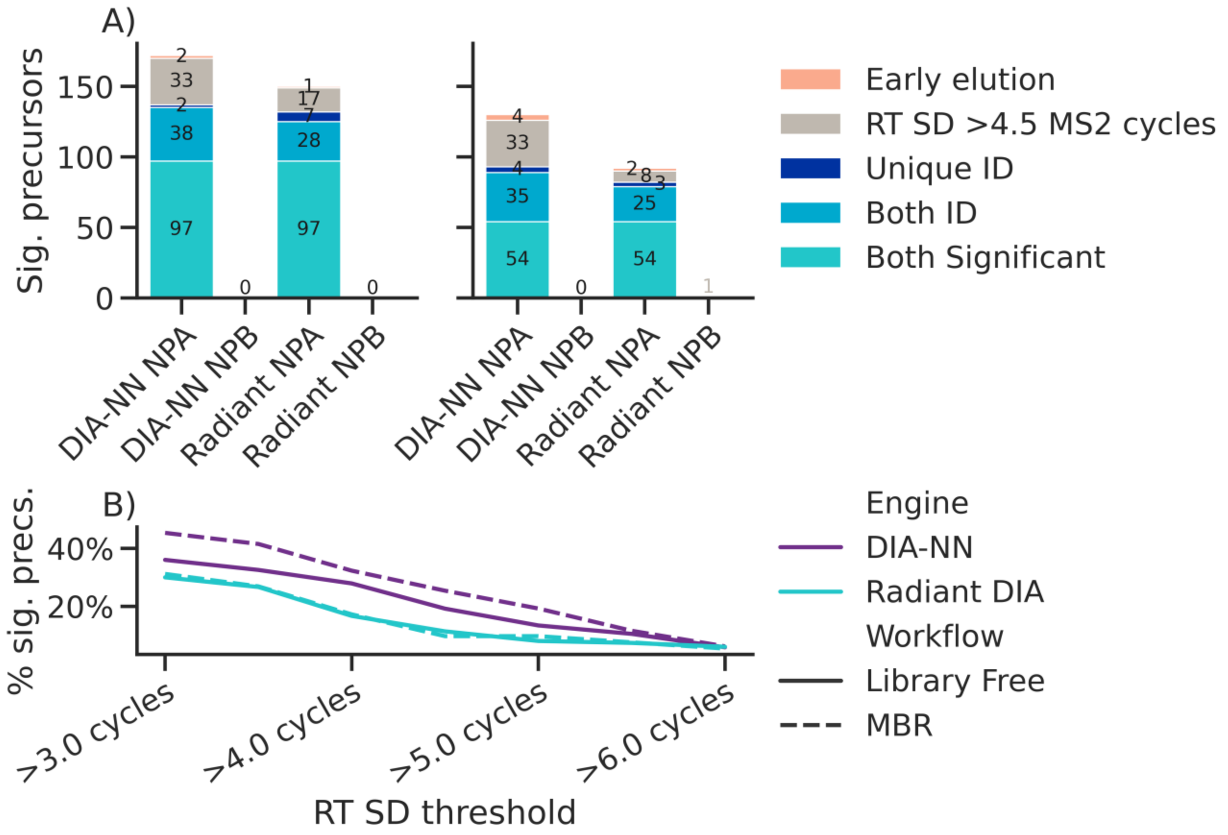


***Supplementary Figure 14.*** *A) Number of significant differentially abundant precursors identified for both nanoparticles in the Cancer Study searched with NR library, grouped by whether the precursor was identified as eluting during the column void volume (before 9 min RT), had an RT standard deviation greater than 3.375 sec (>4.5 MS2 scan cycles), or was uniquely identified or significant in a single engine's results. B) Fraction of significant precursors for each workflow with RT SD exceeding various thresholds, demonstrating that a smaller proportion of Radiant DIA discoveries have discordant RTs as compared to DIA-NN.*

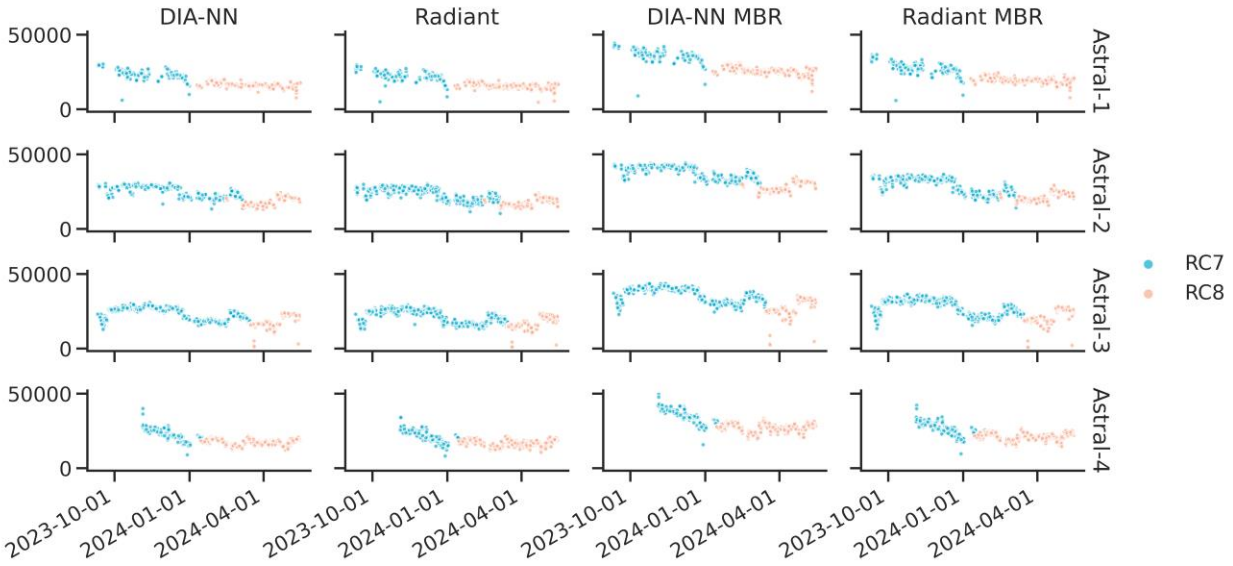


***Supplementary Figure 15.*** *Number of identified precursors for each of 2,561 RC injections, acquired over an eight-month period on four Astral instruments. Two different mixtures ("RC7" and "RC8") were employed.*

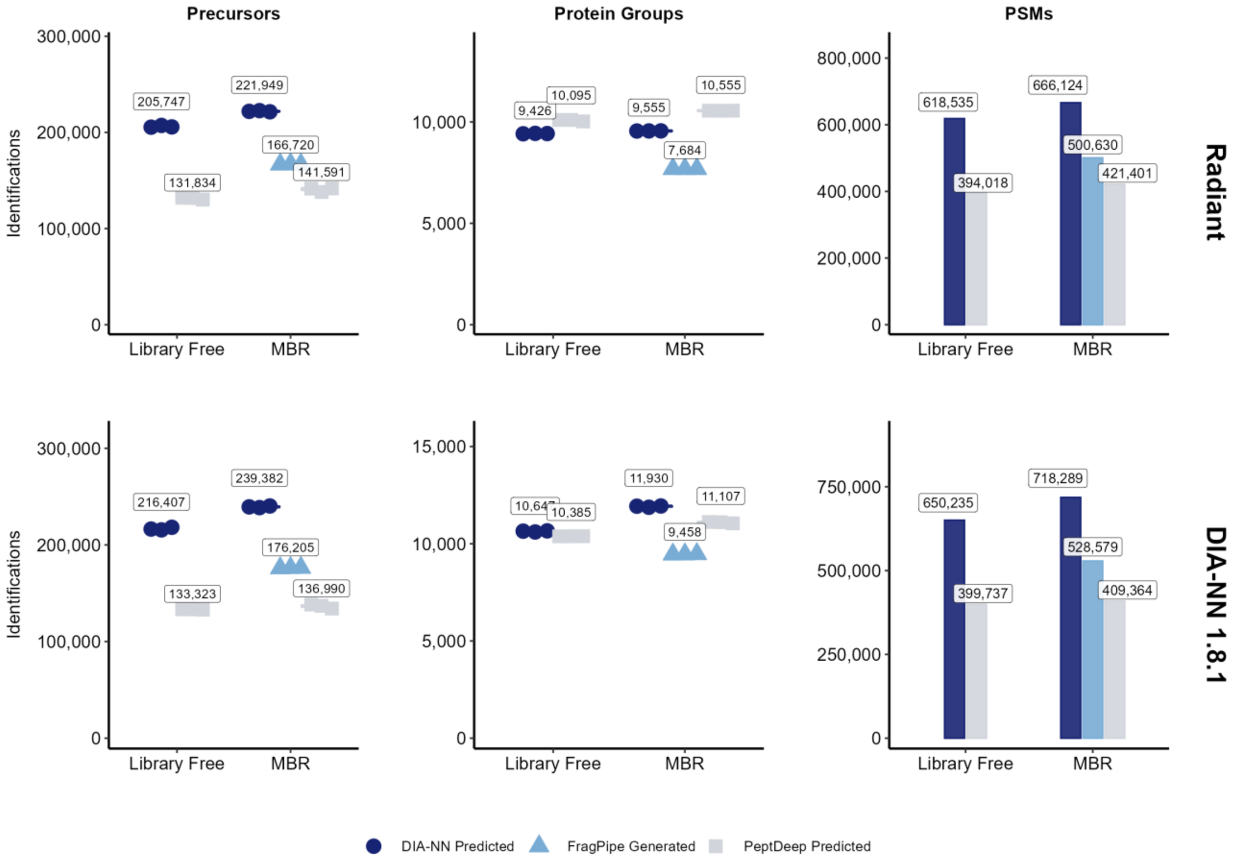


Hour Proteome dataset, 15 min gradient.

***Supplementary Figure 16.*** *Number of identified PSMs, precursors, and protein groups for three library generation strategies, demonstrating that Radiant DIA and DIA-NN have similar performance trends with various library intputs, enabling diverse workflows and experiment-specific library optimizations.*

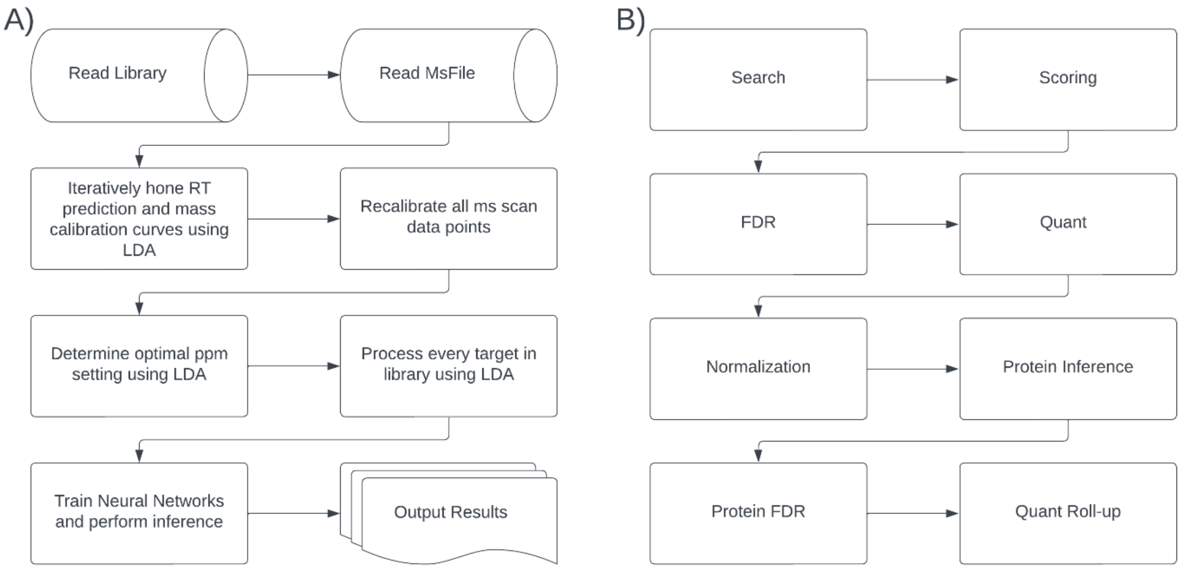


***Supplementary Figure 17. A)*** *Block diagram of Radiant DIA search engine, showing high-level steps taken while processing each individual acquisition (file).* ***B)*** *Block diagram of Fulcrum Pipeline workflow. These steps can be performed by a variety of modular implementations, allowing multiple approaches to function within the same overall workflow.*

### Supplementary Methods

#### Radiant DIA Search Engine

Radiant DIA processes each file independently, making the search process parallelizable, and thus suitable for scalable execution in a cloud environment. To process a single file, first input files are read. For each target in the library, a corresponding decoy is generated by mutating the sequence at the second and penultimate residues, using the mutation pattern ACDEFGHIKLMNPQRSTUVWY → LSEDLLSVLVLQLNLTSSLLS; fragment ion masses are adjusted according to the corresponding mass shifts. Small batches of the library are then scored by LDA to build an RT alignment model and determine an optimal RT window filter by using splines generated by a specified number (default is 1000) of PSMs at 5% FDR. Results from these batches are cached and used to compute a mass calibration curve, which is then used to recalibrate the full set of MS data. The calibration curve is calculated using least squares to fit a polynomial, the order of which is specified by the user (default is 2$^{nd}$ order). Subsequently, mass tolerances for extracting transition traces are optimized by processing a small batch of library targets at values ranging from 3 to 50 ppm. This is done in an ascending fashion and is halted when target counts do not peak above a maximum after three further iterations. Mass tolerance optimization is performed after mass recalibration to ensure the optimal tolerance accounts for measurement precision and does not compensate for inaccurate mass measurements.

Using the optimized RT window, recalibrated spectra, and optimized mass tolerance, the full library is then scored using LDA. Precursor quantities are computed by summing detected fragment intensities over all scans where the total intensity of all utilized fragments is greater than a configurable fraction (65% in this manuscript's analyses) of the total in the highest single scan.

The NN classifier is trained using all PSMs passing a 50% FDR threshold, or the top 50,000 PSMs, whichever is larger. The classifier consists of three fully-connected layers with a configurable hidden layer width (defaulting to half the number of scoring features) using ReLU activation, and is trained using binary cross-entropy loss and a multi-fold cross-validation scheme that ensures each PSM was not used to train the network that scores it. Cross-fold normalization (as in Percolator) is applied (in log space) to ensure scores are well-calibrated between folds.[1–3]

After NN training, PSM FDR estimates are recomputed based on the NN classifier scores, and results are written to the output Parquet file with a user-configurable FDR threshold.

#### Fulcrum Pipeline Orchestration Engine

##### *PSM and Precursor FDR Estimation by Blocking MixMax*

Blocking MixMax consists of three steps. First, the score space is quantized into "blocks" using Spark's `QuantileDiscretizer`, which efficiently divides the score space into bins which each contain roughly equal numbers of input PSMs. By controlling the number of blocks and the accuracy of the binning it is possible to control the trade-off between speed of calculation and granularity of this quantization. In practice we have found that using 20,000 to 100,000 blocks and relative error of 0.0001 gives acceptable performance and granularity of *q*-value estimates. Second, after discretization, the number of targets and decoys within each bin is computed; these counts can be efficiently computed in a distributed fashion. In the final step, a single-threaded Python calculation (using Numpy and Numba's `njit`[4] for performance) performs the MixMax calculation.[5] This calculation is very similar to the implementation from `crema`[6] (which itself is a near-direct translation of the published algorithm[5] into Python), with two key modifications. First, it employs the target and decoy counts within each block rather than counting each row with equal weight. Second, the algorithm is modified to simultaneously compute two *q*-value estimates in a single pass; the two estimates handle the effectively tied scores between all PSMs in a single block in opposite ways, either interpreting decoys as scoring better or worse than the targets, giving (respectively) upper- or lower-bound estimates for *q*-value. This calculation is guaranteed to give results that bound the result of MixMax computed without quantization, and when quantization gives exactly one PSM per block the upper and lower bounds are identical. Importantly, the computational complexity of this calculation is determined solely by the number of blocks employed in quantization. For a given number of blocks, the runtime of this step will be approximately constant for any number of input PSMs. While the complexity and runtime of quantization and count aggregation will grow with PSM count, the increase will be sub-linear, and these calculations can be easily distributed, ensuring that blocking MixMax can efficiently handle thousands of MS acquisitions.

Blocking MixMax also permits efficient estimation of posterior error probabilities (PEPs) for by computing the proportion of decoys within each block, applying a rolling average filter (with configurable width) and taking a cumulative maximum to ensure a monotonic relationship between score and PEP.

Results from blocking MixMax are assigned to PSMs within each block without interpolation. Upper- and lower-bound *q*-values are reported along with PEPs. In all downstream analyses the upper bound *q*-value is employed to ensure that filtering is conservative, though in practice there is very little difference between these estimates.

Global precursor FDR is computed by the same process as PSM FDR after selection of the best-scoring PSM for each precursor. The Fulcrum Pipeline also supports peptide-level FDR estimation, though this was not employed in this manuscript's analyses.

##### *Protein Inference, Scoring, and FDR Estimation*

In parallel, each connected component of the resulting bipartite peptide-protein graph is formed into groups by picking the protein sequence with the highest heuristic score. Several heuristics are available; in this manuscript we employ the "number of sibling peptides" ("NSP"),[7] computed as the sum of peptide probabilities (1 - PEP) associated with each protein. Proteins sharing the same set of peptides are grouped together. Grouping proceeds iteratively until all peptides are assigned to groups. This grouping process is performed globally to ensure peptides are consistently grouped across the entire experiment, and to avoid unnecessary computational overhead from repeated peptide annotation and protein grouping for each file.

After protein grouping, scores are calculated for each protein group. While it is theoretically possible to re-use calculated heuristic scores from grouping for scoring and FDR estimation, the Fulcrum pipeline utilizes a separate calculation to maximize flexibility. In analyses presented here, we employ NSP to score protein groups, which we found gave better-calibrated FDR estimates (as assessed by entrapment, described below) than alternative scores such as minimum *q*-value. Unlike the NSP heuristic used for protein grouping, protein group NSP used for confidence estimation is computed at the precursor level (though it is possible to instead use peptide-level NSP, this was not employed in this manuscript's analyses). Protein group FDR is then estimated globally using either TDC (employed in analyses presented here) or MixMax *q*-value calculation from `crema`.[6]

#### Local and Cloud Execution Environments

Radiant DIA and the Fulcrum Pipeline (henceforth: Radiant and Fulcrum) are designed for a wide range of environments, from a single computer to a cluster of many nodes. In this work, we employed either a local execution or cloud execution environment. The former runs all calculations in a single Docker container that contains Radiant and Fulcrum. The latter employs containerized Radiant searches running on a Kubernetes cluster and an auto-scaling Spark cluster for Fulcrum, each deployed within AWS. These environments differ markedly in capabilities and availability of compute resources, but there were only very minor differences in the code and configurations used to execute workflows. Other than the choice of search plugin used in the Fulcrum workflow, the same Fulcrum modules and configurations were used in both environments.

For local execution, a Docker image was prepared containing Radiant and Fulcrum, as well as runtime dependencies such as Python and Spark. For each analysis, a container was configured with appropriate volumes, and a TOML file passed to the Fulcrum CLI. Radiant was run sequentially on all input MS files using the local execution plugin described above. Fulcrum output Parquet files were written to local storage. For local execution analyses presented in this manuscript, we employed a `c7a.8xlarge` AWS instance with 32 CPUs and 64 GiB of memory. The same instance type was used to perform DIA-NN analyses.

Cloud execution requires additional setup and orchestration, which was automated within Seer's AWS environment. This system took a list of input MS files (stored in AWS S3) and a TOML configuration and set up an auto-scaling Spark cluster using Databricks. Necessary Python libraries, including Fulcrum and the cloud Radiant search plugin were installed in the cluster, and the Fulcrum workflow executed using the provided parameters. Radiant searches were run using the cloud search plugin, which executed them in parallel, using one container per MS file in an AWS EKS cluster. Radiant results were written to S3 in Parquet format and read using Spark for subsequent workflow steps. Fulcrum results in Parquet format were also written to S3.

Radiant DIA/Fulcrum Pipeline cloud execution analyses presented in this manuscript used the following configuration:

- Radiant DIA searches (each MS file): 8 CPU / 32 GiB memory
- Spark Cluster:
  - 1 `i3.2xlarge` driver node (8 CPU / 61 GiB memory)
  - 1-20 `i3.2xlarge` worker nodes (8-160 CPU / 61-1220 GiB memory)

Note that the use of autoscaling ensured that only resources necessary for execution were allocated to the Spark cluster. The utilized configuration provides sufficient resources to scale to experiments of thousands of files with reasonable performance, without incurring unnecessary costs for smaller runs. While we did not specifically track the number of nodes used for each analysis, aggregate costs for execution of each analysis were collected from AWS.

For comparison purposes, we also evaluated our cloud-distributed DIA‑NN pipeline, running either in single-pass mode, or using our optimized MBR pipeline, termed "Scalable MBR".[8] While this pipeline also employs parallel processing of individual files where possible, it requires that results be collected onto a single processing node at one or more points during processing. Scalable MBR reduces the resource requirements but does not permit the same level of distributed processing or autoscaling possible with Fulcrum. For DIA-NN cloud analyses presented here all steps were performed in containers running on AWS EKS with the following resource allocations:

- DIA-NN searches (each MS file): 16 CPU / 64 GiB memory
- Global processing (generation of MBR library, global FDR estimation and reporting): 74 CPU, 750 GiB memory
- Quant normalization: 48 CPU, 512 GiB memory
- Quant rollup: 64 CPU, 750 GiB memory

Due to the complexity of cloud pipelines and the multi-factorial nature of optimizing their configuration, no effort was made to align the resource configurations between the Radiant/Fulcrum and DIA-NN pipeline deployments. We believe that due to the large differences in the implementation and scalability of the two pipelines, configuring the two pipelines with equal resources would result in suboptimal performance for one or both. Instead, these configurations were optimized to balance cost, runtime, reliability, and the ability to efficiently handle runs of thousands of MS acquisitions.

#### Search Engine Evaluation

Each dataset was processed using Radiant DIA within the Fulcrum Pipeline and DIA-NN (version 1.8.1) in both single-pass ("library-free") and two-pass ("MBR") modes. In both modes, searching was initial performed against a library of predicted spectra (described below). All raw files were converted to `mzML` format prior to searching and all searches were performed using `mzML` files. Unless otherwise noted, community datasets were searched using local execution, and Seer datasets searched within Seer's cloud processing environment (both described above).

##### *Search Space and Spectral Prediction*

For all non-entrapment or database-expansion searches, a FASTA database of human proteins (SwissProt, Trembl, and UniParc isoform) was compiled on 2025-08-04 with added contaminants from the MaxQuant database. This database consists of 75,071 protein sequences, with high redundancy due to isoform inclusion. No decoys were added to this database. For Hour Proteome database-expansion analyses, the non-redundant human SwissProt database (20,659 entries) was downloaded from UniProt on 2025-05-01. To add decoys, sequences from the database were shuffled with the Decoy PyRat tool.[9] The tool generates per protein decoys and partial subsets of decoys were appended to the non-redundant FASTA according to database expansion size. For the expansion of 200%, a shuffled database was shuffled a second time, being careful to reject permutations that were in the target or 100% expansion database. For entrapment studies, the 100% and 200% expansion databases were used.

To enable fair comparison of pipelines, spectral prediction was performed with DIA-NN 1.8.1 using default parameters provided in the DIA-NN GUI. For each FASTA, a DIA-NN `speclib` file was generated. Time spent generating predicted libraries was not included in any reported benchmark results, as the libraries can be re-used.

##### *DIA-NN Search*

For both local and cloud searches DIA-NN was run with the following parameters (or equivalent GUI options): `--qvalue 0.01 --matrices --missed-cleavages 1 --met-excision --cut K*,R* --smart-profiling --relaxed-prot-inf --reannotate --peak-center --no-ifs-removal`. All DIA-NN searches were run with the "Optimal Results" option to balance runtime, resource consumption and overall sensitivity.

##### *Local Search Time Comparison*

For local search, log files from DIA-NN were used to establish time to complete first pass and second pass searches. Bash scripting was used for Radiant DIA and the Fulcrum Pipeline to determine end-to-end search time of the process that is shared between multiple tools. The same computing resources were used for each search, as described previously. For hour proteome datasets each length of gradient was processed independently to demonstrate performance tradeoffs at different file sizes from the same MS instrument.

*Cloud Search Time and Cost Comparison*

For cloud searches, pipeline runtime was automatically recorded, covering the full duration of execution, including any start-up and orchestration overhead; this represents an “end-to-end” duration a user would encounter between triggering an analysis and having full results available. Resource utilization for cost estimation was tracked by tagging within AWS and Kubernetes. For Databricks clusters, both Databricks costs and AWS billing data in the associated account was gathered and associated with each run. For Kubernetes containers, allocated costs (including per-node idle costs) were gathered using `kubecost`.

*Identification Filtering and Count Comparison*

Results from all pipelines were similarly filtered to 1% FDR at the PSM, precursor, and PG levels. Unless otherwise noted, all results in this work employed all three levels of filtering. For DIA-NN this corresponds to the `Q.Value`, `Global.Q.Value` (or `Lib.Q.Value` for MBR), and `Global.PG.Q.Value` (or `Lib.PG.Q.Value` for MBR) columns.

*Assessment of FDR control by Entrapment*

To assess the accuracy of FDR estimates, we employed the entrapment technique.[10] We specifically compute lower- and upper-bound estimates of the proportion of incorrect targets (false discovery proportion, or FDP) at the PSM, precursor, or protein group level using the “lower bound“ and “combined“ approaches defined by Wen et al.[10] Briefly, decoys were removed from the list of discoveries and identifications were annotated as target or entrapment using all accessions assigned to each protein group. Entrapment *q*-values were estimated by a Python script implementing the calculations defined by Wen et al.[10] Each analysis was classified as having valid, inconclusive, or invalid FDR estimates if the computed FDP bounds at 1% *q*-value were (respectively) below, contained, or above the 1% threshold.

*Matrix Matched Calibration Curves*

For analysis of Matrix Matched Calibration Curves and determination of Limit of Detection and Limit of Quantification, scripts were downloaded from the repository at `https://github.com/lindsaypino/matrix-matched_calcurves/`. Because the 0% concentration inputs were excluded from search, the flag `--min_noise_points` 0 was added to command options. Result files from Radiant/Fulcrum and DIA‑NN in “long” format were converted to EncyclopeDIA-like “wide” format. Results from each tool were evaluated using PSMs achieving 1% thresholds on precursor and protein levels according to respective search paradigms.

##### *Mixed-Species Quantitative Experiments*

Results from mixed-species datasets from ProteomeXchange with dataset identifier PXD070049[11] were processed to compute fold changes between mixtures for each precursor using raw intensities (without normalization) from the A and B LFQBench Generation Beta datasets. The optimized Astral 15 minute method dataset was analyzed. The FASTA provided in the repository was used to predict a library (Supplemental Methods, Search Space and Spectral Prediction) and the search engine methods used for “Library Free” and “MBR” analyses in other analyses were applied. To account for errors in mixing, log fold changes from each analysis were adjusted to center the median fold change for Human precursors at the expected 1:1 value. Analyses were executed consistent with ProteoBench and prior standard analyses.[12,13]

##### *Skyline Evaluation*

To establish the quality of spectral match, results from DIA NN and Radiant/Fulcrum were converted with custom scripts and input to Skyline Daily[14] (version 24.1.1.254). For predicted library search, the `speclib` file was used for transition import. For second-pass search, the empirical libraries from respective tools were converted to `speclib` and input to Skyline. Retention times were aligned based on results files from respective search algorithm and search mode. From refined data where Skyline extracted peaks, the library dot product was exported. Features shared between algorithms in respective search modes and unique features were parsed, and comparison of aggregate performance shown.

##### *Assessment of Candidate Deamidation Assignments*

For candidate peptide-spectral matches putatively assigned as deamidated precursors we assessed the level of spectral support for the assigned deamidation state. For each candidate precursor passing *q*-value filters and annotated as deamidated, we extracted the reported apex MS2 scan from the corresponding `mzML` file and evaluated spectral support for the assignment. The modified sequence was parsed to identify the reported deamidation site and all possible asparagine (N) and glutamine (Q) candidate sites, and theoretical b- and y-type fragments were generated for the reported deamidated form, relevant alternative localizations, and corresponding unmodified +1 isotope forms. Observed peaks were matched to theoretical fragment m/z values within 15 ppm and were required to exceed an intensity threshold defined as the greater of 100 absolute intensity units or 0.5% of the spectrum base peak intensity. Deamidated and isotopic fragment signals were distinguished by calculating the expected *m/z* values for each form, accounting for fragment charge, and evaluating whether the observed signal was more consistent with the deamidated fragment or the +1 isotope counterpart on the basis of mass error. Candidate identifications were classified as supported when at least one diagnostic fragment supporting the reported assignment was stronger than the corresponding alternative-localization or +1 isotope signal, with no competing evidence observed; mixed when supporting evidence was present together with competing alternative-localization or +1 isotope evidence; weakly supported when supporting fragments were observed but were not stronger than competing evidence; and unsupported when only alternative-localization or +1 isotope evidence was observed. For a small number of matches no qualifying fragment evidence was detected, which were classified as “missing from data”. Automated spectral annotations were generated as examples for each search and identification quality classification with respect to predicted mass annotations and library support used for PSM.

##### *Alternative Library Analysis*

To evaluate the impact of predicted library selection on performance of Radiant DIA, search was executed using Radiant DIA and Fulcrum Pipeline or DIA-NN 1.8.1 as previously described, with alternate libraries substituted. Analysis was executed on the 15-minute gradient, Hour Proteome dataset with all search and FDR filtering kept identical otherwise.

To predict libraries with AlphaPept ecosystem, AlphaPeptDeep v1.4.1[16] was used with settings equivalent to DIA-NN predicted settings. The same FASTA used for Hour Proteome analyses was uploaded and prediction was executed for the ThermoTOF model, consistent with Astral data acquisition. Missed cleavages were set to 1, variable modifications were disabled, peptide lengths of 7-30 residues were allowed for precursor charges +1-4 with max +2 fragment charge. Up to 12 fragments were retained with relative intensity threshold 0.001. The library output was natively supported for search in DIA-NN and Radiant DIA. Library Free and MBR analyses were subsequently executed.

To generate a library with MSFragger within the FragPipe Plus v23.1 ecosystem[17] a first-pass search was performed in MSFragger DIA using the SpecLib Quant DIA pipeline, from which a library was generated for use in DIA-NN and Radiant DIA. To the FASTA used for hour proteome analysis, decoys were added using FragPipe Plus. As with

AlphaPeptDeep, equivalent search space was selected for FragPipe Plus as was for DIA-NN, notably missed cleavages were set to 1, variable modifications were disabled, peptide lengths of 7-30 residues were allowed for precursor charges +1-4 with max +2 fragment charge. MSBooster was run before Percolator, Protein Prophet and FDR filtering steps observed in the validation tab of FragPipe Plus using pre-specified workflows of the SpecLib Quant DIA method. A spectral library was generated using EasyPQP in the pipeline, using default methods. Because PSMs included in the library reflect a prior search, FragPipe analysis was considered analogous to "Library Free" in alternative library comparison. The exported library was natively readable in Radiant DIA and DIA-NN 1.8.1 with subsequent analysis performed consistent with second pass, "MBR" methods.

Comparison of IDs between various libraries was performed using custom R scripts.

##### *Significance Testing for Differential Abundance*

To assess each pipeline's quantitative discovery capability, we performed differential-abundance analysis with `limma`[15] on log transformed intensities from the Cancer Study, evaluating each precursor-nanoparticle pair independently across workflows and libraries. Identifications were first filtered to 1% FDR at PSM and precursor level and requiring fewer than 75% missing values in each group. Resulting *p*-values were converted to *q*-values and results filtered at 1% *q*-value to control the differential-abundance false positive rate.

#### Sample Preparation and DIA LC-MS Acquisition

For Astral run controls, two mixtures of pooled peptides from a variety of sources were prepared. Because these mixtures were used only to assess the consistency of mass spectrometry their exact contents were unimportant; they were meant to provide a large volume of consistent material that could be used over a long time scale. Both run control mixtures generally consist of human peptides derived from plasma utilizing various preparations. A master mix of peptides is generated once per year and aliquoted into many tubes that are stored individually at –80C in 0.1% formic acid and 3% acetonitrile.

Run Control peptides were analyzed using the ThermoFisher™ Orbitrap™ Astral™ mass spectrometer (ThermoFisher Scientific). Tryptic peptides were loaded on an Acclaim™ PepMap™ 100 C18 (0.3 mm ID x 5 mm) trap column and then separated on a 50 cm µPAC™ HPLC column (ThermoFisher Scientific) at a flow rate of 1 µL/minute using a gradient of 5-25% solvent B (0.1% FA, 100 % ACN) in solvent A (0.1% FA, 100% water) over either 13 or 22 minutes, resulting in a 22-minute or 33-minute total run time. 400 ng of peptides per injection was analyzed in DIA using 3 m/z isolation from 380-980 m/z. MS1 scans were acquired at 240k resolution on the Orbitrap analyzer every 0.9 seconds and MS2 at >80k resolution on the Astral analyzer. Ions were accumulated for up to 5 ms before measurement.
