## Supplemental File 2 for "Radiant DIA: A Fast, Sensitive, and Accurate Search Engine for Quantitative Proteomics"

### DIA-NN Library Free

Library cache: shared_libfree_predicted_library.parquet
Selected spectra: 7

Contents:
- 5 strong supported examples
- 1 supported, isotopic-confounded example
- 1 unsupported example

Selection emphasis:
- strong examples require Supported, localized calls with at least one observed localizing deamidation fragment that is explicitly predicted in the method-specific library
- isotopic-confounded examples retain supporting evidence but have stronger competing isotope-like evidence overall
- unsupported examples show no retained deamidation-supporting fragments in the exported spectrum

Experimental top, Predicted library bottom | RT 32.72 min | precursor m/z 912.4380
ID quality: Supported, localized

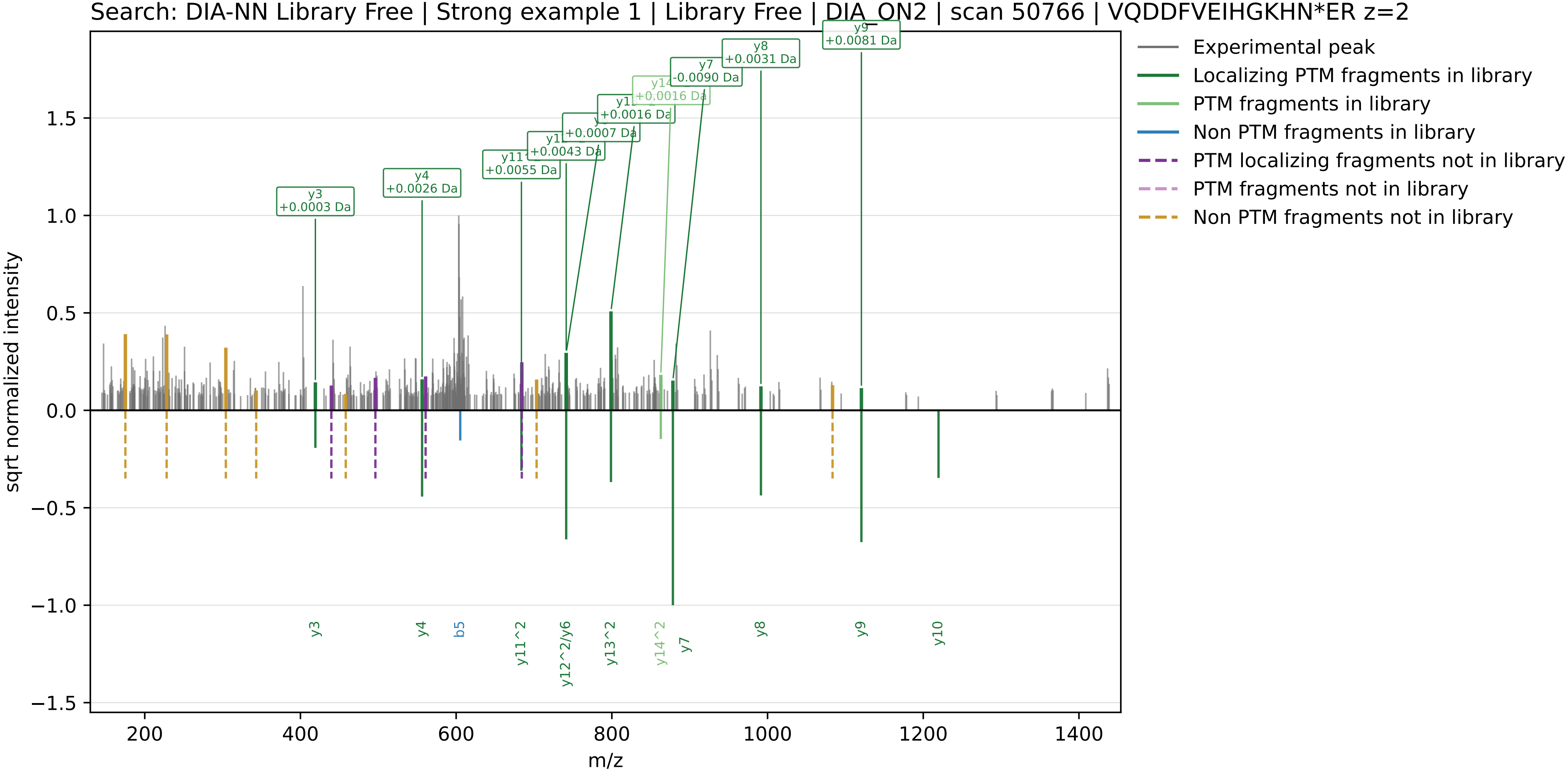

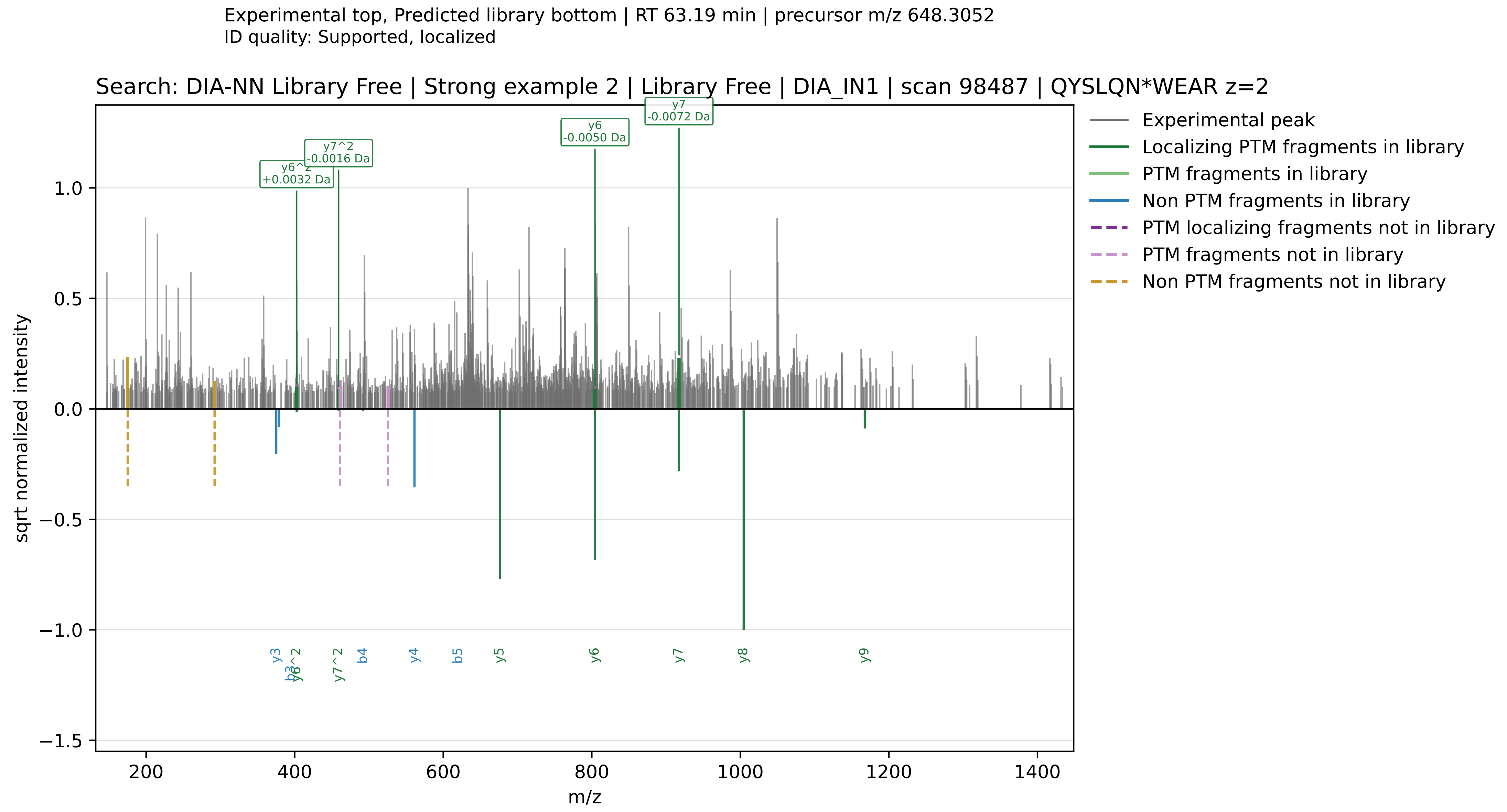

Experimental top, Predicted library bottom | RT 63.19 min | precursor m/z 648.3052
ID quality: Supported, localized
Search: DIA-NN Library Free | Strong example 2 | Library Free | DIA_IN1 | scan 98487 | QYSLQN*WEAR z=2
y7
-0.0072 Da
y6
-0.0050 Da
y7^2
-0.0016 Da
y6^2
+0.0032 Da
Experimental peak
Localizing PTM fragments in library
PTM fragments in library
Non PTM fragments in library
PTM localizing fragments not in library
PTM fragments not in library
Non PTM fragments not in library
sqrt normalized intensity
y3
b3
y6^2
y7^2
b4
y4
b5
y5
y6
y7
y8
y9
m/z

Experimental top, Predicted library bottom | RT 69.83 min | precursor m/z 743.6754
ID quality: Supported, localized

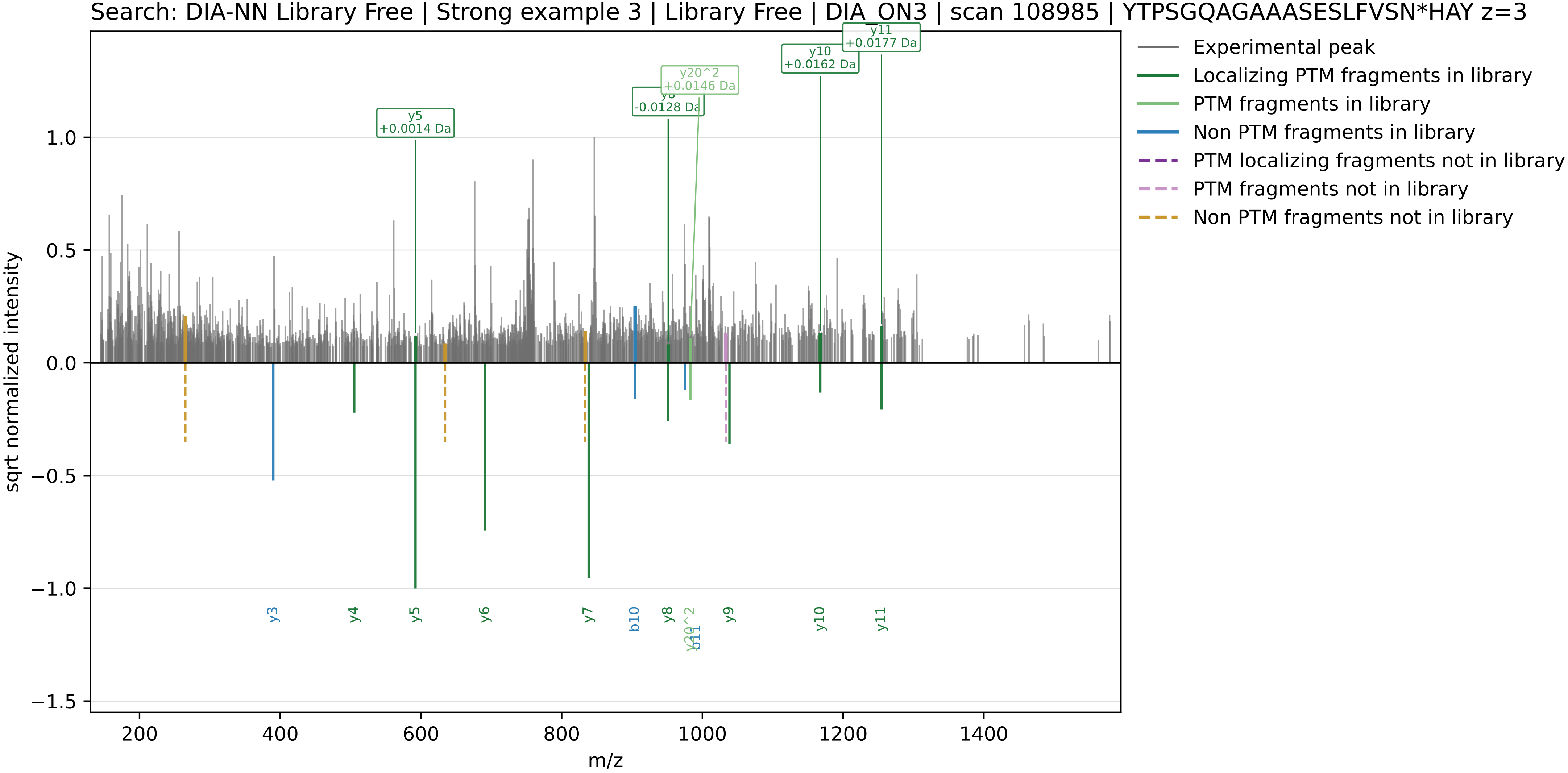

Experimental top, Predicted library bottom | RT 89.38 min | precursor m/z 706.0923
ID quality: Supported, localized

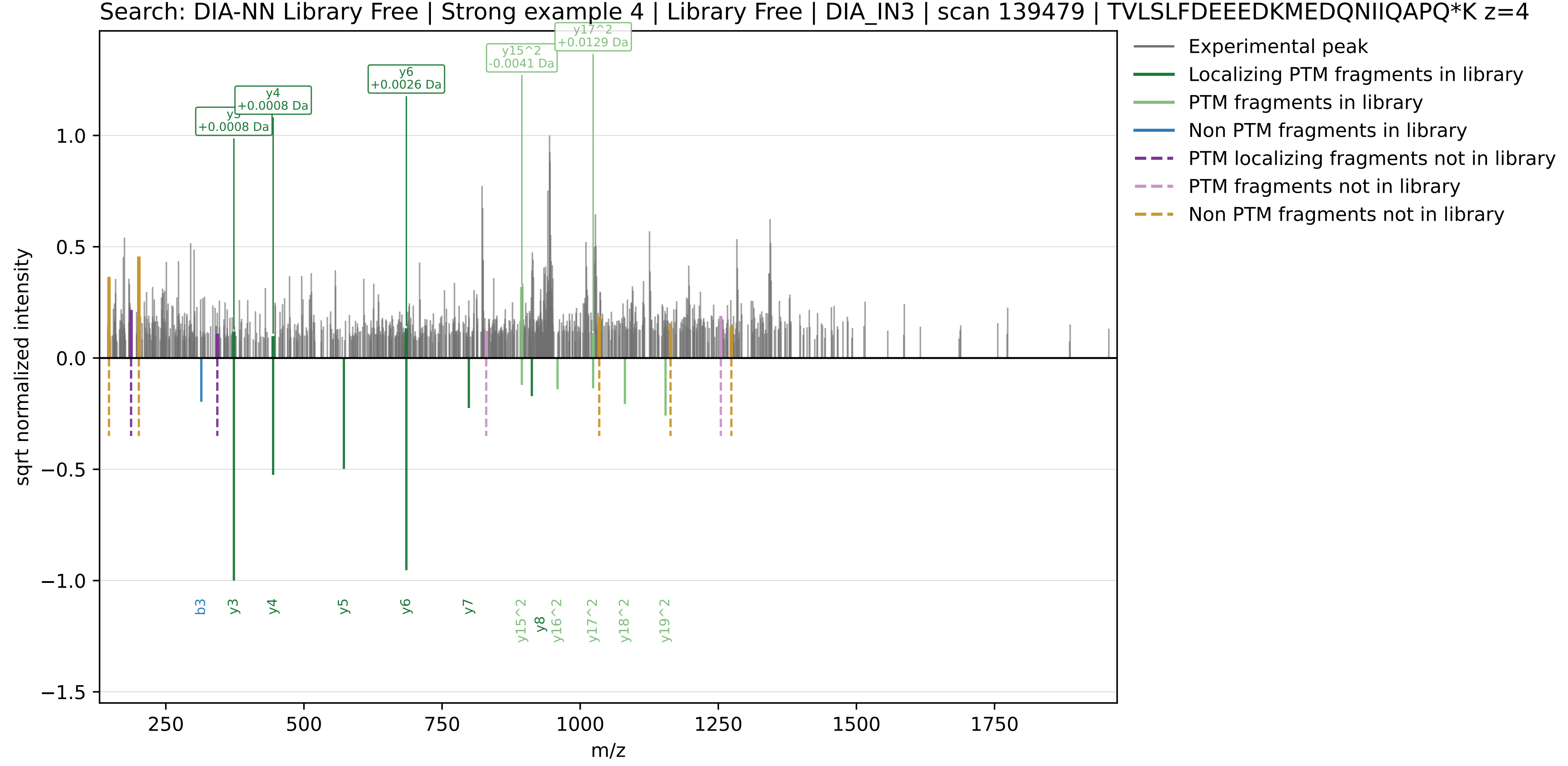

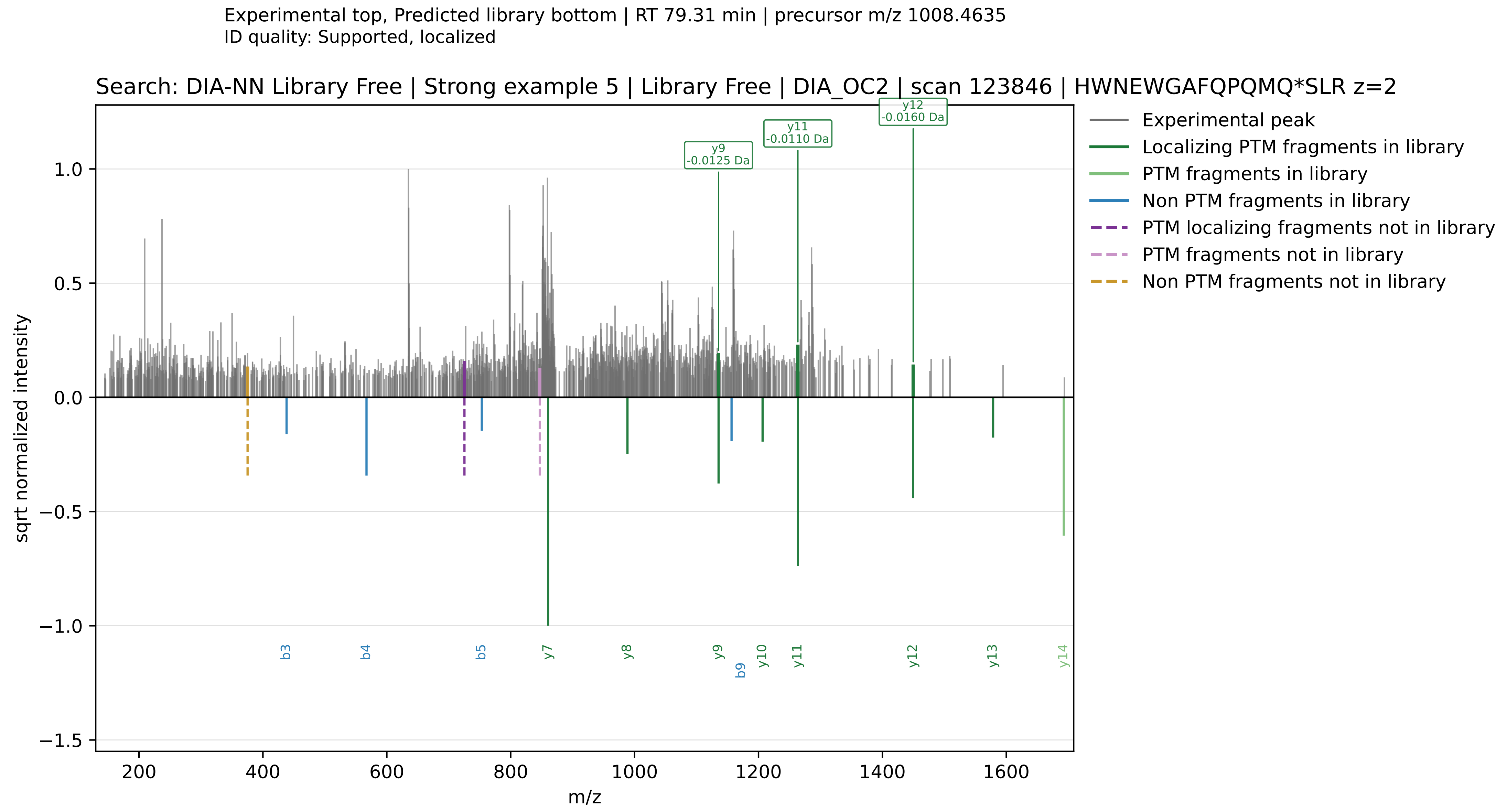

Experimental top, Predicted library bottom | RT 79.31 min | precursor m/z 1008.4635
ID quality: Supported, localized
Search: DIA-NN Library Free | Strong example 5 | Library Free | DIA_OC2 | scan 123846 | HWNEWGAFQPQMQ*SLR z=2
y9 -0.0125 Da
y11 -0.0110 Da
y12 -0.0160 Da
Experimental peak
Localizing PTM fragments in library
PTM fragments in library
Non PTM fragments in library
PTM localizing fragments not in library
PTM fragments not in library
Non PTM fragments not in library
sqrt normalized intensity
m/z
b3
b4
b5
y7
y8
y9
b9
y10
y11
y12
y13
y14

Experimental top, Predicted library bottom | RT 53.51 min | precursor m/z 590.7777
ID quality: Supported, isotopic-confounded

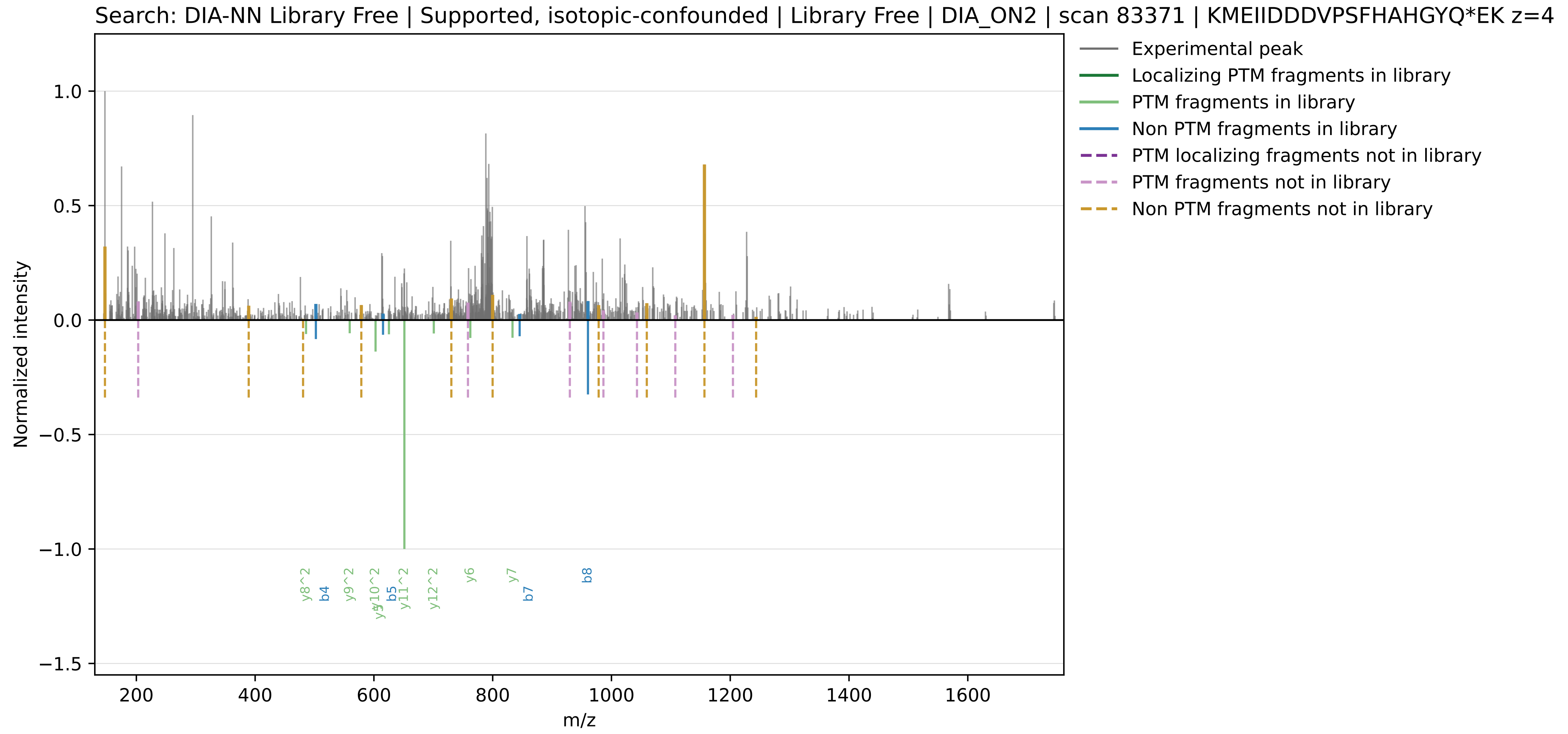

Experimental top, Predicted library bottom | RT 59.36 min | precursor m/z 785.0819
ID quality: No supporting fragment evidence

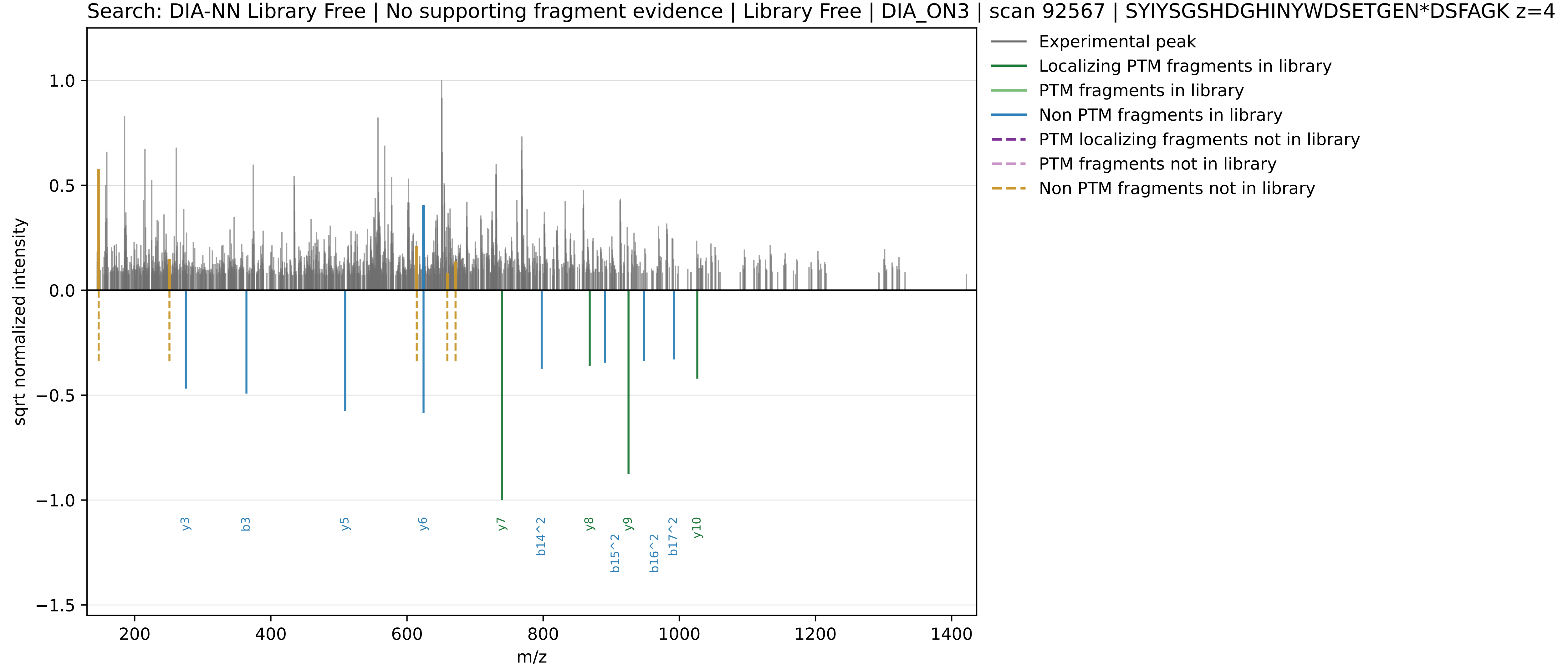

DIA-NN MBR

Library cache: diann_mbr_predicted_library.parquet
Selected spectra: 7

Contents:
- 5 strong supported examples
- 1 supported, isotopic-confounded example
- 1 unsupported example

Selection emphasis:
- strong examples require Supported, localized calls with at least one observed localizing deamidation fragment that is explicitly predicted in the method-specific library
- isotopic-confounded examples retain supporting evidence but have stronger competing isotope-like evidence overall
- unsupported examples show no retained deamidation-supporting fragments in the exported spectrum

Experimental top, Predicted library bottom | RT 33.20 min | precursor m/z 912.4380
ID quality: Supported, localized

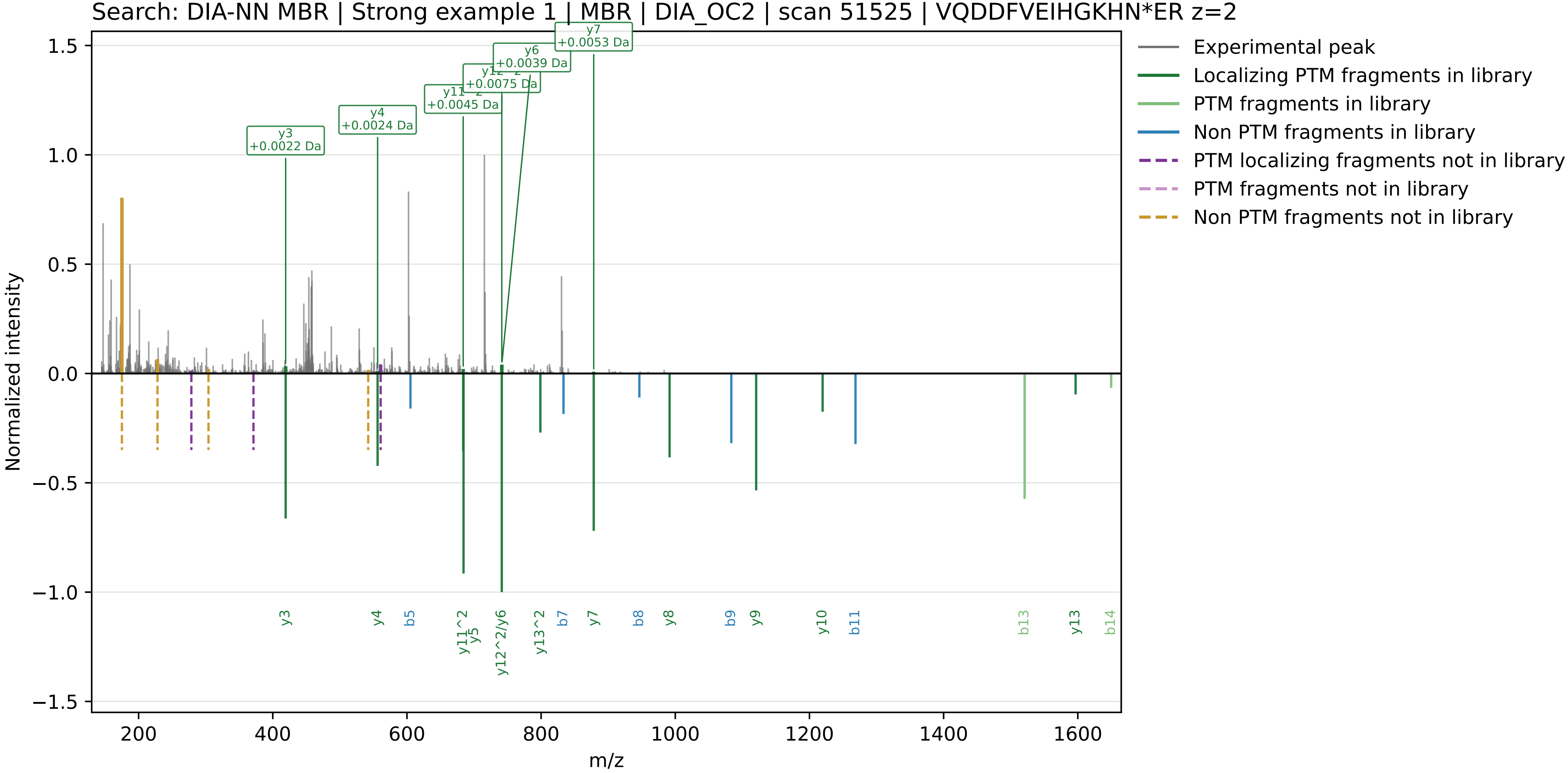

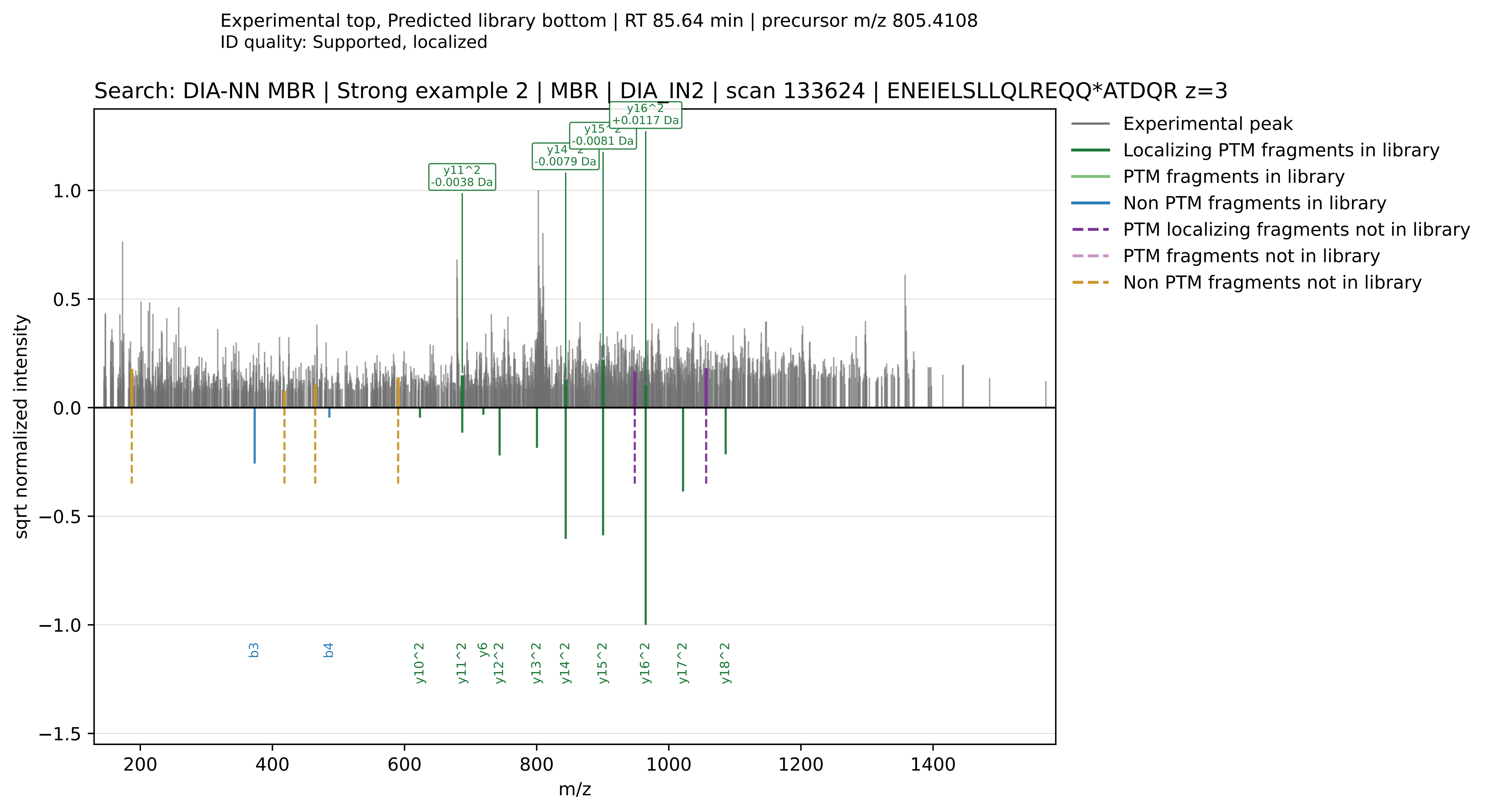

Experimental top, Predicted library bottom | RT 85.64 min | precursor m/z 805.4108
ID quality: Supported, localized
Search: DIA-NN MBR | Strong example 2 | MBR | DIA_IN2 | scan 133624 | ENEIELSLLQLREQQ*ATDQR z=3
y11^2 -0.0038 Da
y14^2 -0.0079 Da
y15^2 -0.0081 Da
y16^2 +0.0117 Da
Experimental peak
Localizing PTM fragments in library
PTM fragments in library
Non PTM fragments in library
PTM localizing fragments not in library
PTM fragments not in library
Non PTM fragments not in library
sqrt normalized intensity
m/z
b3
b4
y10^2
y11^2
y6
y12^2
y13^2
y14^2
y15^2
y16^2
y17^2
y18^2

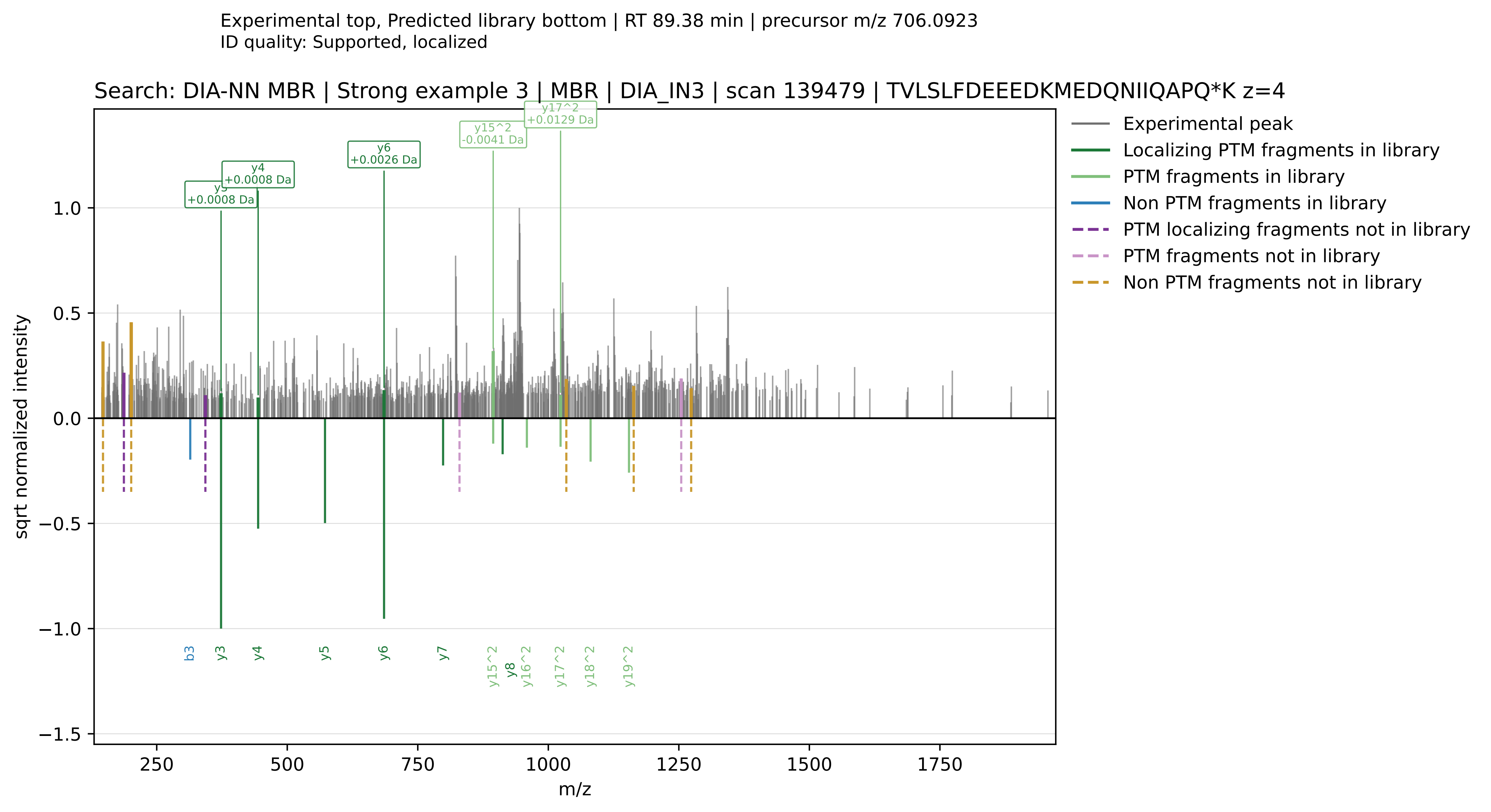
Experimental top, Predicted library bottom | RT 89.38 min | precursor m/z 706.0923
ID quality: Supported, localized
Search: DIA-NN MBR | Strong example 3 | MBR | DIA_IN3 | scan 139479 | TVLSLFDEEEDKMEDQNIIQAPQ*K z=4
Experimental peak
Localizing PTM fragments in library
PTM fragments in library
Non PTM fragments in library
PTM localizing fragments not in library
PTM fragments not in library
Non PTM fragments not in library
y3 +0.0008 Da
y4 +0.0008 Da
y6 +0.0026 Da
y15^2 -0.0041 Da
y17^2 +0.0129 Da
b3
y3
y4
y5
y6
y7
y15^2
y8
y16^2
y17^2
y18^2
y19^2
sqrt normalized intensity
m/z

Experimental top, Predicted library bottom | RT 65.27 min | precursor m/z 587.9466
ID quality: Supported, localized

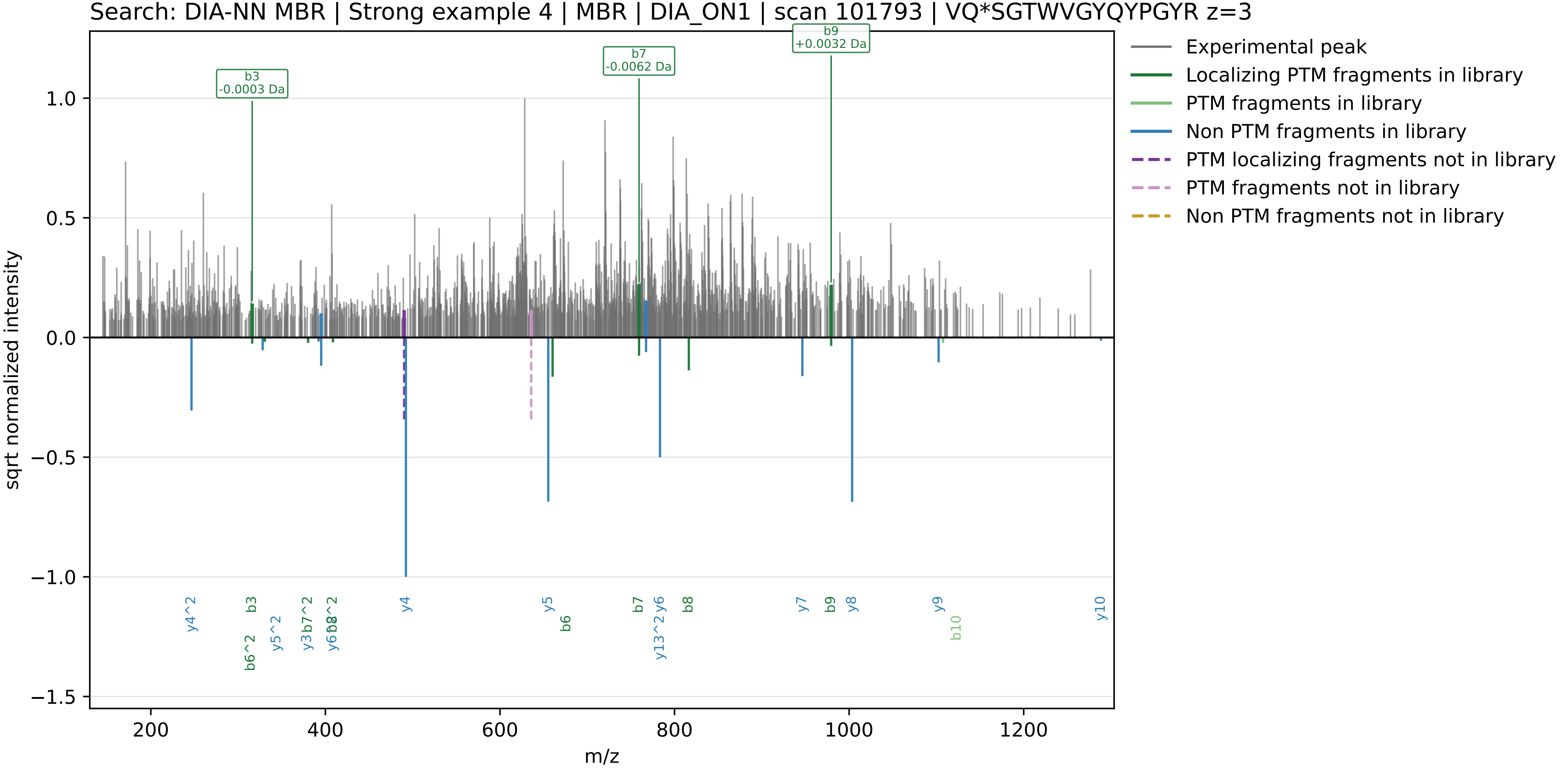

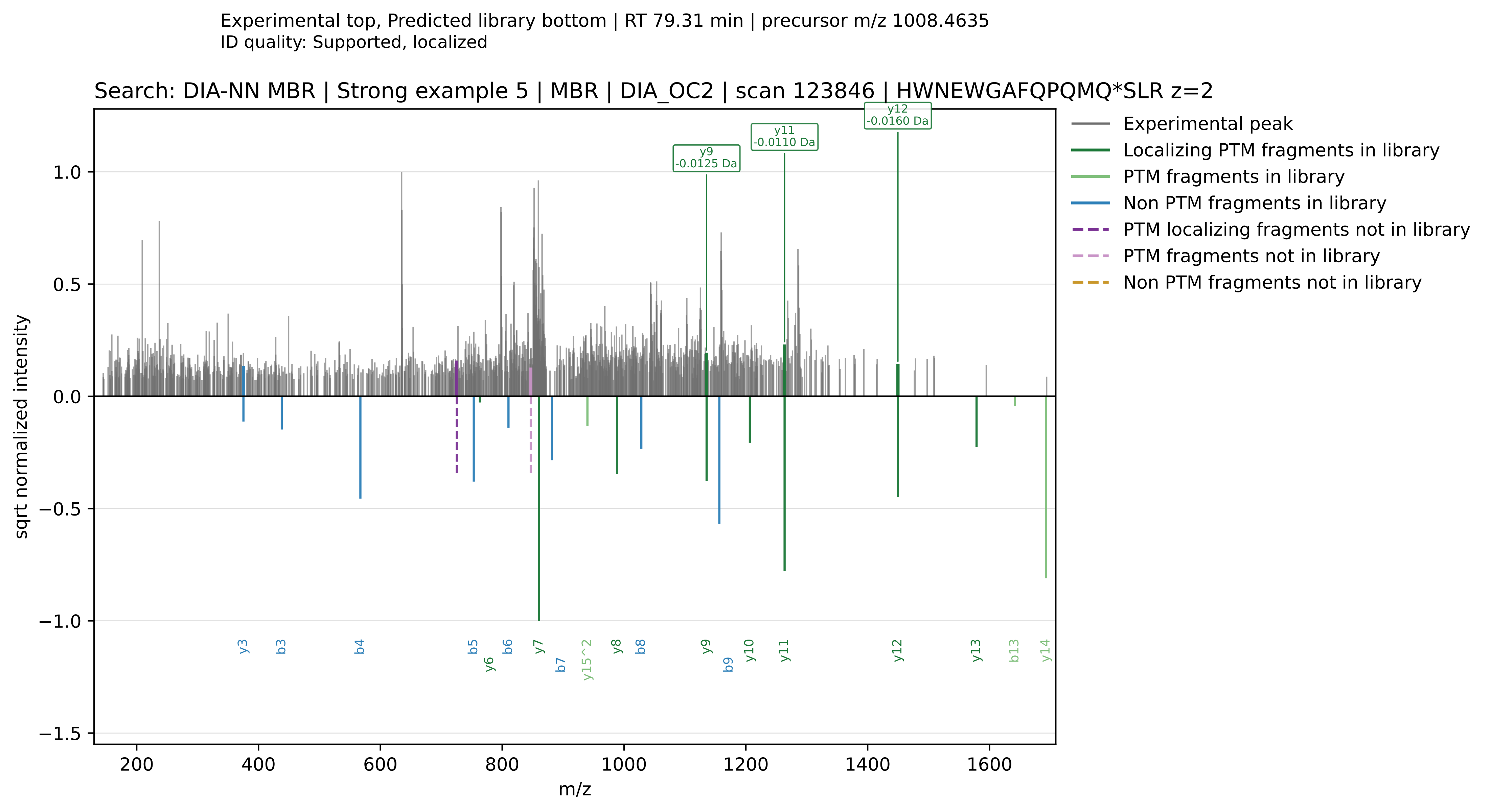
Experimental top, Predicted library bottom | RT 79.31 min | precursor m/z 1008.4635
ID quality: Supported, localized
Search: DIA-NN MBR | Strong example 5 | MBR | DIA_OC2 | scan 123846 | HWNEWGAFQPQMQ*SLR z=2
y9 -0.0125 Da
y11 -0.0110 Da
y12 -0.0160 Da
Experimental peak
Localizing PTM fragments in library
PTM fragments in library
Non PTM fragments in library
PTM localizing fragments not in library
PTM fragments not in library
Non PTM fragments not in library
sqrt normalized intensity
m/z
y3
b3
b4
b5
y6
b6
y7
b7
y15^2
y8
b8
y9
b9
y10
y11
y12
y13
b13
y14

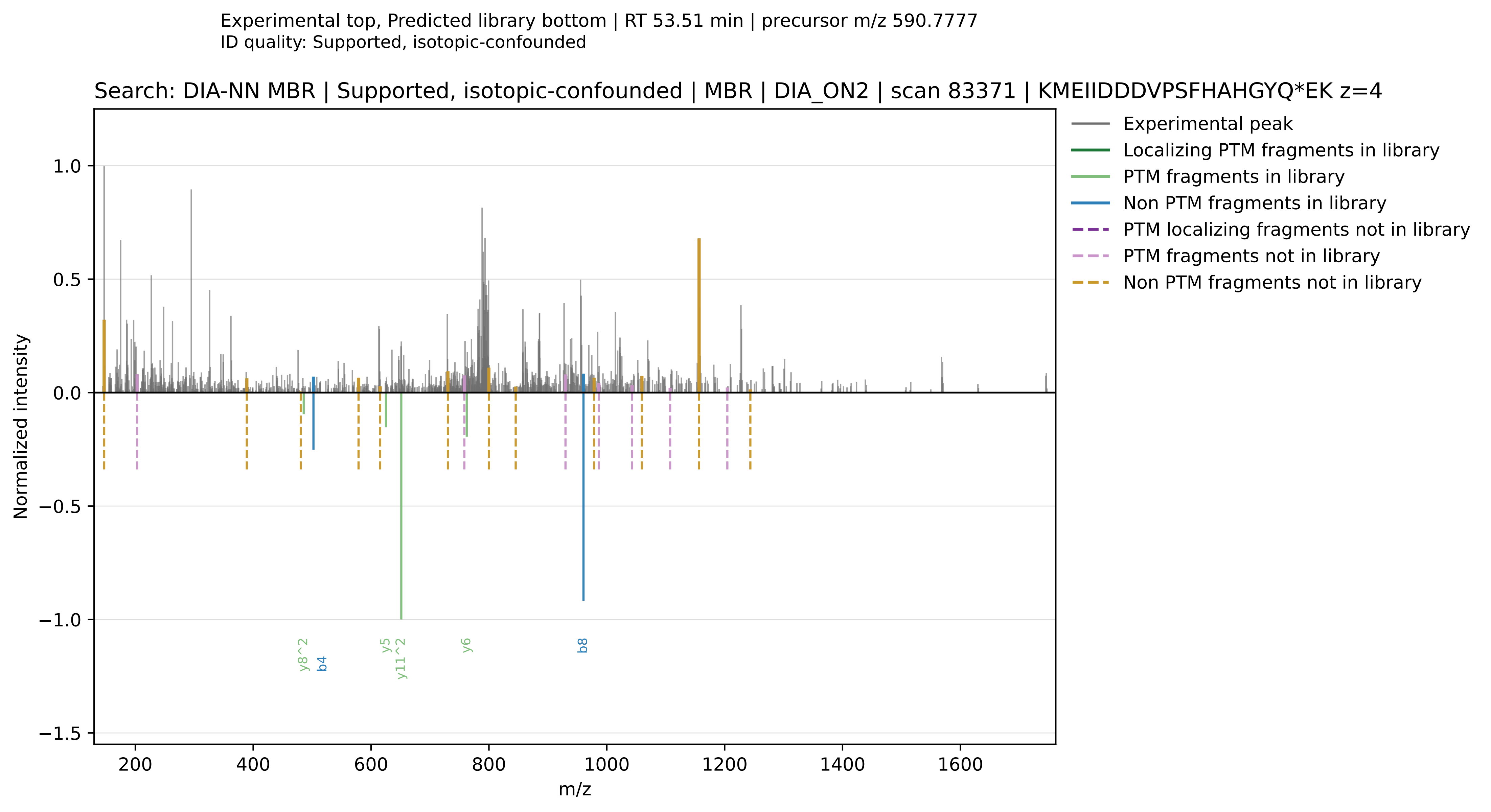

Experimental top, Predicted library bottom | RT 53.51 min | precursor m/z 590.7777
ID quality: Supported, isotopic-confounded
Search: DIA-NN MBR | Supported, isotopic-confounded | MBR | DIA_ON2 | scan 83371 | KMEIIDDDVPSFHAHGYQ*EK z=4
Experimental peak
Localizing PTM fragments in library
PTM fragments in library
Non PTM fragments in library
PTM localizing fragments not in library
PTM fragments not in library
Non PTM fragments not in library
Normalized intensity
m/z
y8^2
b4
y5
y11^2
y6
b8

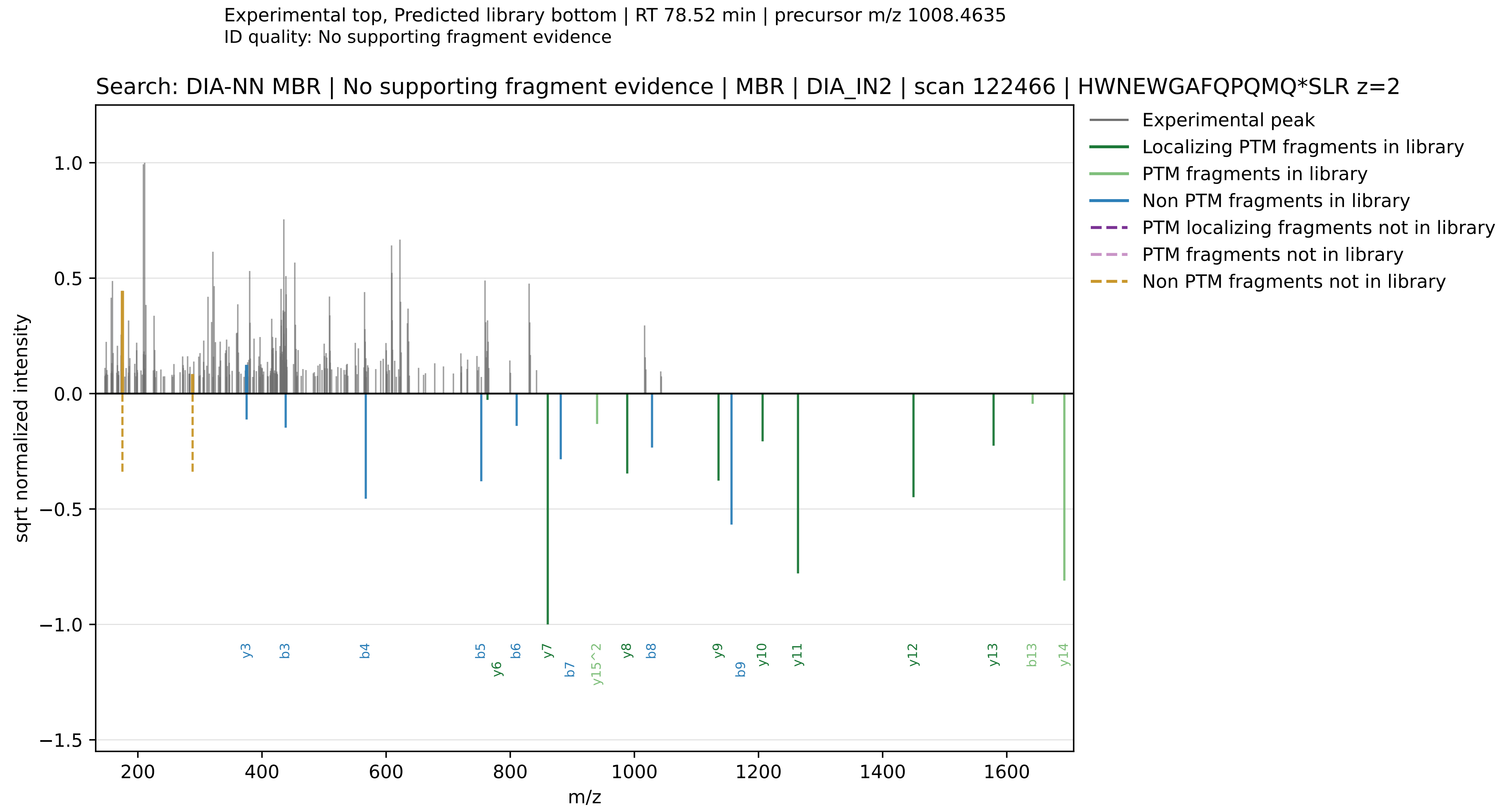

Experimental top, Predicted library bottom | RT 78.52 min | precursor m/z 1008.4635
ID quality: No supporting fragment evidence
Search: DIA-NN MBR | No supporting fragment evidence | MBR | DIA_IN2 | scan 122466 | HWNEWGAFQPQMQ*SLR z=2
Experimental peak
Localizing PTM fragments in library
PTM fragments in library
Non PTM fragments in library
PTM localizing fragments not in library
PTM fragments not in library
Non PTM fragments not in library
sqrt normalized intensity
m/z
1.0
0.5
0.0
−0.5
−1.0
−1.5
200
400
600
800
1000
1200
1400
1600
y3
b3
b4
b5
y6
b6
y7
b7
y15^2
y8
b8
y9
b9
y10
y11
y12
y13
b13
y14

Radiant Library Free

Library cache: shared_libfree_predicted_library.parquet
Selected spectra: 7

Contents:
- 5 strong supported examples
- 1 supported, isotopic-confounded example
- 1 unsupported example

Selection emphasis:
- strong examples require Supported, localized calls with at least one observed localizing deamidation fragment that is explicitly predicted in the method-specific library
- isotopic-confounded examples retain supporting evidence but have stronger competing isotope-like evidence overall
- unsupported examples show no retained deamidation-supporting fragments in the exported spectrum

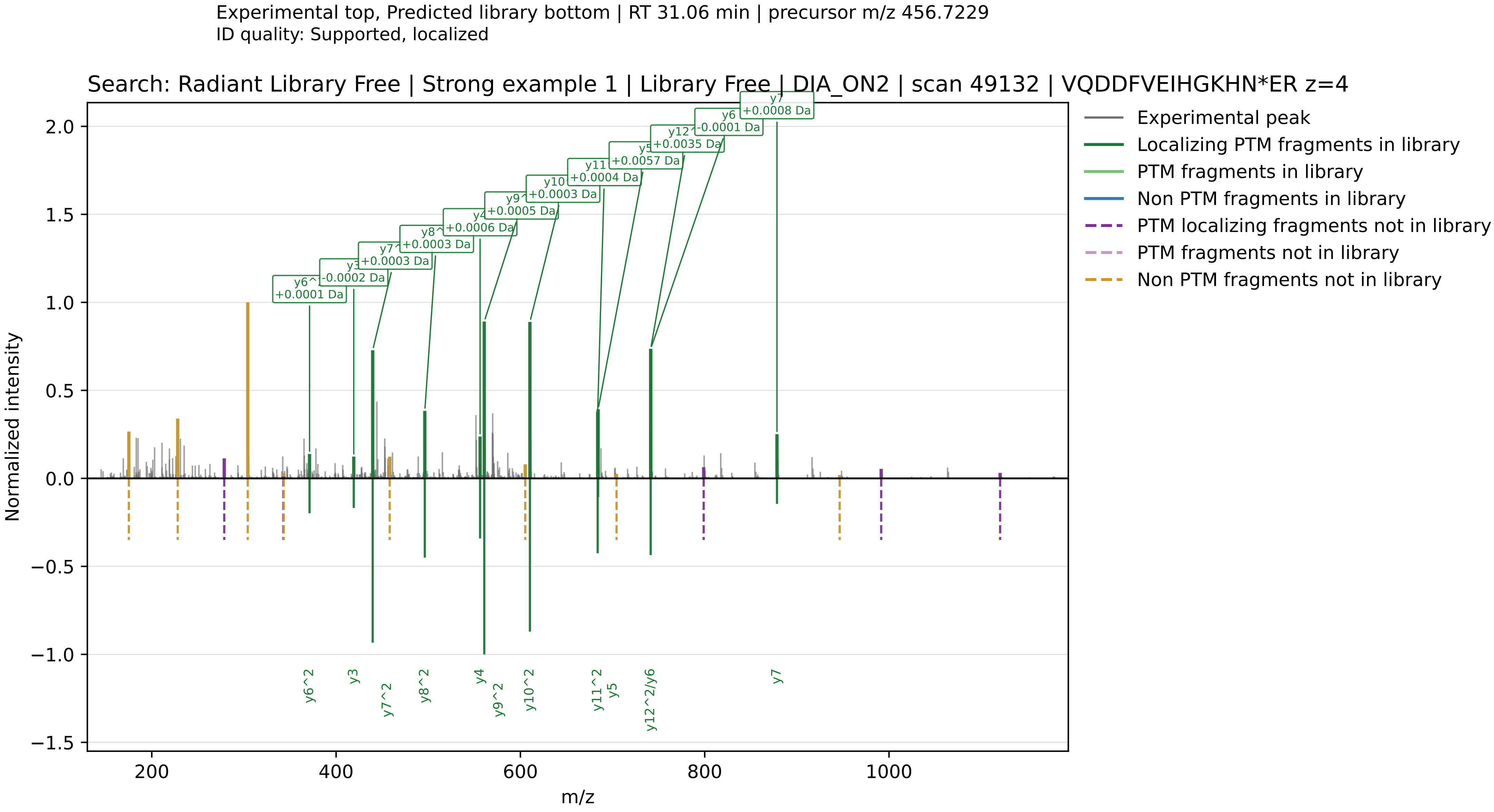

Experimental top, Predicted library bottom | RT 31.06 min | precursor m/z 456.7229
ID quality: Supported, localized
Search: Radiant Library Free | Strong example 1 | Library Free | DIA_ON2 | scan 49132 | VQDDFVEIHGKHN*ER z=4
Experimental peak
Localizing PTM fragments in library
PTM fragments in library
Non PTM fragments in library
PTM localizing fragments not in library
PTM fragments not in library
Non PTM fragments not in library
y6^2 +0.0001 Da
y3 -0.0002 Da
y7^2 +0.0003 Da
y8^2 +0.0003 Da
y4 +0.0006 Da
y9^2 +0.0005 Da
y10^2 +0.0003 Da
y11^2 +0.0004 Da
y5 +0.0057 Da
y12^2 +0.0035 Da
y6 -0.0001 Da
y7 +0.0008 Da
y6^2
y3
y7^2
y8^2
y4
y9^2
y10^2
y11^2
y5
y12^2/y6
y7
Normalized intensity
m/z

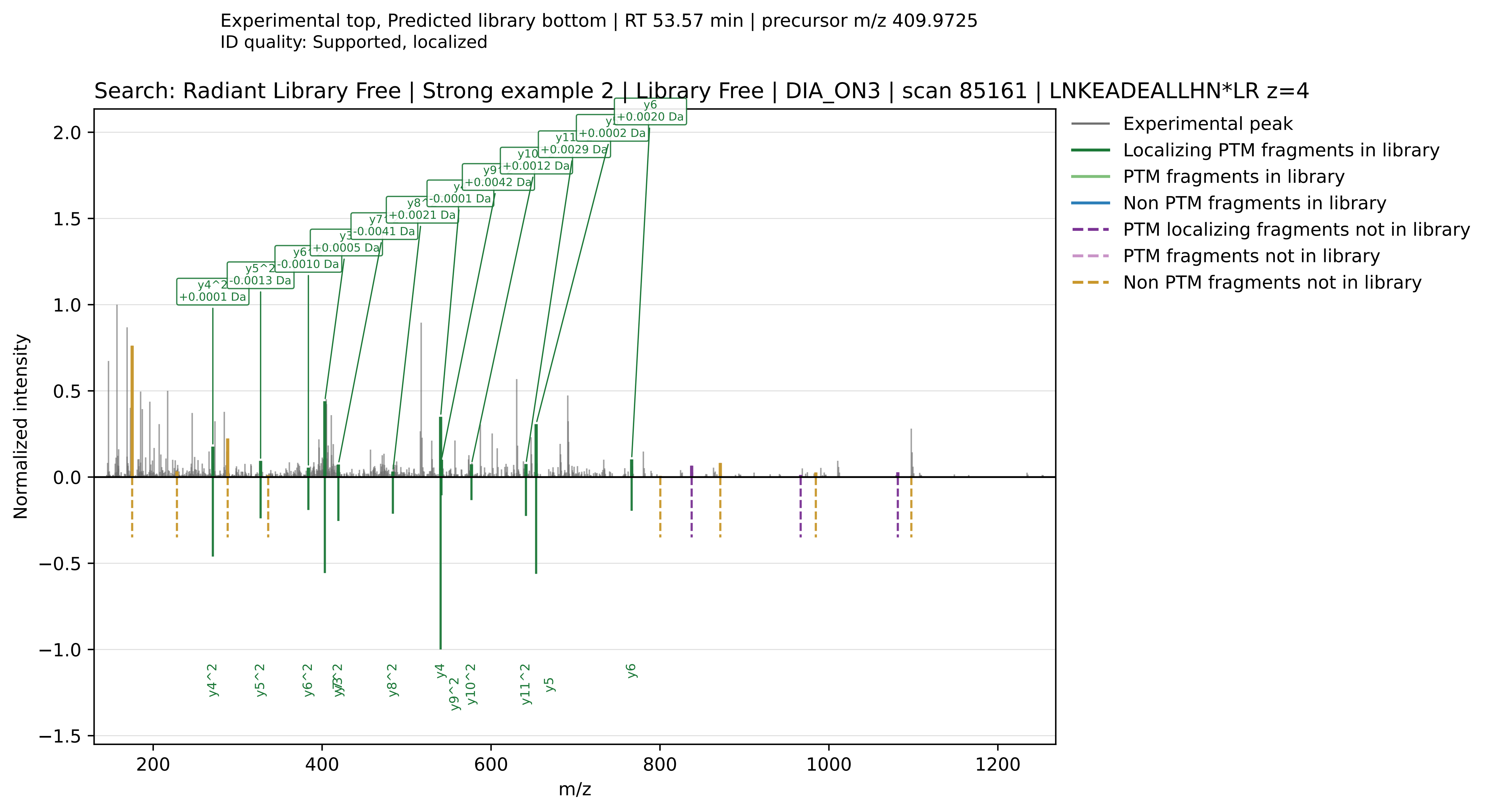

Experimental top, Predicted library bottom | RT 53.57 min | precursor m/z 409.9725
ID quality: Supported, localized
Search: Radiant Library Free | Strong example 2 | Library Free | DIA_ON3 | scan 85161 | LNKEADEALLHN*LR z=4
Experimental peak
Localizing PTM fragments in library
PTM fragments in library
Non PTM fragments in library
PTM localizing fragments not in library
PTM fragments not in library
Non PTM fragments not in library
y4^2 +0.0001 Da
y5^2 -0.0013 Da
y6 -0.0010 Da
y3 +0.0005 Da
y7 -0.0041 Da
y8 +0.0021 Da
-0.0001 Da
y9 +0.0042 Da
y10 +0.0012 Da
y11 +0.0029 Da
+0.0002 Da
y6 +0.0020 Da
Normalized intensity
m/z
y4^2
y5^2
y6^2
y7^2
y8^2
y4
y9^2
y10^2
y11^2
y5
y6

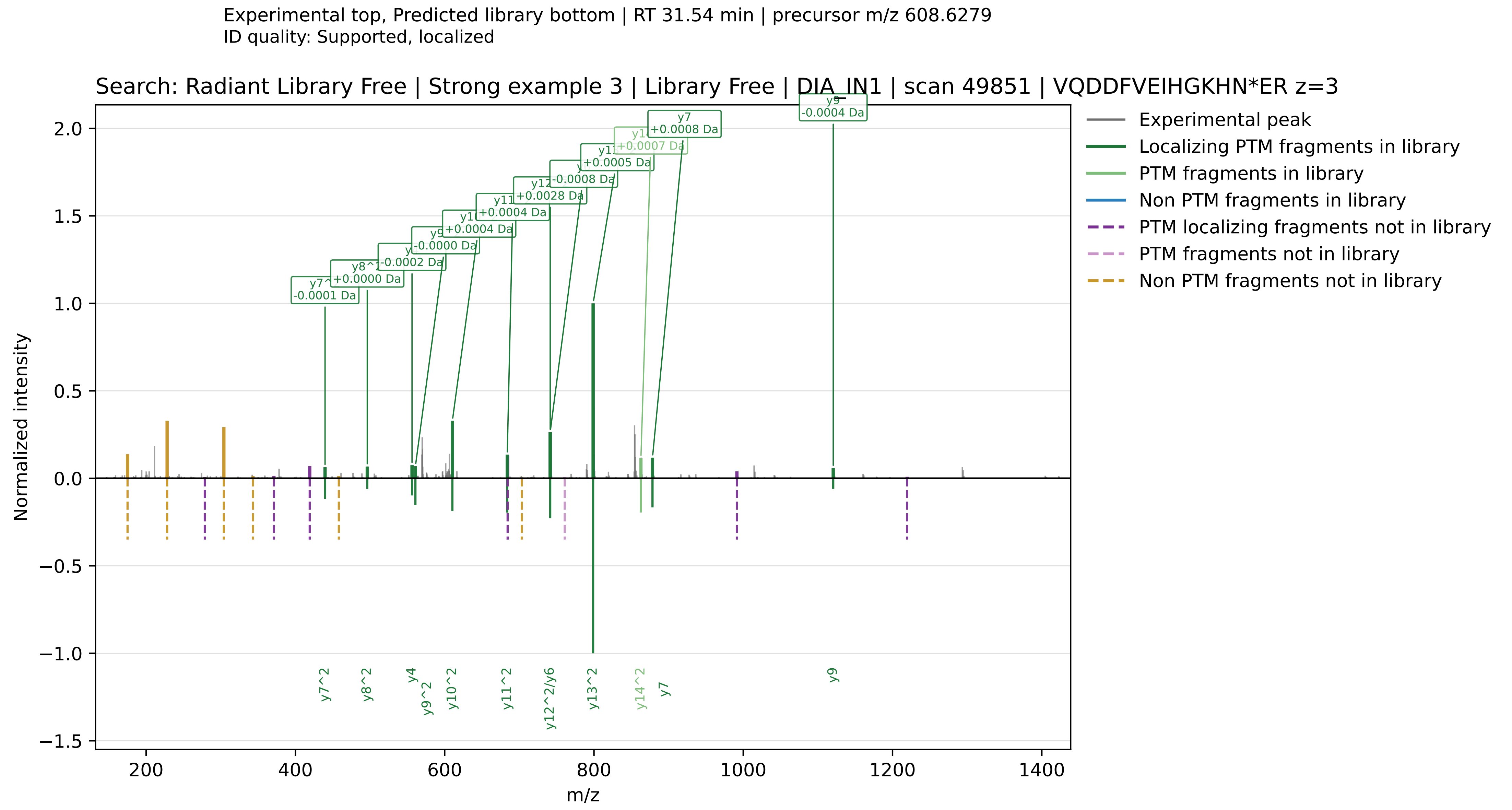

Experimental top, Predicted library bottom | RT 31.54 min | precursor m/z 608.6279
ID quality: Supported, localized
Search: Radiant Library Free | Strong example 3 | Library Free | DIA_IN1 | scan 49851 | VQDDFVEIHGKHN*ER z=3
Normalized intensity
m/z
Experimental peak
Localizing PTM fragments in library
PTM fragments in library
Non PTM fragments in library
PTM localizing fragments not in library
PTM fragments not in library
Non PTM fragments not in library
y9 -0.0004 Da
y7 +0.0008 Da
+0.0007 Da
+0.0005 Da
-0.0008 Da
+0.0028 Da
+0.0004 Da
+0.0004 Da
-0.0000 Da
-0.0002 Da
+0.0000 Da
-0.0001 Da
y7^2
y8^2
y4
y9^2
y10^2
y11^2
y12^2/y6
y13^2
y14^2
y7
y9

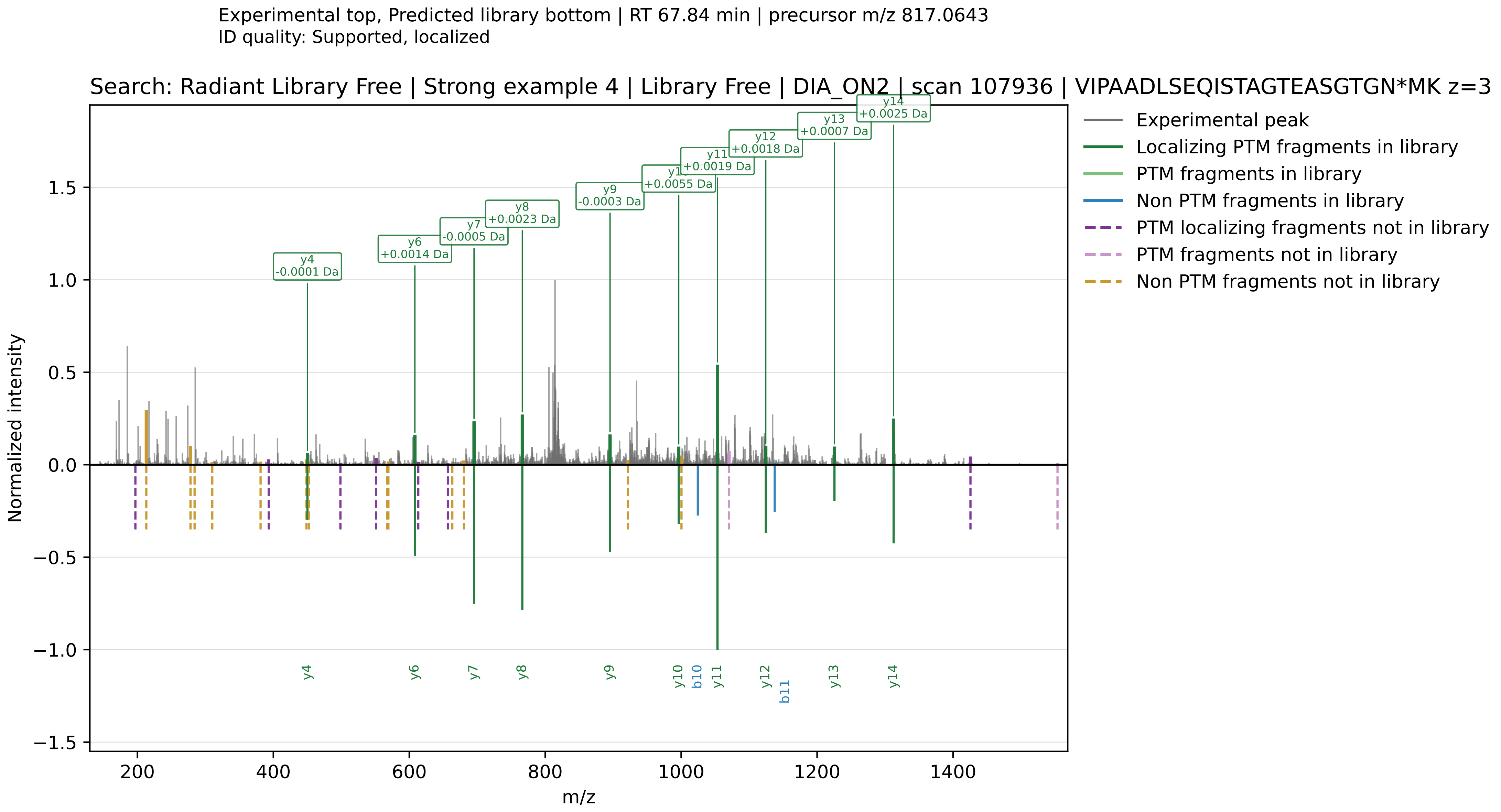

Experimental top, Predicted library bottom | RT 67.84 min | precursor m/z 817.0643
ID quality: Supported, localized
Search: Radiant Library Free | Strong example 4 | Library Free | DIA_ON2 | scan 107936 | VIPAADLSEQISTAGTEASGTGN*MK z=3
Experimental peak
Localizing PTM fragments in library
PTM fragments in library
Non PTM fragments in library
PTM localizing fragments not in library
PTM fragments not in library
Non PTM fragments not in library
y4 -0.0001 Da
y6 +0.0014 Da
y7 -0.0005 Da
y8 +0.0023 Da
y9 -0.0003 Da
+0.0055 Da
+0.0019 Da
y12 +0.0018 Da
y13 +0.0007 Da
y14 +0.0025 Da
y4
y6
y7
y8
y9
y10
b10
y11
y12
b11
y13
y14
Normalized intensity
m/z

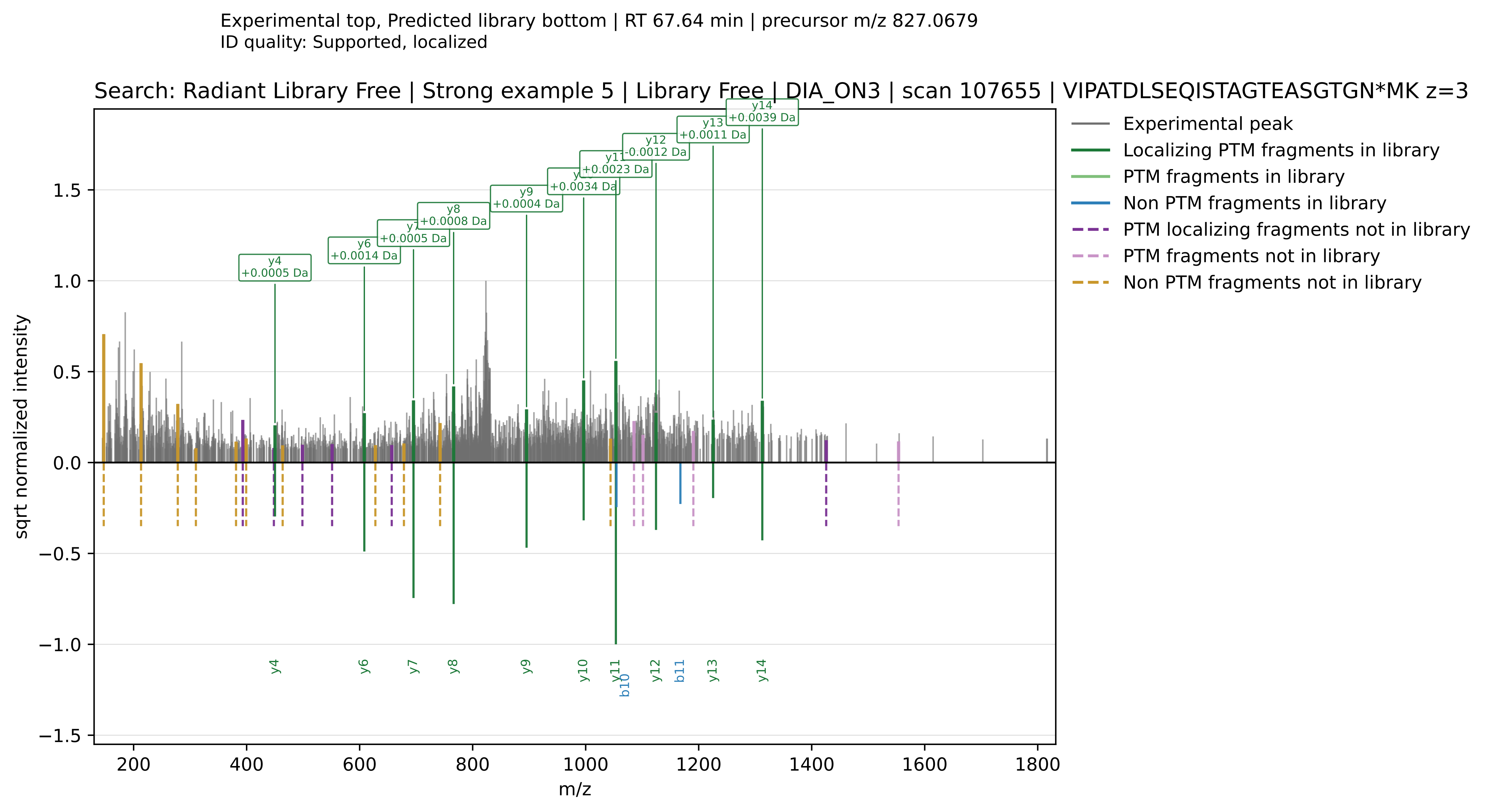
Experimental top, Predicted library bottom | RT 67.64 min | precursor m/z 827.0679
ID quality: Supported, localized
Search: Radiant Library Free | Strong example 5 | Library Free | DIA_ON3 | scan 107655 | VIPATDLSEQISTAGTEASGTGN*MK z=3
Experimental peak
Localizing PTM fragments in library
PTM fragments in library
Non PTM fragments in library
PTM localizing fragments not in library
PTM fragments not in library
Non PTM fragments not in library
y4 +0.0005 Da
y6 +0.0014 Da
+0.0005 Da
y8 +0.0008 Da
y9 +0.0004 Da
+0.0034 Da
+0.0023 Da
y12 -0.0012 Da
y13 +0.0011 Da
y14 +0.0039 Da
sqrt normalized intensity
m/z
y4
y6
y7
y8
y9
y10
y11
b10
y12
b11
y13
y14

Experimental top, Predicted library bottom | RT 68.06 min | precursor m/z 851.1599
ID quality: Supported, isotopic-confounded

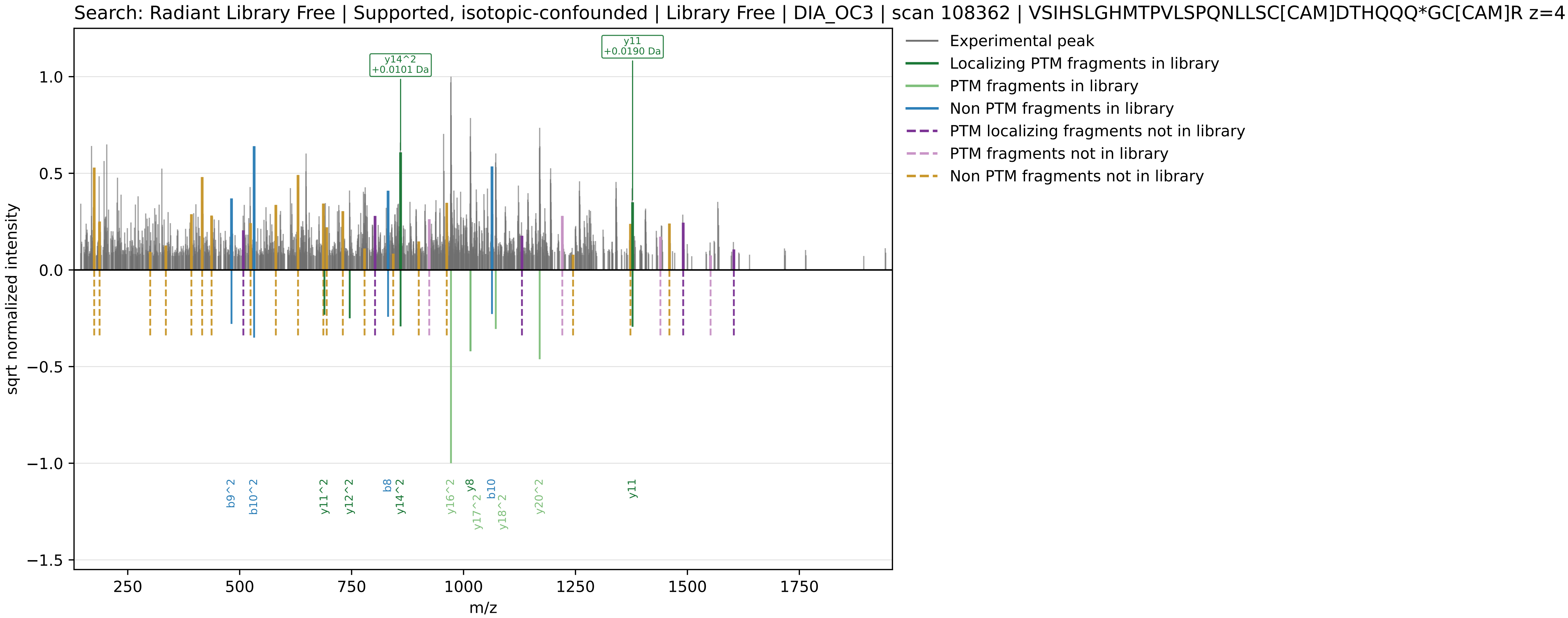

Experimental top, Predicted library bottom | RT 88.06 min | precursor m/z 971.1425
ID quality: No supporting fragment evidence

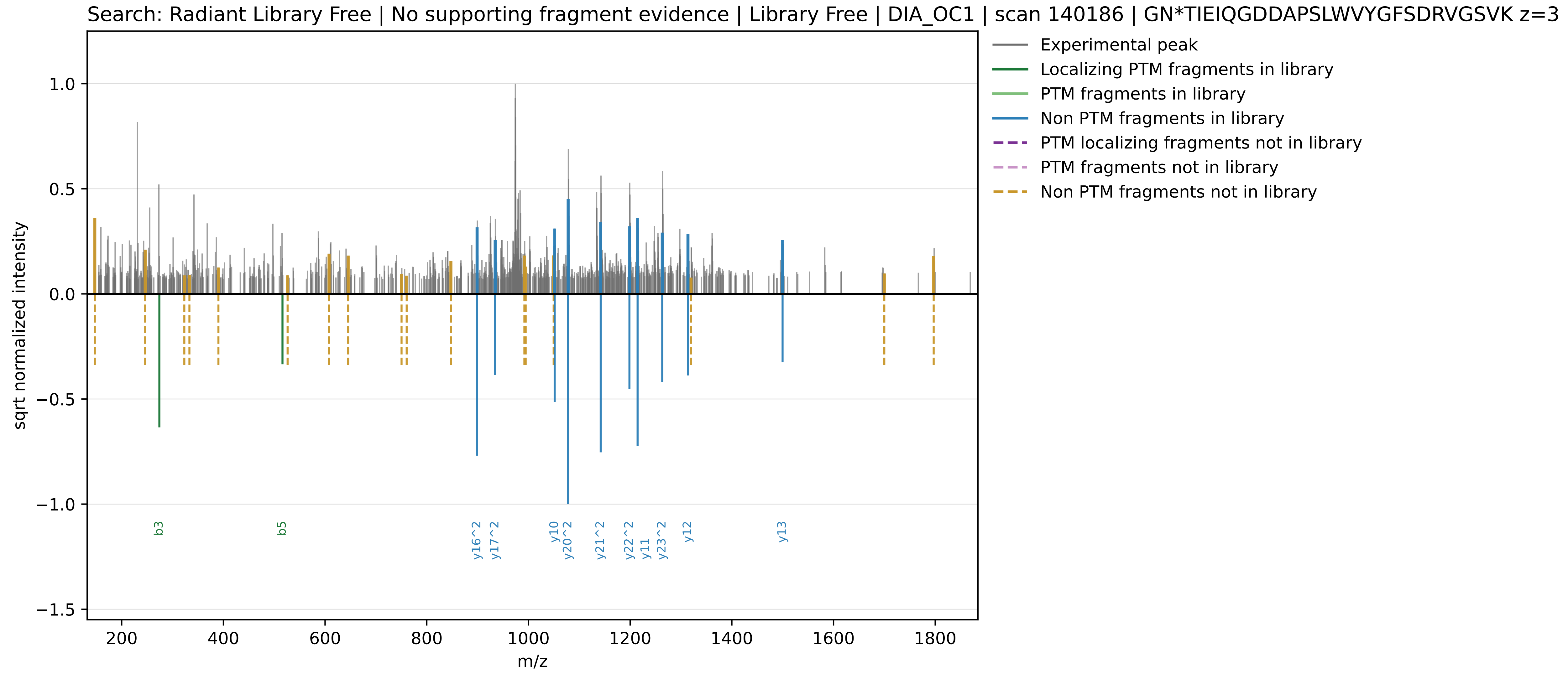

Radiant MBR

Library cache: radiant_mbr_predicted_library.parquet
Selected spectra: 7

Contents:
- 5 strong supported examples
- 1 supported, isotopic-confounded example
- 1 unsupported example

Selection emphasis:
- strong examples require Supported, localized calls with at least one observed localizing deamidation fragment that is explicitly predicted in the method-specific library
- isotopic-confounded examples retain supporting evidence but have stronger competing isotope-like evidence overall
- unsupported examples show no retained deamidation-supporting fragments in the exported spectrum

Experimental top, Radiant MBR library bottom | RT 31.87 min | precursor m/z 456.7229
ID quality: Supported, localized

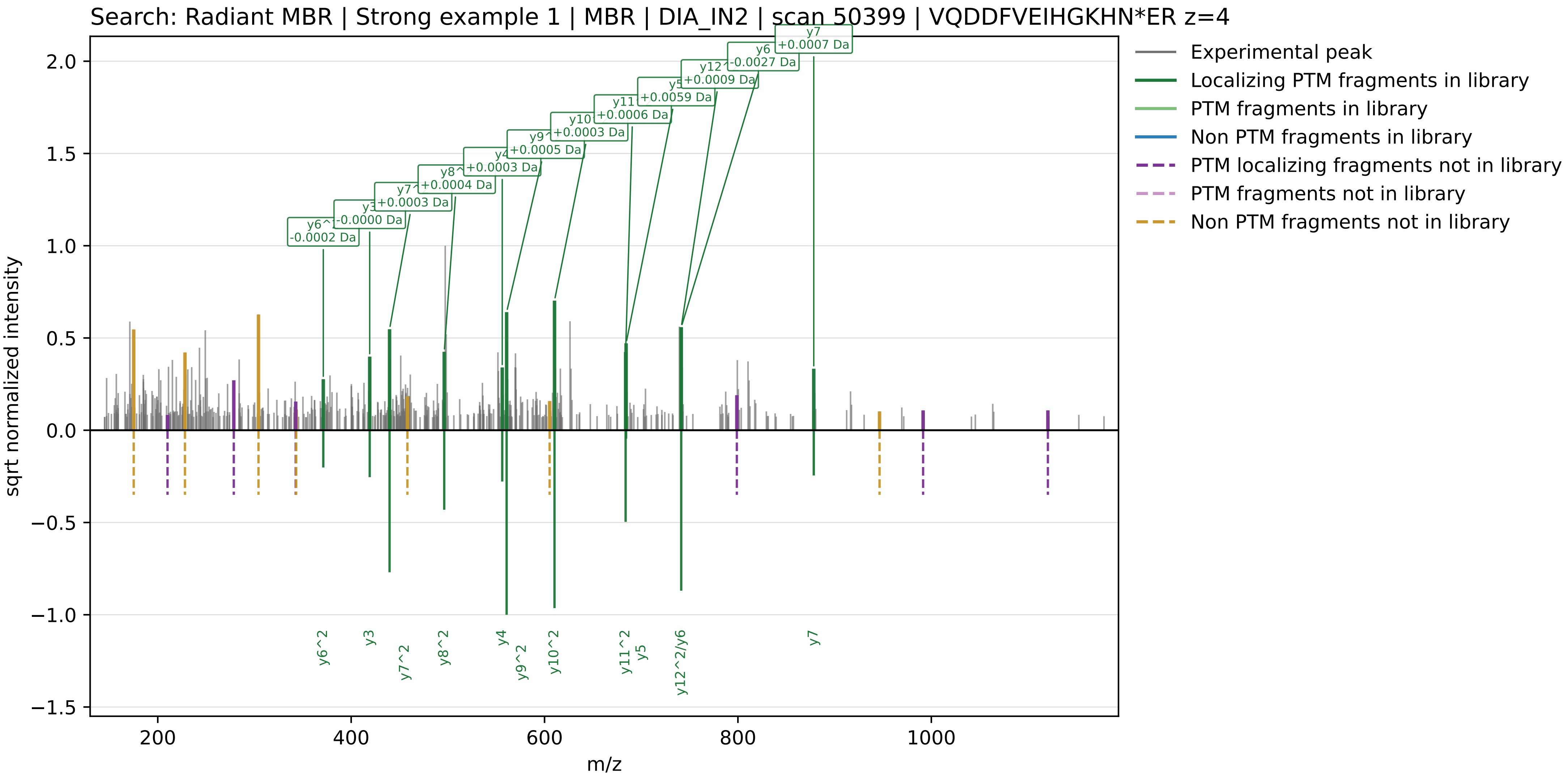

Experimental top, Radiant MBR library bottom | RT 31.54 min | precursor m/z 608.6279
ID quality: Supported, localized

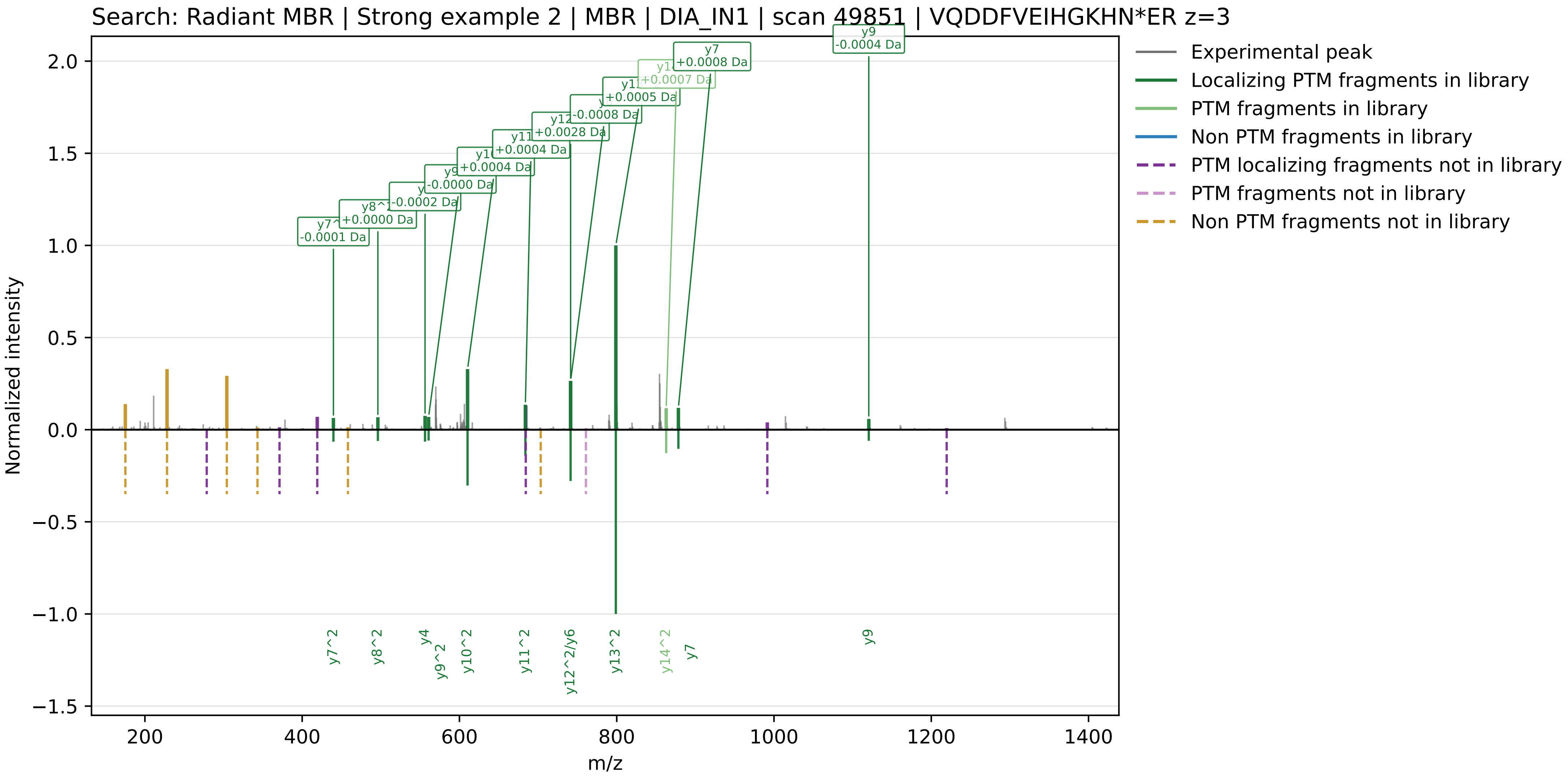

Experimental top, Radiant MBR library bottom | RT 64.81 min | precursor m/z 496.0086
ID quality: Supported, localized

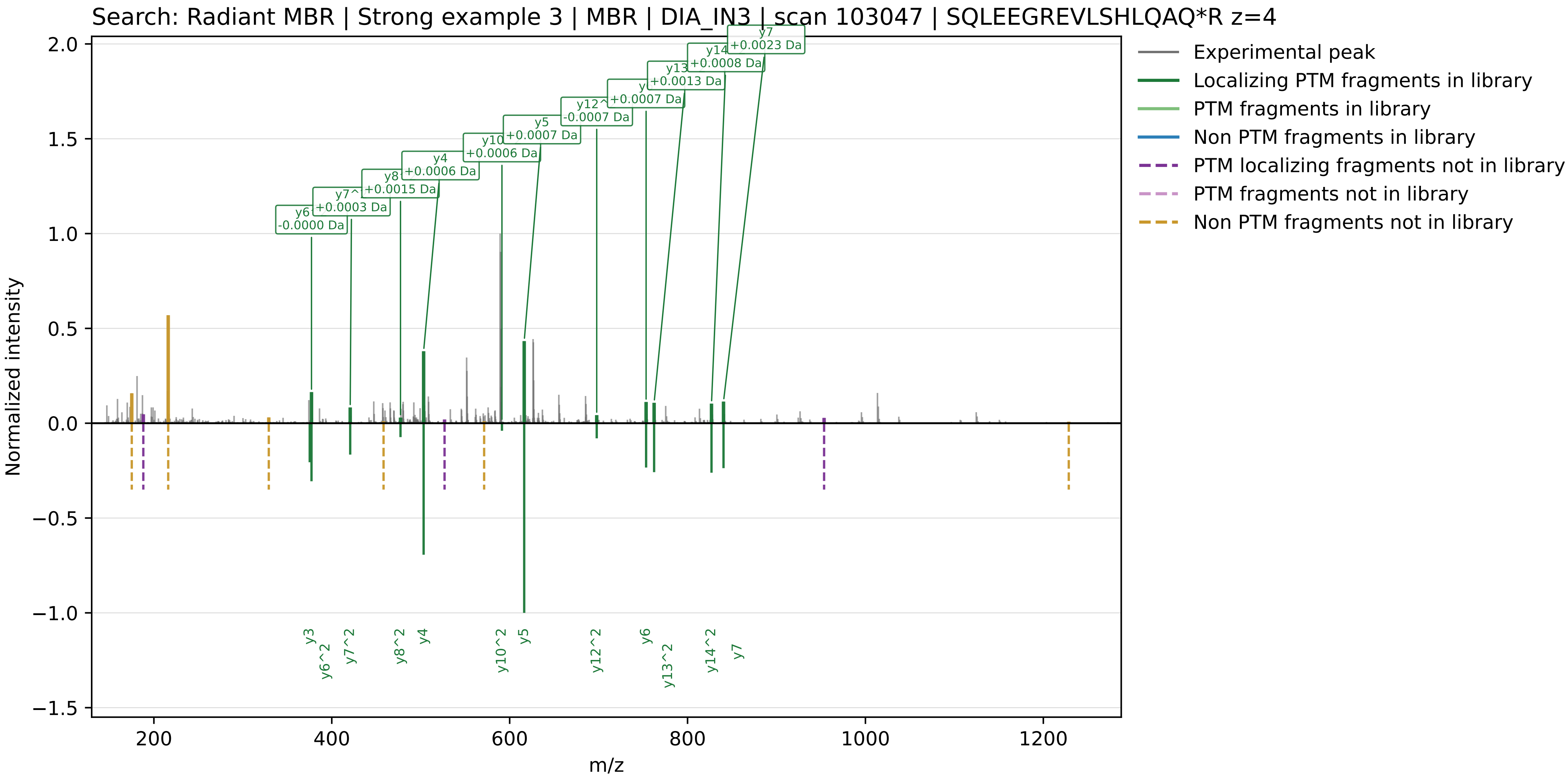

Experimental top, Radiant MBR library bottom | RT 67.64 min | precursor m/z 827.0680
ID quality: Supported, localized

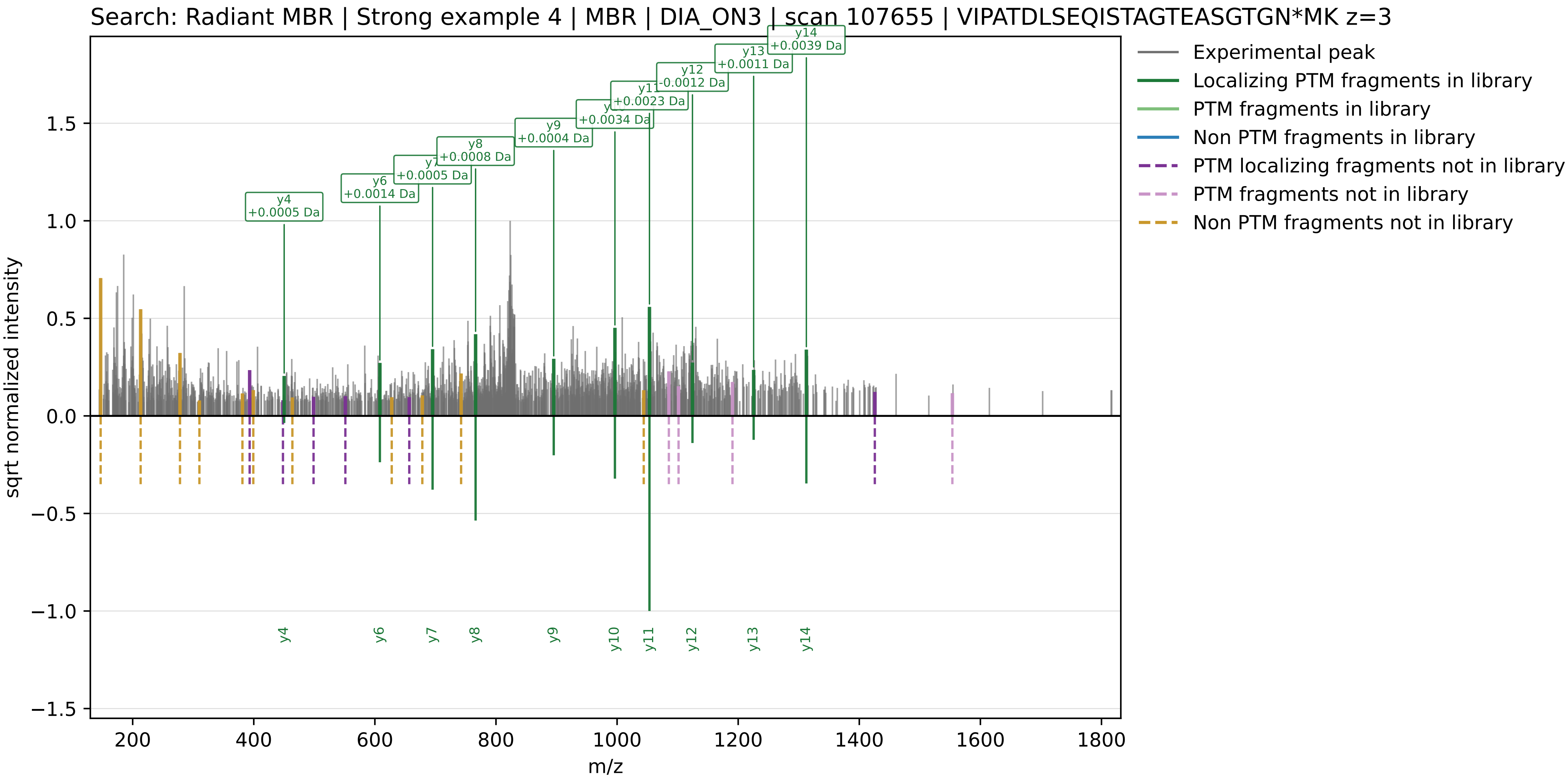

Experimental top, Radiant MBR library bottom | RT 58.68 min | precursor m/z 576.6484
ID quality: Supported, localized

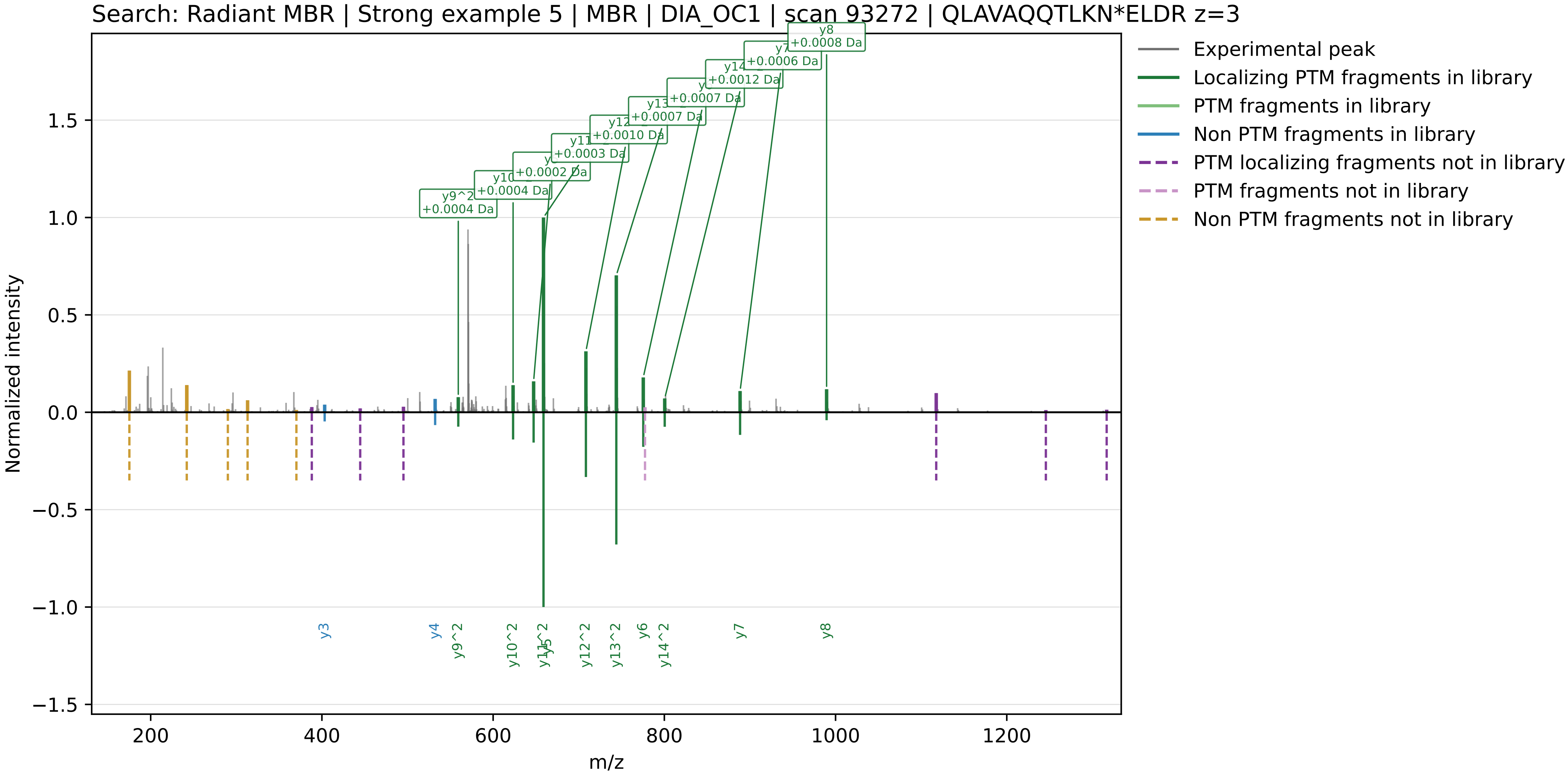

Experimental top, Radiant MBR library bottom | RT 85.12 min | precursor m/z 1025.8129
ID quality: Supported, isotopic-confounded

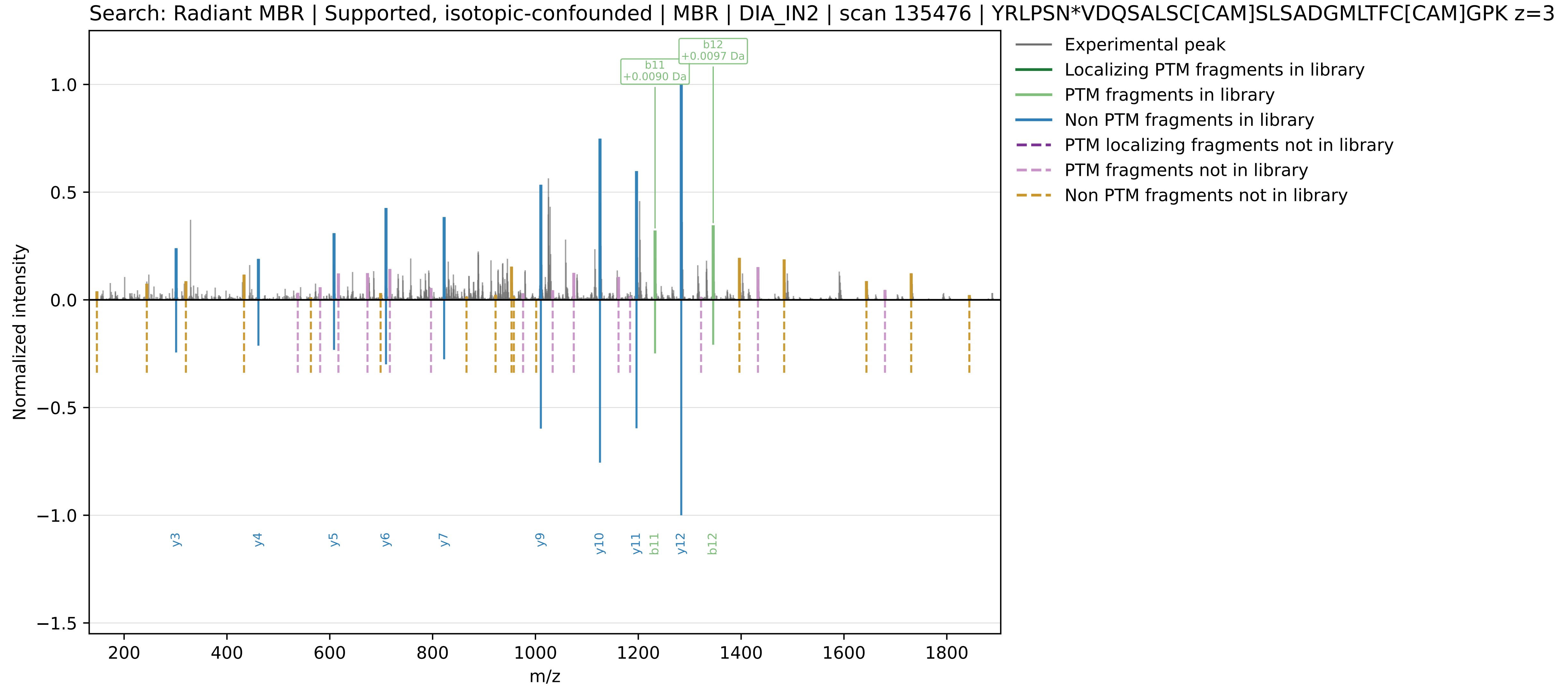

Experimental top, Radiant MBR library bottom | RT 88.06 min | precursor m/z 971.1425
ID quality: No supporting fragment evidence

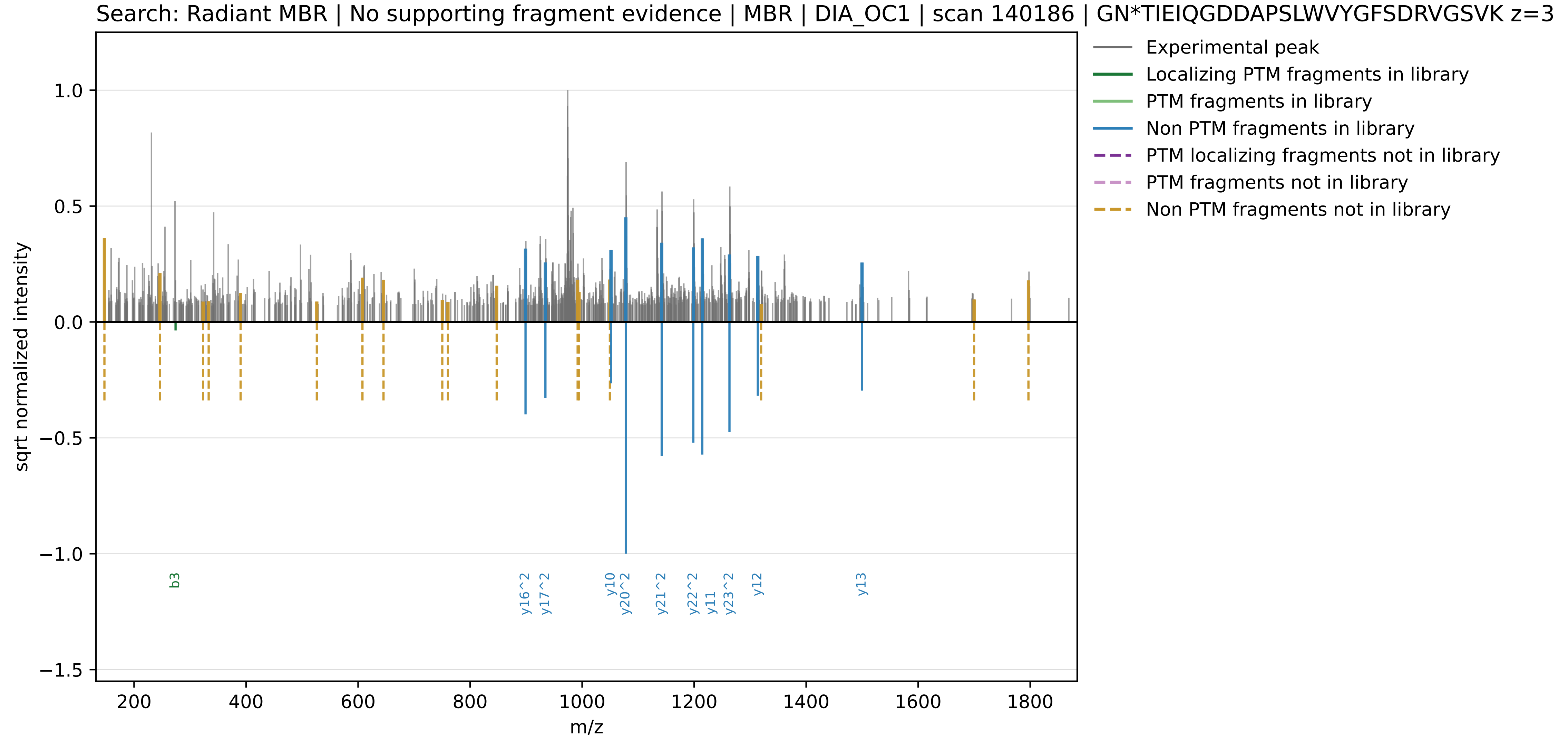
